# Quantifying uncertainty in the inference of generalized coalescents

**DOI:** 10.1101/150342

**Authors:** Timothy C. Wallstrom, Tanmoy Bhattacharya, Jon F. Wilkins

## Abstract

We develop inference methods for generalized coalescent models, such as the *Λ*- and *Ξ*-coalescents, which have recently been proposed for populations with broad offspring distributions, repeated selective sweeps, or strong selection. These are all populations that may not be adequately described by the usual Kingman coalescent. A roadblock to the application of such models has been the lack of effective tools for inferring an appropriate model, which stems from difficulties in evaluating the associated likelihoods. We overcome these difficulties by introducing estimators that are both computationally tractable and statistically efficient. We use these estimators to obtain point estimates and confidence intervals for the parameters of the coalescent models, and p-values for the hypothesis that the population is described by the Kingman coalescent. Our approach is based on the theory of unbiased estimating equations, which is more general than composite likelihood and may be applicable in other areas of statistical genetics. Our main focus is on inference from linked site-frequency spectra using parameterized families of *Λ*-coalescents. We show that useful inferences may be made from non-singleton data alone if singletons are suspect due to sequencing or data-cleaning errors, although the data requirements are greatly increased. We apply our method to mitochondrial sequence data from Gadus morhua, the Atlantic cod.

## 1. Introduction

Generalized coalescent models, such as the Λ- and Ξ-coalescents, have recently been developed as alternatives to the classical Kingman coalescent (Pitman, 1999, Sagitov, 1999, Schweinsberg, 2000, 2003). These models were initially designed for populations evolving neutrally with broad offspring distributions, in which the family size of an individual can be a significant fraction of the population (Eldon and Wakeley, 2006, Schweinsberg, 2003). They have also been used to model population bottlenecks and strong selection (Durrett and Schweinsberg, 2004, 2005, Eldon et al., 2015). The characteristic feature of the generalized coalescents, which distinguishes them from the ingman coalescent, is that more than two lineages can coalesce in a single event; in this case, we speak of a “multiple collision.” The shape of a generalized coalescent tree is often quite different from that of the Kingman coalescent tree.

One of the challenges in working with generalized coalescents is that they come in great variety, in contrast to the Kingman coalescent, which is essentially unique up to time reparameterization. The space of Λ-coalescents, for example, corresponds to the space of measures on the unit interval (Pitman, 1999), which is infinite dimensional. Despite the great variety of coalescents that are mathematically possible, it is reasonable to conjecture that only a limited subspace is biologically relevant. In previous work, simple parametric subspaces of Λ-coalescents have been introduced, based on biologically-motivated models of the offspring distribution (Eldon and Wakeley, 2006, Schweinsberg, 2003). These models interpolate smoothly between the classical Kingman coalescent and the star coalescent, as a shape parameter *φ* is varied over an interval. As an example, in Figure 1, we show sample trees as *φ* is varied in a particular model. Note that as *φ* increases the shape of the tree changes, so that an increasing fraction of the tree is found in the external branches. Note also that as *φ* increases, the prevalence of “multiple collisions”—coalescent events involving more than two lineages—increases as well.

**Figure 1:**
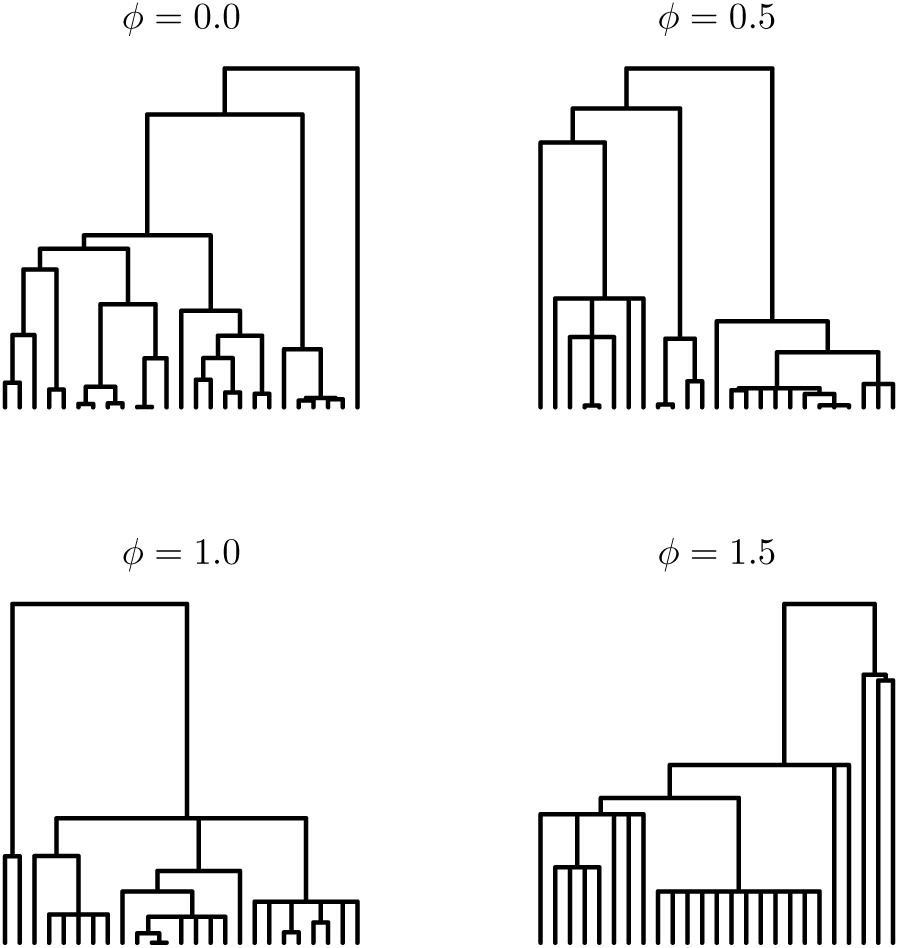
Random samples from the symmetric beta coalescent trees as the parameter *φ*, which governs the shape, increases from the Kingman value of zero. The model is described below, in the section on Genealogical Models. Note that as *φ* increases the number of multiple collisions increases, and the proportion of the total branch length in the external branches also increases. Our goal is to estimate *φ*, and its uncertainty, from site-frequency spectra generated from a mutation process on the trees. In this figure, we provide a single sample for each parameter value. In Supplement A, we provide twelve samples for each parameter value, to give a sense for the variation that still exists for a fixed parameter value.

In this paper we assume given a parametric model of Λ-coalescents, with shape parameter *φ*, and develop methods for estimating *φ* from the site-frequency spectrum (SFS) of a sample of aligned sequences. Specifically, we define point estimates and confidence intervals for *φ*, and calculate *p*-values for the hypothesis that the data are described by the Kingman coalescent. We provide general methods that can be applied to specific models, and analyze the results in two specific models. We develop the methods using synthetic data and then apply the methods to mitochondrial SFS from the Atlantic Cod.

We give a brief description of our problem, providing more precise definitions below. Our data consist of *n* aligned DNA or RNA sequences, which differ at *S segregating sites*, i.e., sites at which not all sequences have the same nucleotide. We may also have an aligned outgroup sequence. Let *S_k_*, for *k* = 0, 1, 2, …, *n*, be the number of sites where *k* of the *n* sequences are mutated relative to the sequence of their most recent common ancestor (MRCA), i.e., the sequence of the root of the coalescent tree. The SFS is the vector 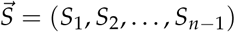. In practice we are not given the root sequence directly, as part of the data, so we need to either estimate it using an outgroup or use a reduced description of the data, known as the “folded SFS,” that does not depend on the root sequence. These issues are discussed below. We assume throughout that at most one mutation can occur at each site; this is the so-called “infinite-sites” model (Kimura, 1969). We also assume the mutations are neutral, so that they appear randomly on the tree, with constant rate proportional to the branchlength. We are primarily interested in *linked* sequence data, i.e., data arising in populations for which recombination can be neglected. The significance of this assumption from an inference perspective is that the data are described by a single coalescent tree.

The SFS provides information about the shape of the tree. In the infinite-sites model, if a mutation occurs on a branch that subtends *k* leaves, then the mutation will be seen in exactly *k* sequences, and will add one to the value of *S_k_*. Therefore, the value of *S_k_* reflects the fraction of the tree that subtends *k* leaves, and that fraction reflects the shape of the tree. The relation between the coalescent tree, the locations of the mutations, and 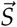, is illustrated in Figure 2.

**Figure 2:**
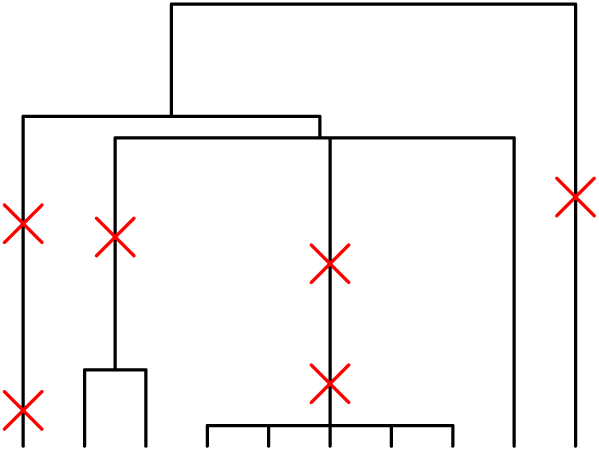
Random mutations on a coalescent tree with *n* = 10 and *S* = 6. The SFS for this data is 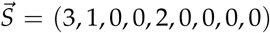. Note that when two mutations occur on the same branch they are assumed to occur at different sites, because of the infinite-sites assumption. In comparing with Figure 1, one can see that as *φ* increases, random mutations are increasingly likely to occur on external than internal branches. Thus, as *φ* increases, 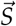 becomes increasingly weighted towards smaller *k* values, and this change can be used to infer *φ* from 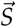.

There are two types of randomness that affect the uncertainty in our estimates of *φ*. The first comes from random variation in the tree, and the second from random variation in the location of the mutations on that tree. Suppose first that we knew the tree exactly. We would still be uncertain about the value of *φ*, because the coalescent process generates a tree randomly, and nearby values of *φ* can produce the same tree, albeit with varying probability. We call this uncertainty the *tree-based uncertainty* in *φ*. Although it is not obvious, it seems to be generally true, and has been proven for some models, that the tree-based uncertainty can be reduced by increasing *n*.

In practice we do not know the tree; we only know the SFS. Although the SFS provides information about the tree shape, this information is noisy, because the mutations land randomly on the tree. The fraction of the tree that subtends *k* leaves is approximately given by the ratio *S_k_*/*S*, but this ratio is subject to large random fluctuations when *S* is small. Thus, the randomness of the mutation process leads to uncertainty in the tree, and this uncertainty translates into additional uncertainty in *φ*. We call this uncertainty the *mutation-based uncertainty* in *φ*. It can be reduced by increasing *S*, which can be increased, in turn, by increasing the length of the sequences.

We therefore have two ways of decreasing our uncertainty in *φ*: increase *n* and increase *S*. In collecting data, we may well face a tradeoff between the number and length of the sequences, and we would like to collect the data so as to minimize the uncertainty in our inferences. But the relative importance of these two aspects of the data is not at all obvious. One of the objectives of this paper is to provide methods for assessing this tradeoff.

We now briefly discuss the statistical issues involved in making parameter inferences. In order to make parameter inferences in either a frequentist or Bayesian framework, one often calculates the statistical *likelihood*, which measures how likely it would be for the observed data to arise for a particular parameter choice. In symbols, if the statistical model of the data *D* is *p*(*D*|*φ*) for a fixed parameter value *φ*, then the likelihood is the same function, considered as a function of *φ* for the observed data *D*. Many of the most powerful tools in computational inference, including Markov Chain Monte Carlo (MCMC) approaches in Bayesian inference (Robert and Casella, 2013), assume that the likelihood can be readily calculated.

The likelihood in our problem, however, is not readily calculated. The problem is that the data are not expressed directly as a function of the parameter, but indirectly as an integral over the coalescent tree. Schematically,

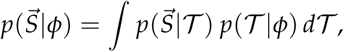

where 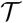 is the coalescent tree. (More detailed formulas will be provided below.) Although it is straightforward to sample from the distribution of 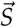, it is difficult to evaluate the functional dependence of 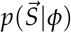 on 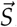 for fixed *φ*, or vice-versa. The difficulty in estimating 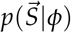by straightforward Monte Carlo sampling methods is that the output space is high-dimensional, so an impracticably large number of samples is needed, unless the output has some special structure that can be exploited.

Our problem falls into a general framework, in which the statistical parameter governs a latent intermediate process, and the observed data contains partial information about the latent process. Since the data could have arisen from many different realizations of the latent process, its probability is obtained as an integral over that process. In other problems, the latent structure might be a modeling construct, such as a state-space or a latent-variable model. This framework is an important special case of what are sometimes known as models with “intractable likelihoods.” Methods for dealing with such models include importance sampling (Robert and Casella, 2013, Stephens and Donnelly, 2000), Approximate Bayesian Computation (Beaumont, 2010, Marjoram et al., 2003), and composite likelihood (Larribe and Fearnhead, 2011, Lindsay, 1988, Varin et al., 2011), all of which have been used extensively in computational statistical genetics.

In this paper, we will use the method of *estimating functions* to tackle our intractable likelihood, a method that does not seem to have been applied previously in statistical genetics, and which may prove useful in other applications in the field. The method of estimating functions was pioneered by Godambe (Godambe, 1960, Godambe and Heyde, 1987), and can be interpreted as a generalization of maximum-likelihood (ML-) inference. The estimating functions are generalizations of the score function, which is the gradient of the log-likelihood. Estimates are obtained by setting the estimating function to zero.

Composite likelihood (Lindsay, 1988, Varin et al., 2011), which *has* been widely using in statistical genetics (Larribe and Fearnhead, 2011), is a special case of the method of estimating functions. In composite likelihood, the estimating function is a linear sum of the score functions of marginal and conditional distributions of the original model. Applications in statistical genetics include estimation of recombination rates (Fearnhead and Donnelly, 2002, Hudson, 2001), genetic mapping of traits (Larribe and Lessard, 2008), and detection of genes under selection (Gray et al., 2009). Some of our estimating functions will be composite score functions, and would therefore fall into this framework. It turns out, however, that composite likelihood is not sufficiently general for some of our models, which is why we have gone to the more general framework.

Of course, one cannot just replace the score function with an arbitrarily chosen estimating function and expect to achieve good results. We are interested in estimators that are *consistent*, in the sense that they converge asymptotically to the correct value, and *efficient*, in the sense that they make good use of the data. If an estimating function is *unbiased*, in a technical sense defined below, then it is guaranteed to be consistent. Also, formulae are available for assessing the efficiency of the estimator. The power of the method stems from the fact that it is frequently possible to define estimating functions that are statistically consistent and efficient, but which are not burdened with the full complexity of the original model.

In this paper, we obtain our estimating functions as the score functions of models that are simplified approximations of the original model. Such models are often called misspecified models, where a model is *misspecified* if the data do not arise from the model for any value of the parameter. We consider two different types of misspecifications.

In the first type of misspecification, we assume that the *S_k_* are statistically independent for different *k*. In fact, the *S_k_ are* approximately independent (Birkner et al., 2013), but they are not strictly independent, because the tree structure introduces correlations. This type of misspecification leads to a unbiased estimating function which is a composite score function, and therefore fits into the framework of composite likelihood. In the second type of misspecification, we treat the sites as unlinked, even when the data itself is linked. This unlinked model is not associated with a composite likelihood of the original linked model. We show in Appendix B, however, that the corresponding score function is nevertheless an unbiased estimating function for the original model.

Statistical inference for generalized coalescents has previously been studied in numerous papers. To deal with the intractable likelihood, the full likelihood has generally been replaced with a marginal likelihood using only part of the SFS, typically the “singletons,” i.e., the sites where only a single sequence is mutated. To be precise, the number of singletons are known and the number of non-singletons are known, but the details of the non-singleton spectrum are not used. The output space is now low-dimensional, and the functional dependence on the model parameter can be estimated computationally and inverted to form the marginal likelihood.

As we will see, inference based on singletons alone can work quite well. There are a number of reasons, however, why this approach is not completely satisfactory, and why it would be useful to have estimates based on the whole SFS. First, it is not clear *a priori* how much information is lost when the non-singleton counts are lumped together. Second, inferences lumping the non-singletons cannot assess how well the model fits the non-singleton portion of the data. Finally, the singleton count is often suspect, due to properties of the sequencing methodology or the sequence cleaning algorithm. For this reason Achaz (2008), for example, has advocated basing inferences on everything but the singletons. It is not clear whether this procedure would be feasible for our problem, or whether the data requirements would be prohibitive. If such estimates *are* feasible, it would also be interesting to know whether, in applications to real datasets, parameter estimates based on the singleton data are consistent with estimates based on the non-singleton data. If the two estimates give inconsistent values it would indicate a problem, either with the data or with the fit of the model to the data.

Another important feature of our approach is that it provides confidence intervals (CI’s) and not just point estimates. Previous studies have mostly only provided point estimates, although in some cases a likelihood surface was obtained, which could be used to construct CI’s. In particular, Birkner et al. (2011) and Steinrücken et al. (2013) generalized the genetree method of Griffiths and Tavaré (1994) and Stephens and Donnelly (2000) to Λ-coalescents, using importance sampling methods of Hobolth et al. (2008). The method involves the full sequence data, and leads to likelihood surfaces for *α* and *r*, where *α* is the parameter of the coalescent model, and *r* is a parameter that scales the tree size. The complexity of modeling the full genetree, however, limits the application of the method to datasets with fewer than about 200 sequences.

As noted, our analysis takes linkage into account. In most previous studies (Bhaskar et al., 2015, Birkner et al., 2013, Spence et al., 2016), it has been assumed that the sites are statistically independent, i.e., unlinked, which corresponds to the limit of infinite recombination. In many cases, the sites can indeed be treated as unlinked, and this model is appropriate. If the sites are linked, however, then the model is ignoring the randomness in the coalescent tree, which is often the largest uncertainty in the problem. It is then an open question how good the inferences will be. As noted above, we have shown that the unlinked estimator, when appropriately defined, is statistically consistent. Even so, the variance of the estimator may be much larger than could be obtained with a more accurate model. One of the goals of this paper is to provide guidance as to whether the unlinked estimator is useful for linked data.

In this paper, we apply our method to two models of generalized coalescents: the space of symmetric beta coalescents (Bertoin, 2010, Gnedin et al., 2014, Schweinsberg, 2003), and the space of Eldon-Wakeley coalescents (Eldon and Wakeley, 2006). These are both two-parameter models, with one parameter characterizing the shape of the coalescent and another the scale. We restrict ourselves to reduced versions of these models that depend only on a one-dimensional shape parameter, although our general approach is applicable to models with vector parameters, which may involve both shape and scale. For each of the models studied, we investigate the effect of varying *n* and *S* on the uncertainty of the estimator using synthetic data. Finally, we use our approach to analyze mitochondrial sequence data from the Atlantic cod.

## 2. Statistical Models

In this section we describe the statistical models we will analyze in this paper. We begin by defining the mutation process as a Poisson process on the coalescent tree, and the way in which random mutations generate a random SFS. We then describe how a random coalescent tree is generated from a set of rates. Combining the tree and mutation processes, we define our fundamental stochastic model, Model F, for the SFS for linked sequence data, as well as the analogous model for unlinked data.

Model F depends on both the mutation rate and on parameters governing the shape and size of the coalescent tree. By conditioning on the number of segregating sites, we can define reduced models that depend only on the shape parameter of the coalescent model. In so doing, we simplify the inference problem, but the simplification does involve an approximation. We show in the main text how conditioning can be interpreted as a modification of the experimental procedure, and in Appendix A how the conditioned likelihood can be derived from the approximation that the shape and the size of the coalescent tree are independent.

In this paper, we analyze only the reduced models, although the fundamental model could be analyzed in a similar fashion. We derive four reduced models, three of which assume linked data and depend on either all of the SFS data (Model A), on singletons only (Model S), or on non-singletons only (Model N), and one which assumes unlinked data and depends on all of the SFS data (Model U). The fundamental model, though not used for statistical inference, is needed for deriving the reduced models.

### 2.1. Mutation model

Let 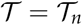 be a coalescent tree with *n* leaves, where we will generally suppress the *n* for brevity. Let 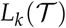 be the total length of all branches subtending *k* leaves, *k* = 1, …, *n* − 1. Let

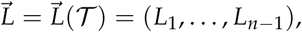

and let 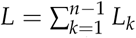 be the total tree length. In place of 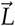, it is often convenient to use the variables 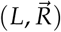, where 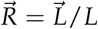. Thus, *R_k_* = *L_k_*/*L* is the *fraction* of the tree’s branch length that subtends exactly *k* leaves, and 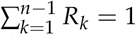.

We assume that the coalescent tree describes the genealogical history of a set of genetic sequences, which we assume initially to be of fixed length. We assume that the sequences are aligned, and say that a sequence is mutated at a site if it differs at that site from the MRCA sequence (or *root sequence*). A site at which *k* of the *n* sequences are mutated is of *size k*. Let *S_k_*, *k* = 0, 1, 2, …, *n*, be the number of sites of size *k*. We are mostly concerned with the *segregating sites*, which are sites of size *k*, with 1 ≤ *k* ≤ *n* − 1. Each *segregating site* divides the set of samples into two proper subsets, corresponding to the sequences that are or are not mutated at that site. The total number of segregating sites, *S*, is defined as 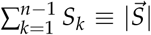, and the SFS is defined as 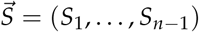. When the limits of a sum or product over *k* are not explicitly given, they are assumed to extend from *k* = 1 to *k* = *n* − 1.

We assume that in the genealogical history of the sample at most one mutation has occurred at each site; this is known as the infinite sites mutation model (Ewens, 2004, Kimura, 1969). This assumption implies, in particular, that if a mutation occurs in the tree, all descendants will possess that mutation, because reversion to the original form would require a second mutation at that site. Similarly, there can be at most two different nucleotides at any site, because additional nucleotides would require additional mutations at that site.

The SFS depends on the root sequence, which is not part of the sequence data, so it is not fully defined by the data alone. There are two ways of dealing with this issue. First, if we have an outgroup sequence, we can choose to use that sequence in place of the root sequence. If the infinite sites model is valid, there can be no mutations at a segregating site between an outgroup and the root, so the outgroup and root sequences will give the same SFS. Second, if an outgroup is not available, or if we do not trust the infinite-sites assumption between the root and the outgroup, we can “fold” the SFS by lumping together *S_k_* and *S_n_*_−*k*_; see Eq. 14 for details. The folded sequence does not depend on the root sequence, so it is well-defined even when the root sequence is unknown, although some information is lost.

We assume that mutations are neutral, so that they do not affect the genealogical process and can be modeled independently of that process. We assume that mutations arise randomly at a uniform rate, which implies that mutations can be described as a Poisson process with rate *μ*, say, per unit branch length. By the infinite sites assumption, a mutation on an edge subtending *k* leaves will induce a segregating site of size *k*. Thus, the number *S_k_* of segregating sites of size *k* is a Poisson process with rate *μL_k_*, and the probability of 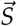is

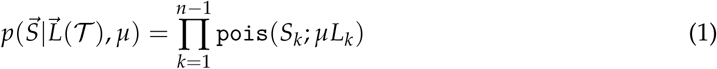

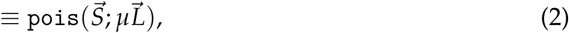

where pois is the Poisson distribution:

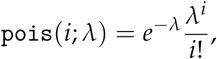

and where for vector arguments, pois is defined as the product over the components:

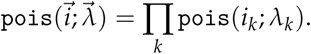

Note that in the present context, in which we assume that the tree 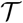 is fixed, the individual *S_k_* are statistically independent.

### 2.2. Tree model

To obtain the distribution of 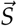 for a population of trees we need to specify the tree distribution. In this paper, we will focus on a particular class of random trees, although the methods we develop can be applied more generally. Specifically, we assume that the trees are randomly generated by a coalescent process in which each coalescent event involves the merger of *k* > 1 lineages into a single lineage. These processes are called Λ-coalescents. We assume that the coalescent rate for all *k*-subsets of *m* lineages is the same. The rate for a coalescent event involving *k* lineages, when there are a total of *m* ≥ *k* lineages, is written generically as *λ_m_*_,*k*_.^1^ These rates cannot be chosen arbitrarily, however, because they must obey consistency conditions. It turns out that any consistent family is characterized by a finite measure Λ on the unit interval [0,1], and that conversely, any such measure defines a consistent family (Pitman, 1999). Given Λ, the rate for a coalescent event involving *k* of *m* lineages is^2^

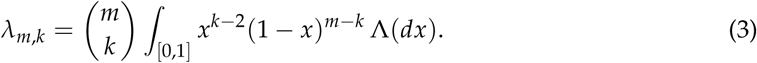

In general, different Λ will lead to coalescent processes with different shapes, or more precisely, different shape distributions. However, if one measure is just a scalar multiple of another, 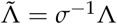, the shape distributions will be unchanged, except that the branchlengths will be scaled by the factor *σ*. For this reason, we focus on normalized coalescents with Λ([0, 1]) = 1, which is equivalent to assuming that *λ*_2,2_ = 1. Any Λ-coalescent can be obtained from a normalized Λ-coalescent through scaling. Within the class of normalized coalescents, we consider parameterized families Λ*_φ_*, where *φ*, which may be a vector, is called the *shape* parameter. To accommodate different scalings, we define

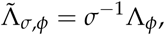

where *σ* is called the *scaling* parameter. Then

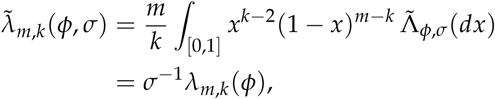

where *λ_m_*_,*k*_(*φ*) is the rate for Λ*_φ_*. We write 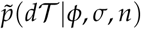 for the distribution of trees with *n* leaves corresponding to a Λ-coalescent with measure 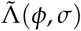, and 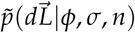 for the induced probability on 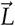. Similarly, we write 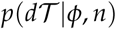 and 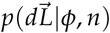, *i.e.*, without the tilde, for the analogous quantities for Λ*_φ_*.

### 2.3. Fundamental model

Integrating over the tree distribution, we obtain the distribution of 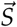 given the model parameters, *μ*, and *n*:

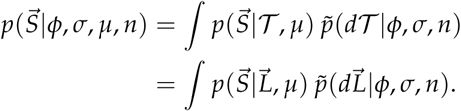

We can simplify this expression by noting that both *μ* and *σ* are scaling parameters, and can be combined as the product *ψ* = *μσ*. Noting that 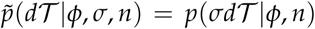, we can write the model more concisely as

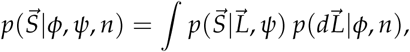

where 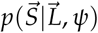is of the form (1), with *ψ* replacing *μ*, or more simply, as

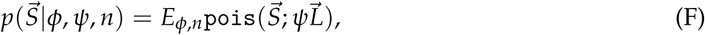

where *E_φ_*_,*n*_ is the expectation over the measure 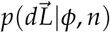. This is our fundamental model for 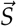; we call it Model F.

The right-hand-side of Eq. F is the expectation of the product of Poissons, one for each component of the data vector. The form of the model is the same if we wish to model only some of the *S_k_*, or if we wish to model binned subsets of the *S_k_*. In the first case, the claim can be established either by starting with the original model and marginalizing away the variables we wish to exclude, or simply by writing down a new model for the variables of interest. Thus, for example, if we are only interested in the probability of the non-singletons, (*S*_2_, ⋯, *S_n_*_−1_), we need only omit *S*_1_ from the product of Poissons. In the second case, the model for the binned variables follows from the fact that the sum of Poisson random variables is also a Poisson random variable, whose rate is the sum of the rates of the summands. Thus, for example, if we let 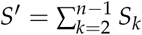 and similarly for *L*′, then

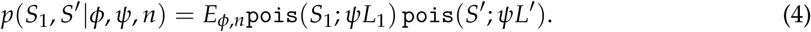

When the sites are fully unlinked, each site has its own coalescent tree, which is statistically independent of the trees at all other sites. To implement the infinite sites approximation, we assume that there are *m* sites, where *m* is very large, and that the mutation rate *per site* is *μ_s_* = *μ*/*m*, so that the probability of more than one mutation at a single site is vanishingly small. Define *ψ_s_* = *μ_s_σ* = *ψ*/*m*. Then, at any site *j* and for large *m*,

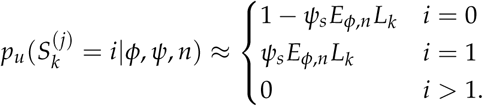

For fixed *i*, the total probability of size *k* mutations over all sites is given by the binomial theorem,

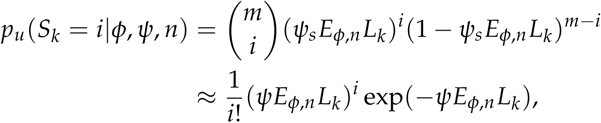

where the latter expression holds in the limit as *m* → ∞, and we have used the facts that in this limit, 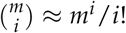 and 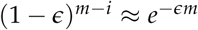 when *∊* ≪ 1. It follows that in this limit,

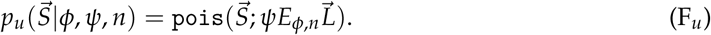

Note that Model F*_u_* is identical to Model F, except that the expectation has been taken inside the Poisson distribution. Thus, the basic unlinked model, and all models derived from it, depend only on the expected branch lengths, and are insensitive to any fluctuations in the coalescent tree.

### 2.4. Reduced models

We will not analyze Model F (or its unlinked counterpart) directly in this paper, although it would be interesting to do so and the model is amenable to our methods, as will become apparent later. Instead, for simplicity and computational tractability, we use it to derive models depending only on the shape parameter *φ*, which should nevertheless be useful in the analysis of real datasets. This simplification is achieved by assuming that *S*, the total number of segregating sites, is fixed. Mathematically, this has the effect of replacing the multivariate Poisson distribution with a multinomial, and more importantly, of causing both the mutation rate and the scaling of the tree length to drop out of the equation. The reduced models we consider can be derived either as approximations to our fundamental model, or as the correct models for a modified experimental setup that arguably would not affect the results very much. We describe the latter here, and refer the reader to Appendix A for the former.

In deriving the model above, we have assumed that the sequences are of fixed length, which means that the total number of segregating sites *S* will vary somewhat with each sample of *n* sequences. Suppose that instead, we fix the number of segregating sites we want to measure and allow the length of the sequences to vary instead. To implement this approach experimentally, we could continue reading additional sites of the sequences until the predetermined number of segregating sites has been reached. This is analogous to the difference between flipping a coin a hundred times and flipping the coin until fifty heads have appeared. In both cases, we could make inferences about whether or not the coin was fair, but the details of the mathematics would be somewhat different.

The distribution of 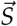, conditioned now on the value of *S*, is

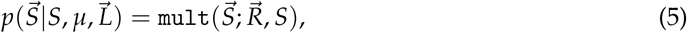

where

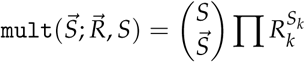

is the *multinomial distribution*. The *multinomial coefficient* is defined by

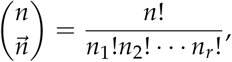

where 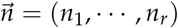 and 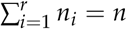. Here, and henceforth, we use the convention that 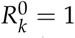 even if *R_k_* = 0. Eq. 5 follows from the well-known fact that a Poisson conditioned on the number of events is the multinomial; the formula can be derived explicitly by dividing Eq. 1 by

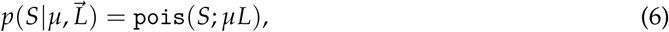

which is obtained by summing Eq. 2 over all 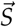 with 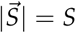. Note that the right hand side of Eq. 5 depends on neither *μ* nor *L*, so we may suppress these variables from the left hand side as well. Integrating over the tree distribution (which now means integrating over 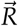), we get

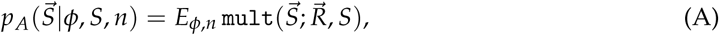

where the only unknown parameter is *φ*.

Eq. A is the first of the four models we shall analyze in this paper; we call it Model A, because it takes *all* the data into account. We now describe Model S, which is based only on the number of singletons and non-singletons. In Model S, we lump all *S_k_* with *k* > 1 into a single observable 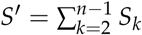. The model is

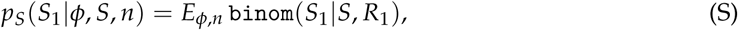

where the binomial distribution is

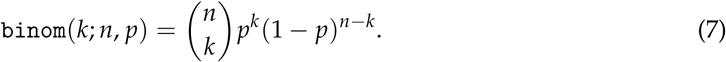

This model can be obtained either from Model A or directly from Model F. The purpose of studying Model S is to compare it with Model A, and to see if knowledge of the size spectrum of the non-singletons tells us anything more than simply knowing the total number of non-singletons. If the results are similar, we can save effort by using Model S.

With Model N (for “non-singletons”) we are interested in assessing the information contained in the non-singletons alone. Letting 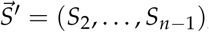, as above, we start from the fundamental probability distribution for 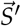,

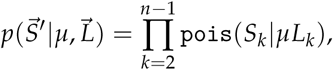

and again imagine an experiment in which sites are added until the total number of *non-singleton* segregating sites reaches some fixed number *S*′. By the same argument used previously, we get

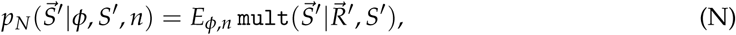

where now 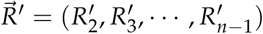, with

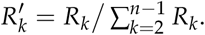

Note that 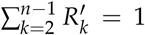. Note also that although the approximation is formally the same as in Model A, it *is* a different approximation, and its accuracy must be assessed separately.

Finally, if we start with the unlinked version of the fundamental model and condition on *S*, we obtain Model U (for “unlinked”):

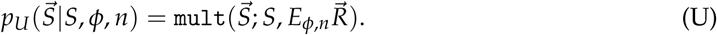

This model will be used to test whether we can make useful inferences on linked data using an unlinked model. The appeal of Model U, of course, is that the unlinked model is much simpler to evaluate computationally than the linked model.

## 3. Inference

The problem we address in this paper is making parameter inferences from linked SFS data. Specifically, we assume that we have an 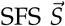, which is well-described by one of our reduced models for some parameter value *φ*_0_. We wish to deduce point estimates and confidence intervals for *φ*_0_. In order to determine the ML-estimate we need to calculate the likelihood, and this is sufficiently complex that it must be done computationally.

Taking Model A as an example, the likelihood is

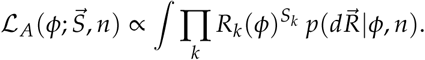

In order to calculate the likelihood we need to integrate over the “nuisance variable” 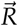, which describes the tree shape. We might hope to approximate 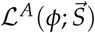 by Monte Carlo integration:

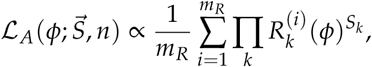

where 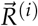 corresponds to the *i*th of *m_R_* samples from 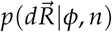.

Unfortunately, there are difficulties with performing this calculation. In typical models, each tree sample will have 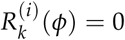 for most of the larger values of *k*. For any value of *k* with *S_k_* > 0, the sample will contribute zero to the likelihood unless 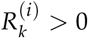. If we were concerned only about a single *k*, this problem would be manageable. The real problem is that we are computing the likelihood from the entire site frequency spectrum, so we must simultaneously have 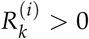 for all *k* values with *S_k_* > 0. Thus, the probability of obtaining a non-zero contribution is a small number *to a power*, which becomes vanishingly small for typical samples of 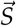, for even moderate values of *n*. In statistical parlance, any particular tree imposes *structural zeros* on the categorical data vector 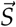, i.e., values of *k* for which *S_k_* is necessarily zero. The problem, stated concisely, is that the data will generally be incompatible with the structural zeros of the vast majority of the sampled trees.

### 3.1. Unbiased estimating equations

Our challenge is to estimate the model parameter *φ* when we cannot calculate the likelihood. Our strategy is to replace the original model with a misspecified model whose likelihood we *can* calculate. We can then estimate *φ* by maximizing the likelihood of the misspecified model. It is not at all clear that this will work, and in general, it won’t (Freedman, 2006). In an effort to choose useful misspecifications, we will require that the estimator be consistent, i.e., that it converge to the correct model parameter in the limit of large sample size. To ensure consistency, we require that the score function of the misspecified model be an unbiased estimating function (defined below) for the original model.

We give a brief overview of unbiased estimating equations, and then show how we apply this approach to coalescent inference. This framework is not as well-known as composite likelihood, and an accessible presentation does not seem to be available in the existing literature. Since this framework may have other applications in statistical genetics, it seems useful to outline the basic theory. The reader who is not interested in the statistical details may wish to skip ahead to the subsection, “Application to coalescent inference,” which summarizes the estimators we will use later in our analysis.

For clarity, it is helpful to work in a general setting. Let *p*(*x*|*θ*) be a statistical model, where *θ* is the parameter vector and *x* the data vector^3^. For fixed *θ*, *p*(*x*|*θ*) is the distribution of the data for parameter *θ*, and for fixed *x*, *p*(*x*|*θ*) is the (unnormalized) *likelihood* for data *x*. If *x* is a discrete parameter then *p*(*x*|*θ*) is a probability, and if *x* is continuous it is a density with respect to some appropriate base measure *dx*. We assume that the parameter *θ* ranges over some subset of *p*-dimensional Euclidean space.

Let *g*(*x*, *θ*) be a *p*-dimensional vector function of the data and the parameter. Given a sample *X* from *p*(*x*|*θ*_0_), an estimate 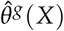 of *θ*_0_ is obtained as the solution to the *estimating equations*,

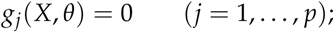

*g* itself is called an *estimating function* (Godambe, 1960). Given (*X*_1_, …, *X_N_*), where the *X_i_* are *N* independent samples from *p*(*x*|*θ*_0_), an estimate 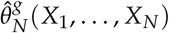 is obtained as the solution of

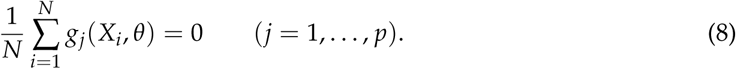

In general, the equations may have multiple roots. We assume whatever regularity is needed to ensure a unique solution in the limit of large *N*; see Yi and Reid (2010) for details. We sometimes write 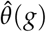 for 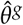.

We are interested in the properties of such estimators—whether they converge to the true value *θ*_0_, and how much they vary. An estimator sequence *δ_N_*, say, is statistically *consistent* if *δ_N_*(*X*_1_, …, *X_N_*) → *θ*_0_ in probability as *N* → ∞. Godambe has investigated a natural sufficient condition for 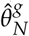 to be consistent. Call *g* an *unbiased estimating function for p* if

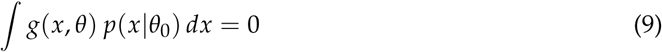

for *θ* = *θ*_0_, for any choice of *θ*_0_. Then 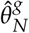 is consistent, and this is the first key result of the theory. Formally, Eq. 9 is the *N* → ∞ limit of Eq. 8. Thus, the value of *θ* that solves Eq. 9 is just the limit of the estimates 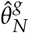 as *N* → ∞. If this value is *θ*_0_, then 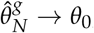, which is the definition of consistency. For a rigorous proof in a more general setting, which also provides a sufficient set of regularity conditions, see Yi and Reid (2010). The condition can be stated more simply (but somewhat more opaquely) as follows:

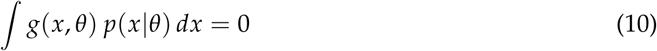

for all *θ*.

The asymptotic sampling variance of an estimator sequence 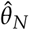 is defined as

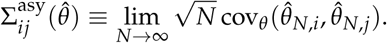

The asymptotic sampling variance of 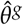 can be computed directly from *g*; this is the second key result of the theory. Let

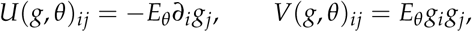

and define the Godambe information as

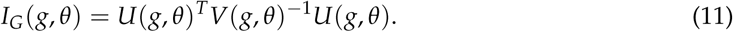

Then 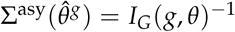, the inverse of the Godambe information, provided *θ* is in the interior of its domain of definition (Yi and Reid, 2010). We write Σ*_G_*, or 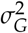 in one dimension, for 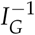.We sometimes suppress the *θ*-dependence in the notation when we are primarily interested in the *g*-dependence, and vice-versa.

#### 3.1.1 Maximum Likelihood

A first example of an estimating function is the well-known (*efficient*) *score function* of standard maximum-likelihood estimation. In fact, the method of unbiased estimating equations can be seen as a generalization of maximum likelihood, and for understanding this generalization it is very useful to understand the most important special case.

Let ℓ(*θ*; *x*) be the log-likelihood functions for *p*(*x*|*θ*),

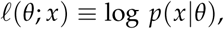

and define *g_p_*(*x*, *θ*) ≡ ∇*_θ_ℓ*(*θ*; *x*); *g_p_*(*x*, *θ*) is called the *score function*. Given data *X* generated by *p*(*x*|*θ*_0_), the *maximum likelihood (ML-)* estimate of *θ*_0_, 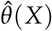, is the value of *θ* maximizing the likelihood, or equivalently, the log-likelihood. Given sufficient regularity, the maximum of the log-likelihood is achieved where the slope of its derivative, i.e., the score function, is zero:

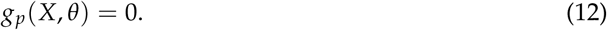

Eq. 12 is known as the *likelihood equation*. The likelihood equation is a special case of an estimating equation. The likelihood equation for a set of *N* i.i.d. samples is just given by the general formula, Eq. 8.

It is well-known (and easy to show) that *g_p_*(*x*, *θ*) satisfies Eq. 10; this is sometimes known as Bartlett’s first identity. Thus, the score function is an *unbiased* estimating function, and the consistency of maximum likelihood follows from the general result for unbiased estimating equations. With respect to the asymptotic variance, it is easy to show that 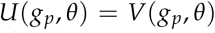 (Bartlett’s second identity), so that by Eq. 12, 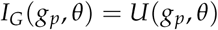. But 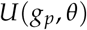 is just the Fisher information, which is defined as

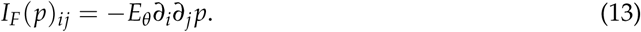

Thus, the classical result that the asymptotic variance of the ML-estimator is the inverse Fisher information is a special case of the general result for an unbiased estimating function, and the Fisher information is a special case of the Godambe information. We write 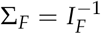 (or 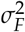 in one dimension) for the inverse Fisher information.

Among all unbiased estimating functions, the score function is distinguished by the fact that it has the smallest asymptotic variance (Godambe, 1960). The *efficiency* of *g* is defined as *I_G_*(*g*)/*I_G_*(*g_p_*), or equivalently, *I_G_*(*g*)/*I_F_*(*p*), which can be no greater than one, a result that generalizes the Cramer-Rao inequality.^4^

#### 3.1.2 Composite Likelihood

As a second example of an estimating function, consider the class of all finite linear combinations of the score functions of any marginal or conditional distribution of *p*. It is easy to show that any marginal or conditional distribution of *p* will satisfy Eq. 10. Therefore, any linear combination will also satisify Eq. 10, and will define an unbiased estimating function.

Such linear combinations are called *composite score functions* (Lindsay, 1988, Varin et al., 2011), and these form a subclass of the class of unbiased estimating functions. It is a proper subclass because, if *g* is the weighted sum of components *g_i_* and *g* is an unbiased estimating function, there is no need for the individual *g_i_* to come from marginal conditional distributions, or even to be unbiased estimating functions. The use of such estimators is generally described as the method of *composite likelihood*. The *composite log-likelihood* is the corresponding sum of the associated log-likelihoods. As noted in the introduction, composite likelihood is widely used in statistical genetics, usually for the purpose of parameter estimation when the full likelihood is intractable.

In this paper, the composite log-likelihoods will all correspond to misspecified models when exponentiated. In particular, all of the weights will be equal to one. It is easy to see, however, that a weighted sum of marginal and conditional log-likelihoods will not usually correspond to any statistical model. Furthermore, the efficiency of the estimator (see below) can often be improved by adjusting the weights (Lindsay, 1988, p. 229). We have not yet explored this possible improvement.

#### 3.1.3 Misspecified models

Finally, we consider the score functions obtained from misspecified models, which are of particular interest in this paper. We reserve the symbol *p* for the “correct” model, i.e., the model that is assumed to correctly describe the complete data vector for the true but unknown parameter value *θ*_0_. We use the symbol *q* for any other model, which is possibly a misspecified model. By *misspecified* we mean that the model does not correctly describe the data for any parameter value *θ*. We assume that the allowed values of *θ* are the same for all models, and that if *p* and *q* are densities, the base measure *dx* is the same for both models. The definitions of the likelihood and score functions, given above for *p*, carry over for *q*. To distinguish the correct model from the misspecified model, we use subscripts: *ℓ_p_* and *ℓ_q_*, *g_p_* and *g_q_*, etc.

Of course, the score function *g_q_* will not necessarily be an unbiased estimating function. We say that a model *q* is an *unbiased misspecification of p* if it is misspecified and its score function *g_q_* = ∇*ℓ_q_* is an unbiased estimating function for *p*. Huber (1967) and White (1981, 1982) first showed that the asymptotic variance of 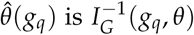, provided that *θ* is an interior point in its domain. As a result, the inverse of Eq. 11, when estimated with sample data, is often called the Huber-White “sandwich estimator” for the asymptotic variance. We emphasize that 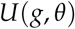 and *V*(*g*, *θ*), which are used to compute *I_G_*(*g_q_*, *θ*), are defined in terms of expectations with respect to the true density *p*(*x*|*θ*), *not* with respect to *q*(*x*|*θ*).

#### 3.1.4 Efficiency

All of the results in this section are valid regardless of how badly the models are misspecified. But there is nevertheless a penalty for a badly misspecified model: the efficiency of the estimator may be low. When the misspecification is mild we expect the efficiency to be high, but when the misspecification is severe we expect the efficiency to be low.

If *q* = *p* then 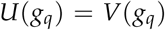, as noted above. In this case the efficiency is one, because the most efficient estimator is the ML-estimator for the original model. A discrepancy between 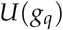 and *V*(*g_q_*) is a sufficient condition for misspecification, as emphasized by White (1982). It is not a necessary condition; we can have 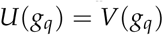 even if the model is misspecified. Nevertheless, we find in practice that our misspecified models do show discrepancies between 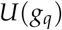 and *V*(*g_q_*), and that the discrepancy is larger when the model is badly misspecified.

To evaluate this discrepancy in one dimension, we will calculate the quantity

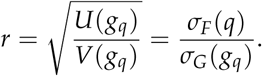

The numerator, *σ_F_*(*q*), would be the asymptotic standard deviation of the ML-estimator *if q were the correct model*, and the denominator, *σ_G_*(*g_q_*), is the actual asymptotic standard deviation of the estimator 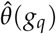. We find empirically that in our misspecified models *σ_F_*(*q*) < *σ_G_*(*g_q_*), *i.e.*, that the width of the distribution in the misspecified model is too small. This phenomenon is well-known in the context of composite likelihood; see Pauli et al. (2011) for a detailed analysis. Thus, we expect to find that *r* < 1 for a misspecified model, and that the value of *r* will indicate, at least roughly, the degree of misspecification. This analysis suggests, incidentally, that a Bayesian analysis that approximates the statistical model using *q* will tend to produce incorrectly narrow posteriors.

#### 3.1.5 Application to coalescent inference

In our problem there are two obvious simplifications that would facilitate the computation of the likelihood. The first is to treat the *S_k_* as independent, so that the misspecified model is the product of the individual marginal distributions:

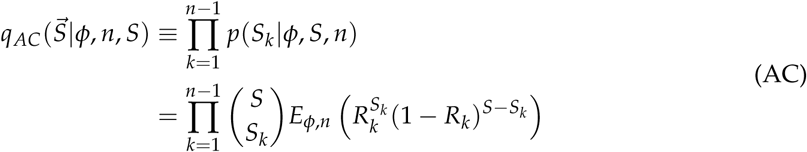

(Note that by taking the product of the marginal distributions, we change the model so that Σ *S_k_* is no longer necessarily equal to *S*.) We use the notation *ℓ_AC_* for the corresponding composite likelihood, and *g_AC_* for the corresponding composite score function. Thus,

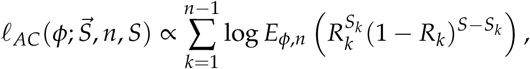

and

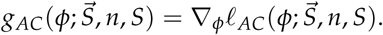

We write 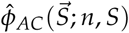 for the estimator of *φ* obtained either by maximizing 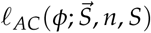or by setting 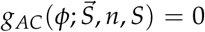. (These two conditions are equivalent provided that the estimator is continuously differentiable and unimodal. These conditions generally hold in our applications, except for the non-singleton likelihood, which is generically bi-modal; a discussion of this bimodality can be found below Figure 25 in Supplement B.) We often write 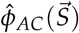, suppressing the functional dependence on *n* and *S* in the notation.

Similarly, if we are only interested in non-singletons, we consider a composite misspecification of Model N:

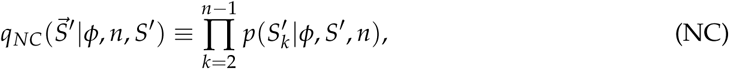

where *S*′ and 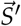 are defined above; *ℓ_NC_* and *g_NC_* are defined as the corresponding composite log-likelihood and composite score functions, in analogy to the corresponding definitions for AC. The corresponding estimator is written 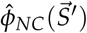.

It has been shown by Fu (1995) for the Kingman coalescent and by Birkner et al. (2013) for the symmetric beta coalescents that the *L_k_* are nearly pairwise independent, and it easy to confirm that the same is true of *R_k_* and *S_k_*. On this basis we expect that this composite likelihood will be only a mild misspecification of the linked likelihood, and that its efficiency should be high.

The second useful simplification is to replace the linked likelihood with the unlinked likelihood. Like the composite likelihood, the unlinked likelihood has the advantage that terms for the individual *S_k_* are largely independent, in that the *S_k_* values in a sampled 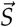 are not correlated by their mutual dependence on a sampled tree. (They are still correlated by the fact that they must sum to *S*.) The unlinked likelihood is *not*, however, a composite likelihood. It turns out, rather surprisingly, that it is nevertheless a consistent misspecification for Model A. We provide a proof in Appendix B. In practice, we will use the composite misspecification of the unlinked model:

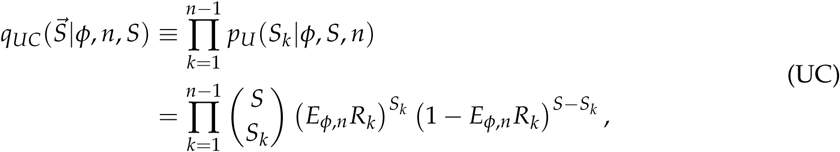

which removes the requirement that the *S_k_* sum to *S*. It turns out that 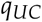 is also a consistent misspecification of Model A, although it again does not give a composite likelihood; see Appendix B. We write 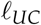 for the logarithm of 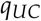, considered as a function of *φ*, 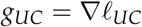, and 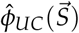 for the corresponding estimator.

The pseudolikelihood of Birkner et al. (2013) is essentially Model UC, but with *Z_k_* = *L_k_*/*E_φ_L* replacing *R_k_*, and values of *k* lumped above some threshhold. As the authors observe, *E_φ_Z_k_* ≈ *E_φ_R_k_* when *n* is large (Birkner et al., 2013, Figure S6), so this may be a useful approximation, particularly since there are now efficient algorithms for computing *Z_k_* (Spence et al., 2016). If we use this approximation, however, our consistency result no longer holds, so we have chosen to use *E_φ_R_k_* in this paper.

The unlinked model is a fairly drastic alteration of its linked counterpart. As we see below, tree fluctuations often contribute more randomness to the SFS than mutational fluctuations, at least for the values of *n* and *S* we consider, so the unlinked model discards most of the randomness in the original model. Thus, it is very interesting to study the uncertainty in the associated estimator, which can only be done through computation.

Finally, we consider inference based on Model S, using singletons alone. We set *ℓ_S_*(*φ*; *S*_1_, *n*, *S*) = log *p*(*S*_1_|*φ*, *n*, *S*),

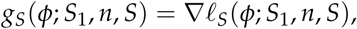

and 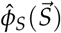 for the corresponding estimator. (Note that *S* is used in two different senses, as an abbreviation for “singletons,” and in 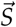 and *S_k_* to indicate “segregating sites.” The context will always make the meaning clear.)

#### 3.1.6 *p*-values and confidence intervals

In our applications, we will assume that the data 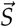 are well-described by a model of the form 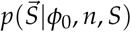, where *φ*_0_ is unknown. Our goal is to estimate *φ*_0_. We define *p*-values and confidence intervals, suppressing *n* and *S* in the notation for simplicity.

Let *H*_0_ be the hypothesis that the data are described by the Kingman Model. In both of the genealogical models we consider, which are defined below, the Kingman Model corresponds to *φ*_0_ = 0, which sits at the boundary of the parameter interval. Thus, we shall write *H*_0_ as the condition that *φ*_0_ = 0. Let 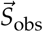be the observed SFS, let 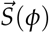 be a random variable with distribution 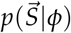, and let 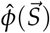 be an estimator of *φ*. The *p*-value for *H*_0_, given 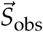, is the probability that the estimator for data generated with *φ*_0_ = 0 is at least as great as the observed value:

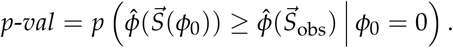

We say that *H*_0_ is *ruled out* if *p* < 0.05.

Let 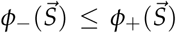be two functions of the data 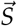 only, which do not depend on *φ*_0_. (In general, they will also depend on *n* and *S*, which are temporarily suppressed in our notation.) If 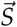 is sampled from the distribution 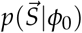, then for each 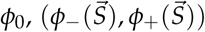 is a random interval. We say that this interval is a *confidence interval (CI) of confidence level* 1 − *γ*, or a 1 − *γ* CI, if it contains the true value *φ*_0_ with probability at least 1 − *γ*, for all values of *φ*_0_. We define CI’s so that for any *φ*_0_, the maximum probability that the interval is entirely to the left of *φ*_0_ is no greater than *γ*/2, and similarly for the probability that it is entirely to the right. We provide details in Appendix C, which also addresses a number of technical issues involved with *p*-values and CI’s.

## 4. Genealogical models

To illustrate our approach we focus on parameterized families of Λ-coalescents. We consider two such families, the symmetric beta coalescents and the Eldon-Wakeley coalescents. Both families are one-dimensional families interpolating smoothly between the Kingman and star coalescents, as explained below.

### 4.1. Symmetric beta coalescent

The two-parameter family of *beta coalescents* has Λ = *Beta*(*a*, *b*), where *Beta*(*a*, *b*) is the beta distribution, with density

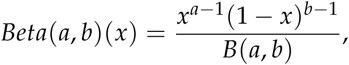

where *B*(*a*, *b*) is the beta function, Γ(*a*)Γ(*b*)/Γ(*a* + *b*). The beta function is only defined for *a* > 0 and *b* > 0. However, the beta distribution has a well-defined limit of *δ*_0_ as *a* → 0 (if *b* > 0), and a well-defined limit of *δ*_1_ as *b* → 0 (if *a* > 0), where *δ_q_*(*dx*) is the unit point mass at *q*. Therefore, *Beta*(*a*, *b*) is well-defined for any *a* ≥ 0 and *b* ≥ 0, except when *a* = *b* = 0.

The one-parameter family of *symmetric beta coalescents* has Λ = *Beta*(*a*, 2 − *a*), where 0 ≤ *a* ≤ 2. In this model, *a* is the shape parameter that we indicate generically by *φ*. For *a* < 2, these coalescents can be obtained from a generalized Moran model, in the limit of large population size, under the assumption that the tail of the offspring distribution *p*(*ν*) decays asymptotically according to a power law with exponent 2 − *a* (Bertoin, 2010):

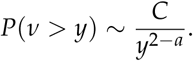

Here, ~ means that the ratio of the two sides of the equation approaches one in the limit of large *y*, and *C* is a constant. (It turns out that only the asymptotic form of the tail affects the limiting process.) The symmetric beta coalescent is often parameterized by *α* = 2 − *a*. For simplicity, we drop the “symmetric” qualifier when our meaning is clear. For *a* < 1, the beta coalescent can also be derived as an appropriate limit of the Wright-Fisher process with discrete generations (Schweinsberg, 2000).

The rates at the endpoints are

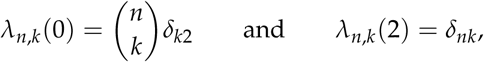

where *δ_ij_* = 1 if *i* = *j* and zero otherwise, which correspond to the Kingman and star coalescents, respectively.

For the symmetric beta coalescent, it has been shown for 0 ≤ *a* < 1 and fixed *k* that *R_k_* converges almost surely to an *a*-dependent limit as *n* → ∞ (Berestycki et al., 2014, Schweinsberg, 2010). In particular, *R*_1_ converges to *a*. As *n* becomes large, therefore, the uncertainty in *a* becomes progressively smaller, and the tree-based uncertainty in *a* is reduced, at least for 0 ≤ *a* < 1.

### 4.2. Eldon-Wakeley coalescent

The Eldon-Wakeley (EW-) or *δ*-coalescent, has Λ(*dx*) = *δ_ψ_*(*dx*) where 0 ≤ *ψ* ≤ 1. The rates are

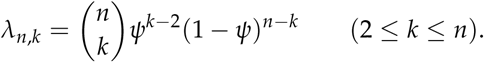

(Note that this *ψ* is different from the *ψ* used above as a scaling parameter for the tree size; the meaning will always be clear from context.) In the EW-coalescent, *ψ* is the shape parameter denoted generically by *φ*. The EW-coalescent is defined on the closed interval [0,1]. As with the symmetric Beta coalescent, the Kingman and star coalescents correspond to the endpoints of the interval, with *ψ* = 0 giving the Kingman coalescent, and *ψ* = 1 the star coalescent.

The EW- and beta coalescents both interpolate between the Kingman and star coalescents, but they differ in that the EW measure is concentrated on a point, and the beta coalescent is typically quite spread out over the unit interval. The log-likelihood can be defined at the Kingman endpoint, i.e., *ψ* = 0 or *a* = 0, but is often singular at the star endpoint, because the likelihood will be zero, and the log-likelihood −∞, if 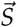 contains anything but singletons. Therefore, we include the Kingman endpoint in our analyses, but truncate the parameter range away from the star endpoint.

### 4.3. Folding

As noted above, unless an outgroup is known, it is not possible to distinguish the possibilities that a site with at most two character states contains *k* or *n* − *k* mutations. *Folding* is the process of combining these two types of sites for all *k*. If 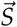is the original site frequency spectrum, formed by counting the differences from an arbitrarily chosen reference sequence, and 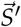 the corresponding folded spectrum, then

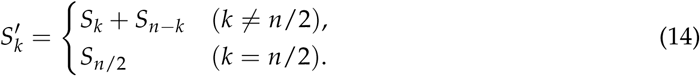

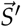 does not depend on the reference sequence. We will *always* assume folded spectra, in both our simulations and datasets. In our experience with synthetic data, inferences are little changed when the data are folded. In this comparison, however, we have used the correct root sequence. We have not assessed the impact of using an incorrect root sequence, which is the concern when an outgroup is used to infer the full SFS of real data.

## 5. Computation

Numerical methods are used for four purposes: (1) To sample from the tree distributions; (2) To compute the log-likelihoods of the misspecified models; through Monte Carlo sampling; (3) To evaluate the Godambe information, *I_G_*, which involves the expectation over the distribution of 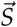. (4) To calculate the estimates of *φ*.

1. To sample the tree distribution we use the “lookdown algorithm” of Donnelly and Kurtz (1996); see also Berestycki (2009). The code was verified, in part, by comparing the expected values of the sampled 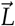 with the expected value given by the independent algorithm in Spence et al. (2016). To sample 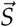 in the reduced models, we first sample 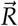 from the tree distribution, and then sample 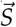 from the multinomial distribution. The code was checked by verifying the consistency of the estimators.
2. The log-likelihoods are calculated by Monte-Carlo sampling, as the log of the mean over *m_R_* tree samples. For example, we approximate *ℓ_AC_* by

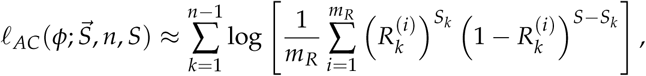

where the *m_R_* samples of 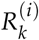 are generated as just described.
3. To estimate the Godambe information, *I_G_*(*g_q_*, *φ*), we first estimate 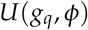 and *V*(*g_q_*, *φ*). Let 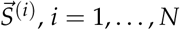, be Monte-Carlo samples from the correctly specified model with parameter *φ*_0_, and let *ℓ_q_* be the misspecified log-likelihood. Then

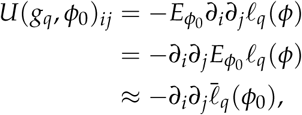

where

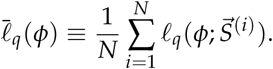

For each individual 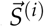, *ℓ_q_* is evaluated by Monte-Carlo integration over *m_R_* random tree samples. For large *N*, 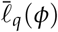 is sharply peaked and nearly quadratic near the peak. So far, we have only implemented this calculation for scalar *φ*. To estimate 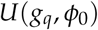, we compute 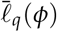for a grid of closely spaced values near *φ*_0_, fit a quadratic to 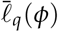 numerically, and take 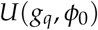 to be twice the coefficient of the quadratic term. To estimate *V*(*g_q_*, *φ*_0_), we use the formula

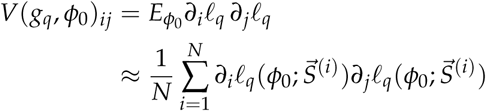

(By definition, *E_φ_g*(*X*, *φ*) = 0 for any unbiased estimating function, such as *∂_i_ℓ_q_*(*φ*; *X*), so we need not subtract off the square of the mean.^5^) Again, we have only implemented this calculation for scalar *φ*. For this case, we compute 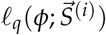 for closely spaced values near *φ*_0_, and compute *dℓ_q_*/*dφ* as the slope.
4. The method for calculating the estimate depends on whether we have a single sample of 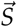 or a large number of i.i.d. samples. The likelihood for a single 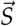 is not sharply peaked, so we fit a smoothing spline to the computed values across a broad grid, and maximize the value of the spline numerically. When we have a large sample, by contrast, the likelihood is regular and sharply peaked, as just noted. In this case, we compute values on a closely spaced grid and fit a quadratic, computing the mean as the maximum of the quadratic.

### 5.1. Computational costs

The computational demands for an estimate for a single dataset are rather small, and for problems of the size considered in this paper, satisfactory results are obtained for values around *m_R_* = 1000 to 10,000. Such calculations may be performed in a matter of minutes to hours on a laptop, depending on *n*. Most of the time is spent computing the 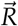 samples, which can be cached for subsequent computations using, for example, different estimators.

For estimating *σ_G_*(*g_q_*) one needs to perform *N* calculations of this form, which takes proportionally longer; also the calculations need to be more precise, because we need the slope and the quadratic curvature of the likelihood, and not just its maximum. For the data in this paper, we have used *m_R_* = 100, 000 and *N* = 5, 000, which is computationally much more demanding than the rough computation of a single maximum. The values of *σ_G_*(*g_q_*), 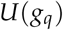, and *V*(*g_q_*) were computed for *n* = 25, 50, 100, 250, 500, and 1000, for *S* = 25, 50, 100, and 250, and for *a* values spaced at intervals of 0.05 and *ψ* values spaced at intervals of 0.01. The results are plotted in Supplement B, and estimates for other values of *n*, *S* and *a* or *ψ* may be obtained by interpolating (or extrapolating) the values given there.

All quantities are much faster to compute for the unlinked estimator 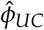 than the linked counterparts, essentially because the latter requires the computation of the moments 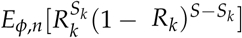, whereas the former needs only the first moment *E_φ_*_,*n*_*R_k_*, at which point *E_φ_*_,*n*_*R_k_* and 1 − *E_φ_*_,*n*_*R_k_* can be taken to the appropriate powers. This is particularly important if one is calculating 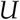 or *V*, which require Monte Carlo sampling over 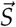 and new moment calculations for each sample. One can speed things up even further by approximating *E_φ_*_,*n*_*R_k_* by *E_φ_*_,*n*_*Z_k_* and calculating the latter using the fast algorithms of Spence et al. (2016), which do not require sampling over the coalescent tree. Of course, each of these steps involves an additional approximation, whose impact must be assessed.

### 5.2. Binning

Binning involves subdividing the set {1, …, *n* − 1} of *k*-values (or {1, …, ⌊*n*/2⌋}, if folded), into subsets. Our binning algorithm is based on the cumulative distribution function (CDF) of the expected normalized site frequency spectrum, *F*(*k*) = Σ_1≤*j*≤*k*_ *ER_k_*, *k* = 1, …, *n* – 1. Our bins are the nonempty subsets of the form

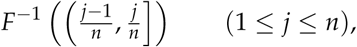

except that we always take the set {1} to be its own bin. Intuitively, we draw horizontal lines at intervals *j*/*n* across the CDF. With each pair of adjacent lines there is a set of *k*-values for which *F*(*k*) falls between the lines; the bins are the nonempty sets. For small *k*, the bins contain only a single element, but as *k* gets larger, the bins contain many elements. For example, for *n* = 50 we use the following bins:

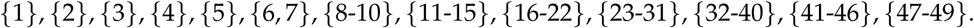

We use the same bins for all parameter values. For the beta coalescent, we compute bins assuming *a* = 1, and for EW we assume *ψ* = 0.1. In our experience, this binning procedure has no discernable effect on the estimates, and we use it for all the analyses in this paper.

### 5.3. Code availability

The routines described here have been implemented in the statistical language R (R Core Team, 2015), and will be available online.

## 6. Results

We present results using synthetic data for the symmetric beta and Eldon-Wakeley coalescents, and then apply our analysis to Atlantic Cod datasets. In presenting these results, we focus on the CI’s and *p*-values. Plots of *σ_G_*(*φ*, *n*, *S*) as a function of *n* and *S* are provided in Supplement *B*.

### 6.1. Inferences based on the entire SFS

In our first set of simulations, we study the uncertainties for inferences based on the entire SFS. The data are generated using Model A, and *φ* is estimating using 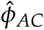.

#### 6.1.1 Confidence intervals

CI’s of confidence level 0.95 for the beta coalescent are shown in Figures 3 and 4. In Figure 3 we show the CI’s for fixed *n* and varying *S*. The most striking feature of these plots is that for fixed *n*, the CI’s are only slightly reduced by increasing *S*, even though the increase is by a factor of ten, from 25 to 250. Figure 4 provides an alternative view of the same data, and shows that, by contrast, the CI’s are sharply reduced by increasing *n* for fixed *S*. In particular, the smallest value of 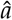 at which the Kingman coalescent is excluded, which is the value of 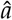 at which the lower boundary of the CI leaves the horizontal axis, decreases only slowly when decreasing *S* for fixed *n*, but rapidly when decreasing *n* for fixed *S*.

**Figure 3:**
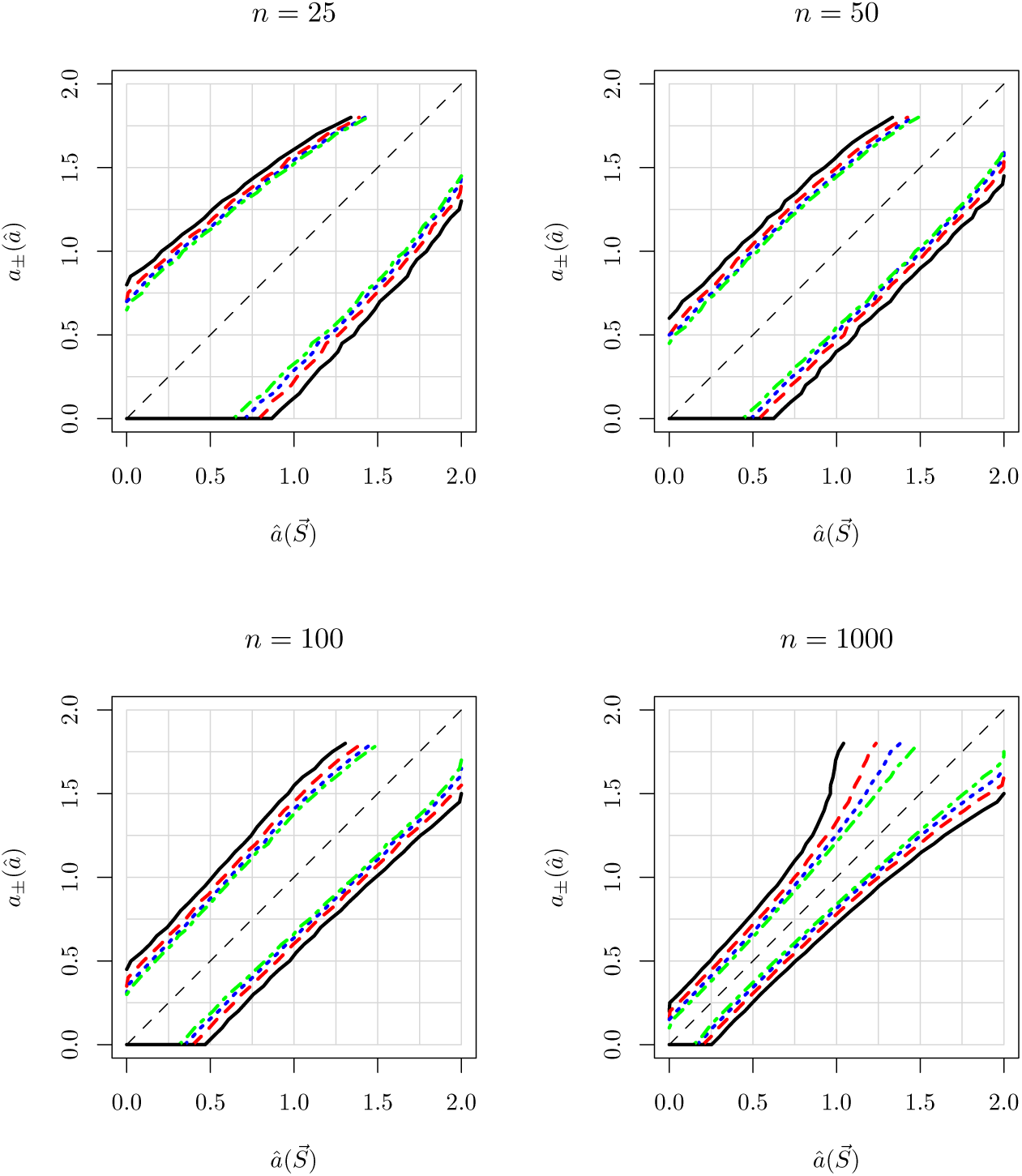
95% CI’s for the beta coalescent, Λ = *Beta*(*a*, 2 − *a*), for fixed *n* as *S* is varied. To determine the CI for (*n*, *S*) and 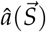, choose the graph corresponding to the value of *n*. The CI is the vertical line segment at 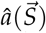, extending between the two lines with color and line type corresponding to *S*. The colors and line types are black/solid for *S* = 25, red/dashes for *S* = 50, blue/dots for *S* = 100, and green/dash-dots for *S* = 250. Note that the lines for the lower boundary of the CI overlap when they reach the horizontal axis, and only the black line is visible. Note also that the CI’s only extend vertically to values of *a* around 1.8; this reflects the fact that we only have *σ_G_*(*a*_0_, *n*, *S*) for *a*_0_ ≤ 1.8, due to the numerical difficulties of calculating *σ_G_*(*a*_0_, *n*, *S*) as *a*_0_ approaches the star endpoint. The dashed black line is the diagonal.

**Figure 4:**
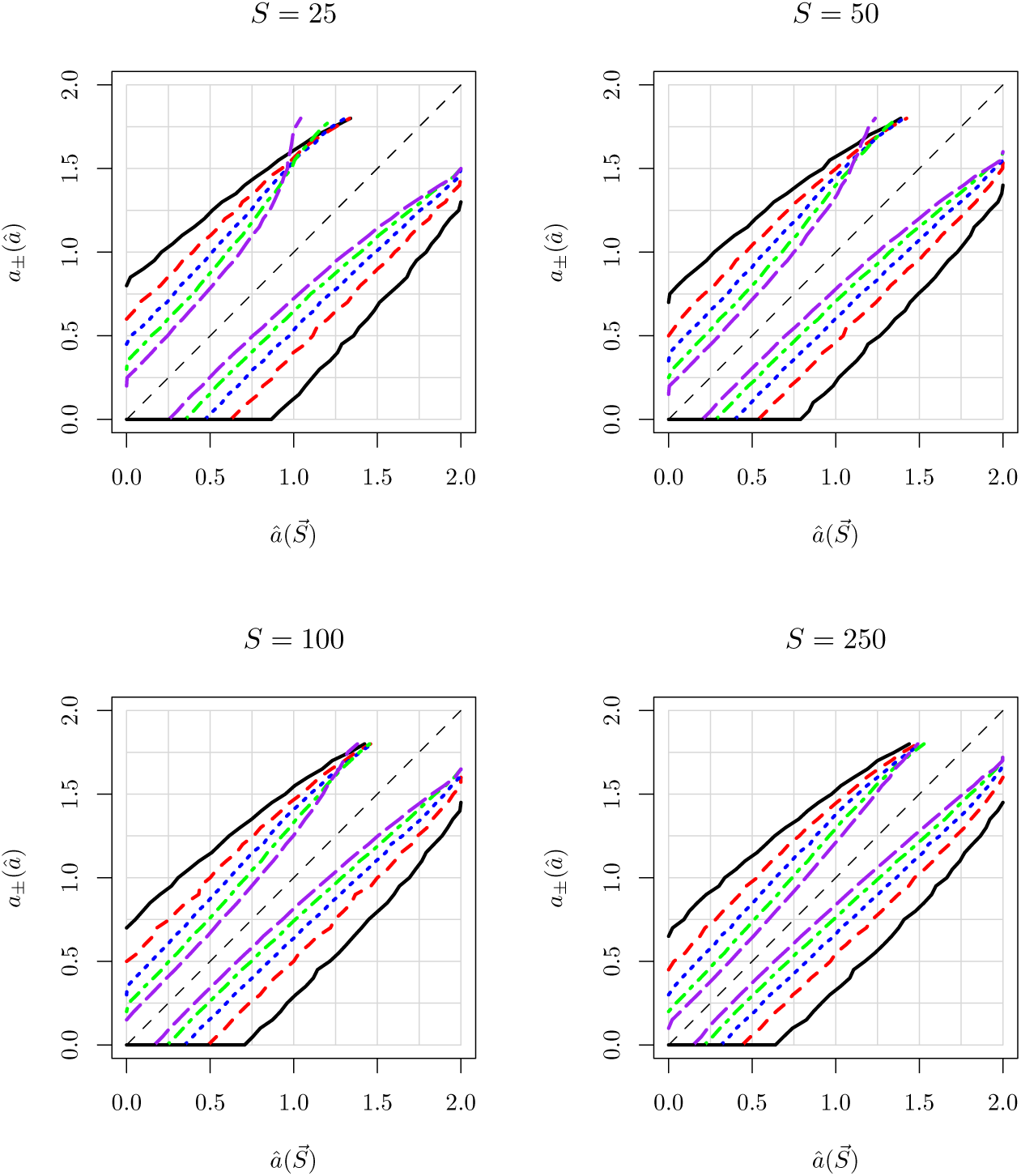
95% CI’s for the beta coalescent, for fixed *S* and varying *n*. Here the color/line type code for the value of *n*, rather than *S*. The colors and line types are black/solid for *n* = 25, red/dashes for *n* = 50, blue/dots for *n* = 100, and green/dash-dots for *n* = 250, and purple/longdashes for *n* = 1000.

The reason that the CI’s are so much more sensitive to changes in *n* than to changes in *S* is that most of the variation in the estimator 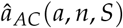 comes from variation in the coalescent tree, which is reduced by increasing *n*, but not by increasing *S*. We can see this directly by examining the asymptotic sampling variance, *σ_G_*(*a*, *n*, *S*). It is only mildly reduced by increasing *S* for fixed *n* (Figure 22), but sharply reduced by increasing *n* for fixed *S* (Figure 23).

In Figures 5 and 6, we present the analogous plots for the EW-coalescent. These figures are quite different from those for the beta coalescent, which shows that inferences from the data are strongly model-dependent. We again note that for fixed *n*, increasing *S* is of limited value, because the tree-based uncertainty provides a floor for the total uncertainty, and that increasing *n* dramatically reduces the uncertainty for small *ψ*.

**Figure 5:**
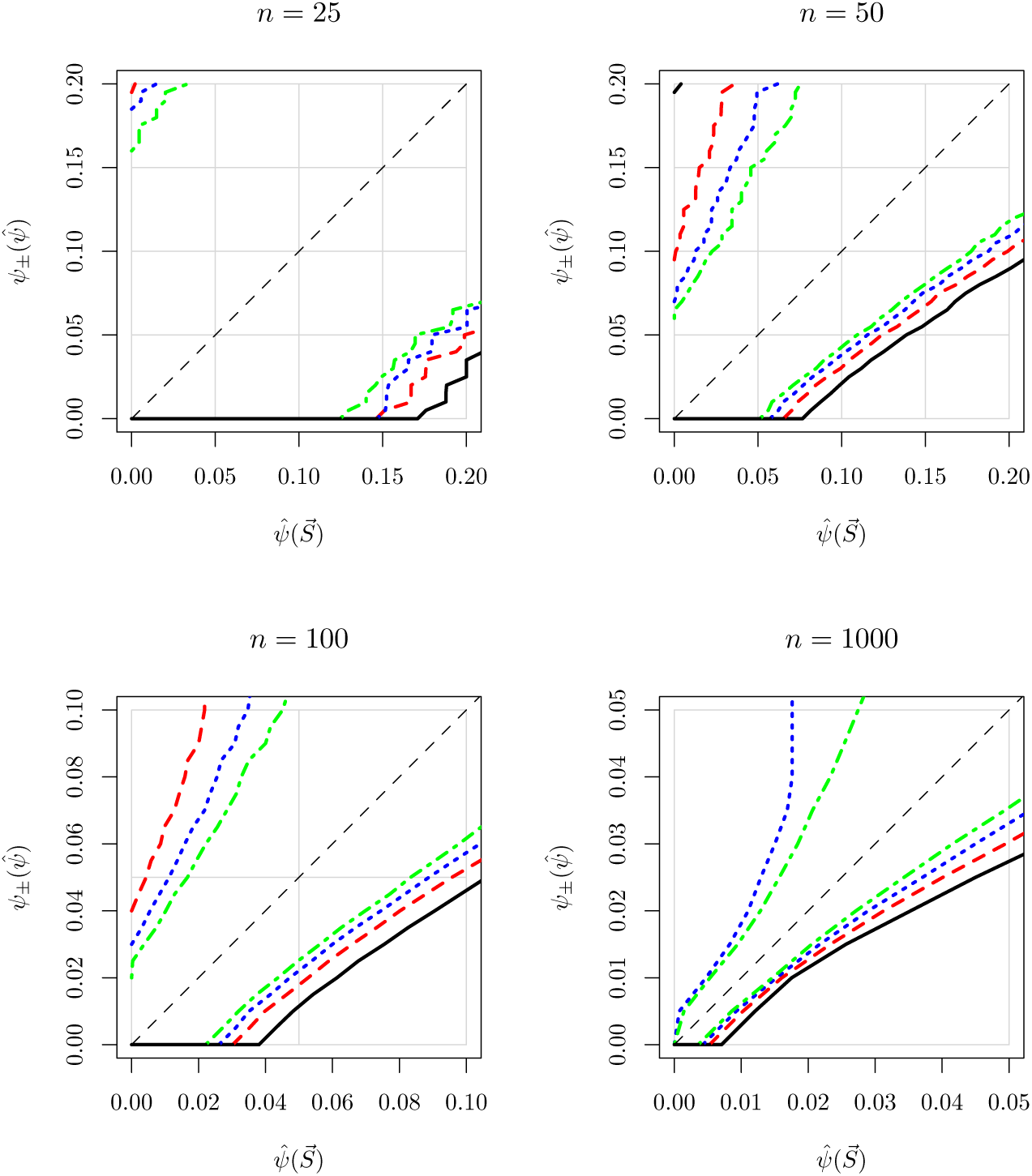
Same as Figure 3, but for EW-coalescent. Note that in the EW-coalescent, the scale of the axes changes as *S* increases.

**Figure 6:**
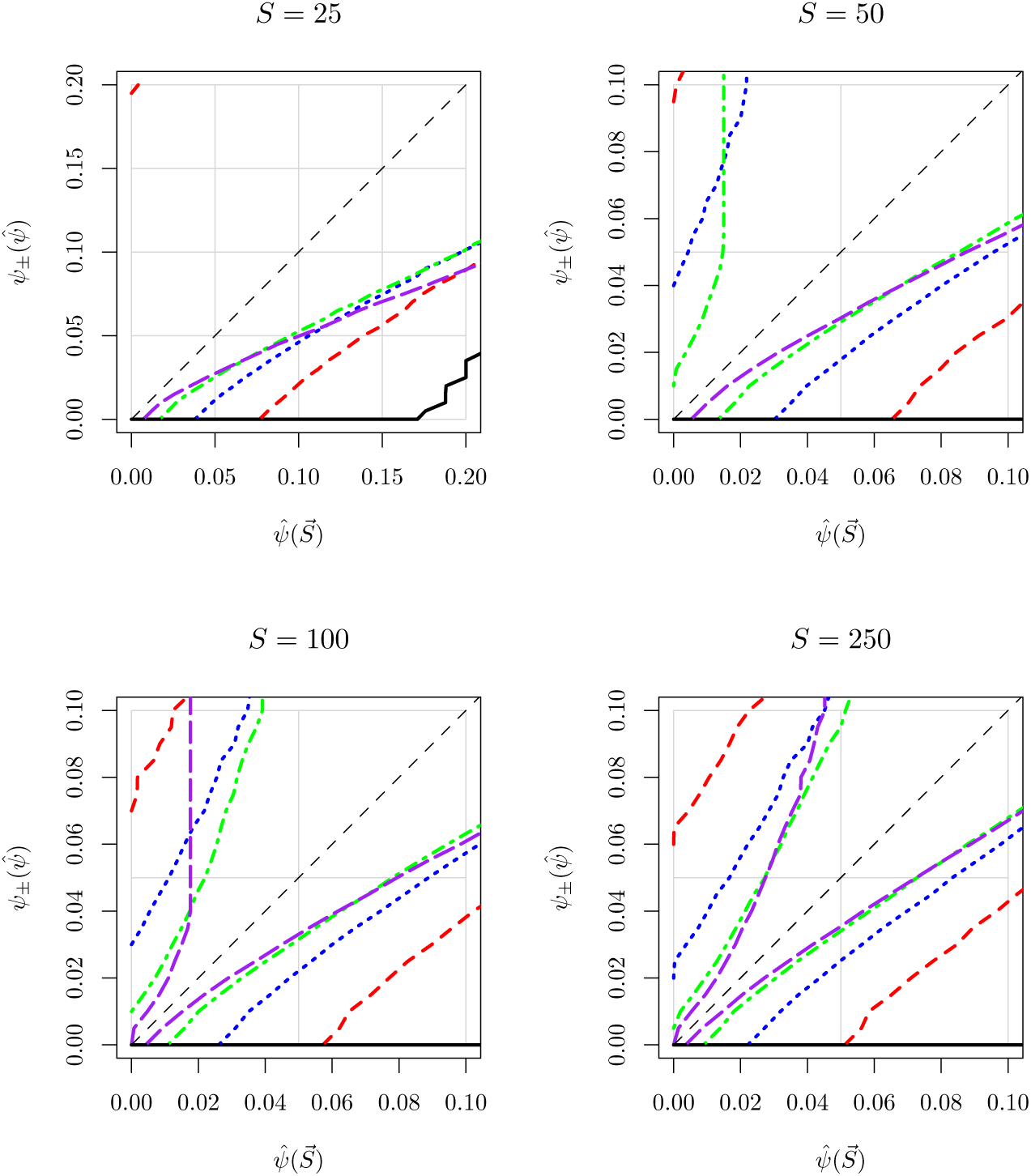
Same as Figure 4, but for EW-coalescent. The scale of the axes changes as *S* increases.

#### 6.1.2 Dependence of CI’s on *n*

One curious feature of Figure 4, for the beta coalescent, is that for values of *a* ≈ 1, the upper boundary of the CI for small *S* increases with increasing *n*; see in particular the plots for *S* = 25 and *S* = 50. The same phenomenon arises even more strongly in the EW coalescent, as apparent in Figure 6; both the lower and upper boundaries of the CI’s become looser as *n* increases. This behavior seems counterintuitive, because we expect our CI’s to get smaller as we increase *n*, which seems to correspond to taking more samples and getting more information.

The graphs are actually correct, but it is important to understand why. In our graphs we are keeping *S* fixed as *n* increases, whereas if we added more sequences, the mean value of *S* would increase in proportion to the increase in the mean total branch length. For the beta coalescent, for example, and for 0 < *a* < 1, the mean total branch length scales asymptotically as *n^a^* (Gnedin et al., 2014, Table 3). Thus, for example, in increasing *n* from 25 to 1000, we would expect *S* for *a* = 0.5 to increase by a factor of 6. These additional segregating sites would provide more information about *a*. Although we have not done the simulations, we expect that the CI’s would then become narrower with increasing *n*, in accordance with intuition.

The increase in the width of the CI with fixed *S* reflects the fact that the trees become increasingly star-like as *n* increases, particularly for larger values of *a*, which means that for fixed *S* there are fewer and fewer non-singletons to help determine the value of *a*. (In contrast, adding more sequences would never reduce the number of non-singleton sites.) Thus, for larger values of *a*, *σ_G_*(*a*, *n*, *S*) increases with increasing *n* for fixed *S*; see Figure 23 in Supplement B. The increase in *σ_G_*(*a*, *n*, *S*) increases the width of the CI.

#### 6.1.3 Ruling out the Kingman Model

In Tables 1 and 2, we give the values of 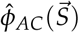 necessary for rejecting the Kingman model at a significance level of 0.05, for different values of *n* and *S*. Recall that for a given value of *n* and *S*, if the estimate 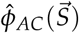 is greater than the value in the table, then the Kingman model has a *p*-value of 0.05 or less. Note that the cutoffs fall more rapidly with increasing *n* than with increasing *S*, as one would expect from the plots of *σ_G_*.

**Table 1:** Cutoffs on 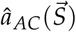 for rejecting Kingman model, beta coalescent.

| $n \backslash S$ | 25 | 50 | 100 | 250 |
| --- | --- | --- | --- | --- |
| 25 | 0.72 | 0.66 | 0.59 | 0.53 |
| 50 | 0.52 | 0.45 | 0.41 | 0.37 |
| 100 | 0.39 | 0.33 | 0.29 | 0.26 |
| 250 | 0.29 | 0.24 | 0.20 | 0.18 |
| 500 | 0.24 | 0.20 | 0.17 | 0.14 |
| 1000 | 0.21 | 0.17 | 0.14 | 0.12 |

**Table 2:**
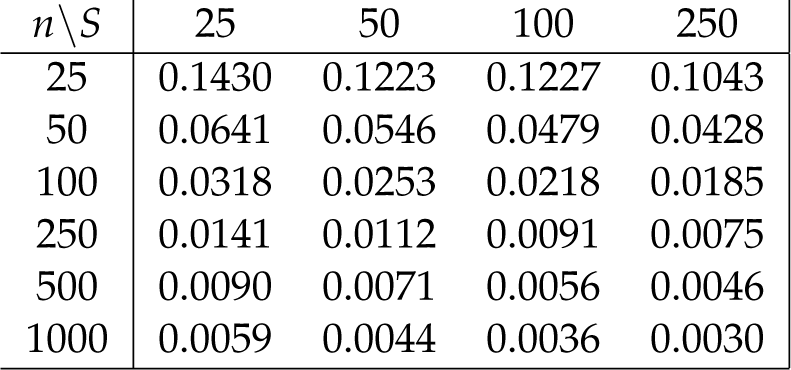
Cutoffs on 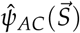 for rejecting Kingman model, EW-coalescent.

### 6.2. Inferences using only part of the data

In our second set of simulations, we use 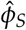 on data generated by Model A and 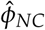 on data generated by Model N to study how much information is available when only part of the data is used.

#### 6.2.1 Singletons

Many previous analyses have based their inferences on the fraction of singletons. In Figure 7, we plot results for specific choices for *n* and *S*, but the results are similar for other values. We see that for the beta-coalescent, the CI’s for 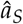 are essentially indistinguishable from those given by 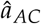. For the EW coalescent, however, the upper boundary of the CI is dramatically better with 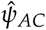 than with.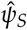.

The cutoffs for rejecting the Kingman model using 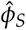 are only slightly larger than those using 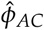; see Tables 3 and 4. (For the EW coalescent, in fact, some entries are very slightly smaller; presumably this is due to numerical errors.)

**Table 3:**
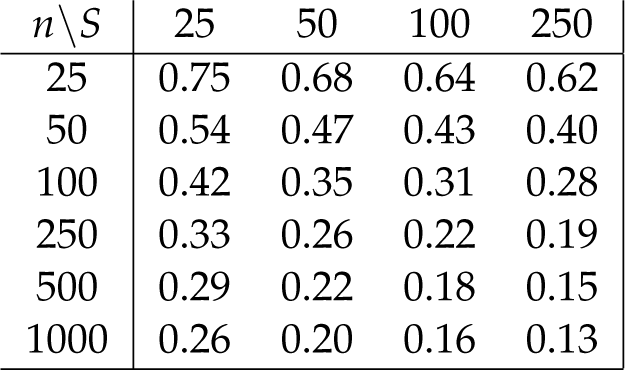
Cutoffs on 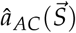 for rejecting Kingman model, beta coalescent.

**Table 4:**
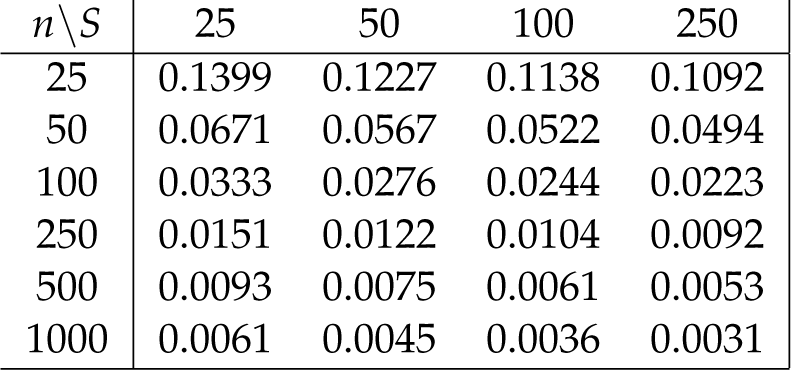
Cutoffs on 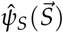 for rejecting Kingman model, EW-coalescent.

| $n \setminus S$ | 25 | 50 | 100 | 250 |
| --- | --- | --- | --- | --- |
| 25 | 0.1399 | 0.1227 | 0.1138 | 0.1092 |
| 50 | 0.0671 | 0.0567 | 0.0522 | 0.0494 |
| 100 | 0.0333 | 0.0276 | 0.0244 | 0.0223 |
| 250 | 0.0151 | 0.0122 | 0.0104 | 0.0092 |
| 500 | 0.0093 | 0.0075 | 0.0061 | 0.0053 |
| 1000 | 0.0061 | 0.0045 | 0.0036 | 0.0031 |

**Figure 7:**
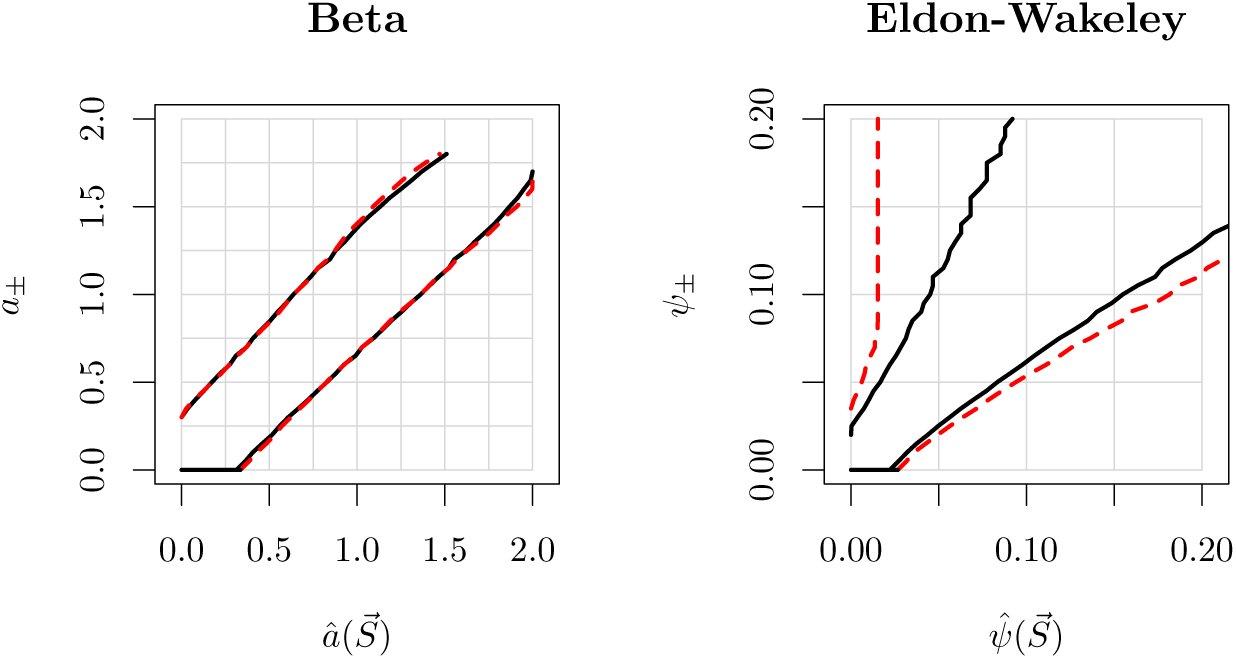
95% CI’s based on 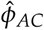 (black/solid) and 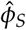 (red/dashed), for *n* = 100 and *S* = 250, for the beta and EW-coalescents.

In the case of the beta coalescent, a very simple estimate of *a*, depending only on the singletons, is

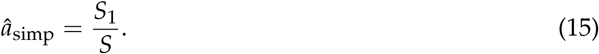

This estimate follows from the fact that for 0 < *a* < 1, *R*_1_ → *a* almost surely as *n* → ∞, and *S*_1_ ≈ *R*_1_ *S* (Berestycki et al., 2014). We have not studied the statistical properties of this estimator, and Table 3 is not directly applicable, since it pertains to 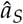. Nevertheless, if 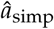 is above the cutoffs in Table 3, it is at least suggestive that the data may not be compatible with the Kingman coalescent, and that a more careful assessment may be warranted.

#### 6.2.2 Non-singletons

Although singletons provide excellent inferences for the beta coalescent, the singleton counts may be unreliable. This is because sequencing errors that affect each read independently are much more likely to affect the count of singletons than the other counts (Achaz, 2008). If we throw out the singletons, is there enough information in the non-singletons to make useful inferences?

In Figure 8, we show beta coalescent CI’s for data from Model N using 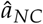. To make useful inferences, *n* and preferably also *S* need to be quite large. Under these circumstances, it is possible to get a CI that excludes the Kingman value of *a* = 0, although the CI are still very wide.

**Figure 8:**
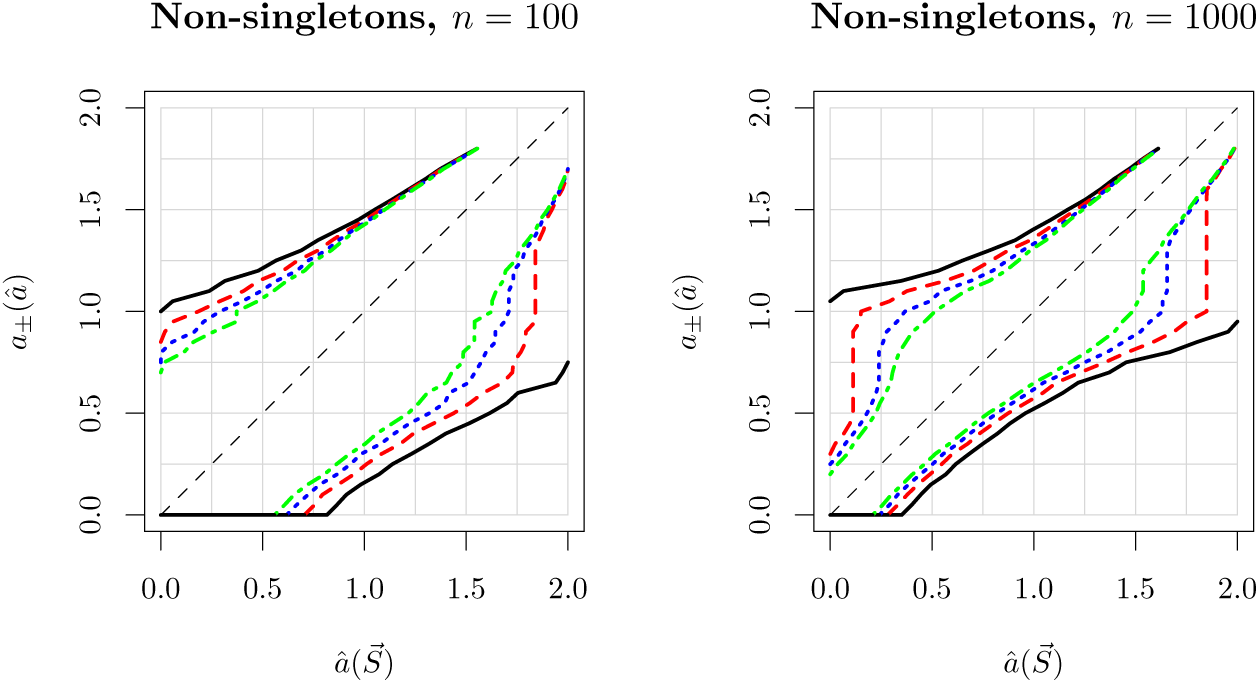
95% CI’s for the beta coalescent using 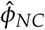 on data generated with Model N, for *n* = 100 (left) and *n* = 1000 (right), and *S*′ = 25, 50, 100, 250.

The cutoffs for rejecting the Kingman model using 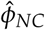 are significantly larger than those using either 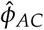 or 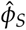; for the beta coalescent cutoffs, see Table 5. In comparing these tables, it is important to keep in mind that the uncertainties for 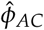 depend on *n* and *S*, whereas those for 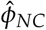 depend on *n* and *S*′, where *S*′ is the total number of nonsingletons. Even for large *S*, *S*′ may be small, which pushes us into a high uncertainty portion of the table. The values of *S*′ may be quite low, which is why we have added a column for *S*′ = 10.

**Table 5:** Cutoffs on 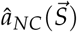 for rejecting Kingman model, beta coalescent.

| $n \setminus S'$ | 10 | 25 | 50 | 100 | 250 |
| --- | --- | --- | --- | --- | --- |
| 25 | 1.95 | 1.52 | 1.28 | 1.06 | 0.87 |
| 50 | 1.31 | 1.01 | 0.86 | 0.78 | 0.70 |
| 100 | 0.90 | 0.68 | 0.59 | 0.52 | 0.47 |
| 250 | 0.61 | 0.46 | 0.38 | 0.33 | 0.29 |
| 500 | 0.49 | 0.35 | 0.30 | 0.25 | 0.22 |
| 1000 | 0.46 | 0.29 | 0.24 | 0.20 | 0.17 |

Incidentally, the reason we use Model N to study 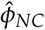, rather than Model A, is mainly pragmatic. If we used Model A the total number of non-segregating sites *S*′ would vary from sample to sample, which would complicate both the theory and the computation.

### 6.3. Inferences using the unlinked likelihood

In our final set of simulations, we study the performance of the unlinked estimator 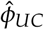 on *linked* data generated by Model A, to study how much the uncertainty estimates are degraded when we use an estimator that neglects all tree variation. By Theorem 2 in Appendix B, Model UC is an unbiased misspecification of Model A. The CI’s for 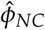 are compared with those of 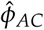 Figure 9 (beta coalescent) and Figure 10 (EW coalescent).

**Figure 9:**
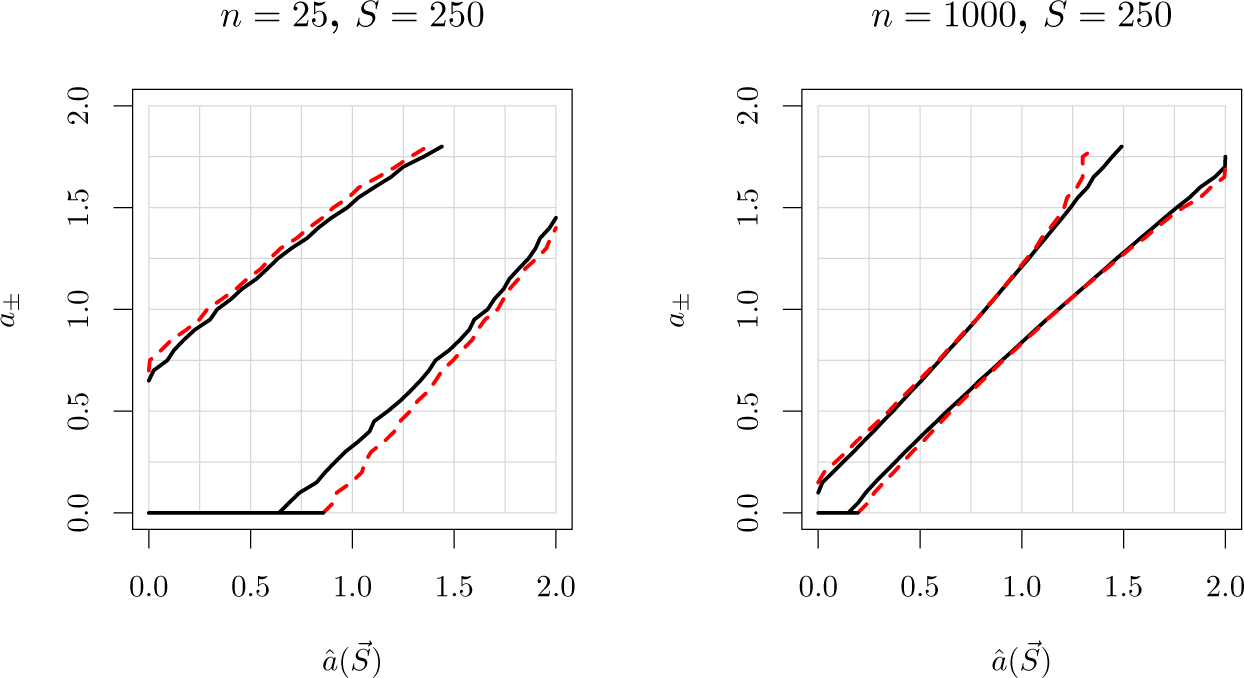
95% CI’s for the beta coalescent for *n* = 25 and *n* = 1000, with *S* = 250 in both cases. The outside (red) lines are for 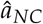, and the inside (black) lines are for 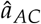.

**Figure 10:**
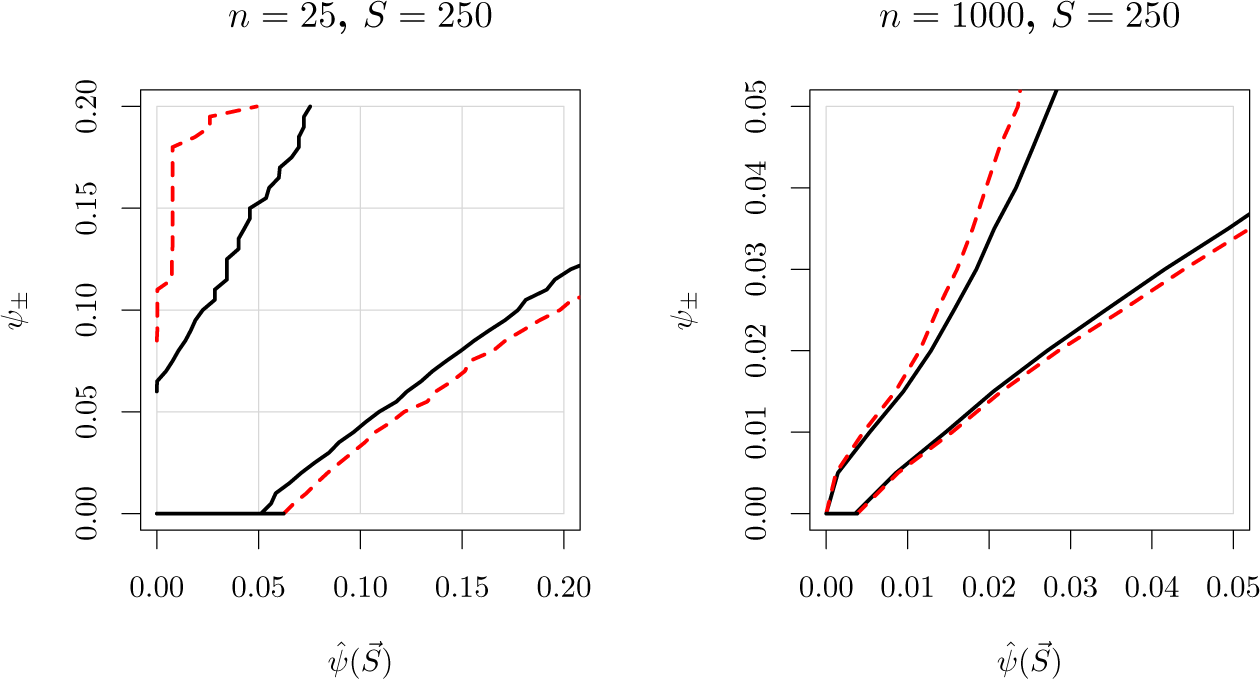
Same as Figure 9, but for EW coalescent.

Overall, the CI’s for 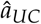 for the beta coalescent are remarkably similar to those from 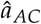, which is quite surprising, given the magnitude of the approximation. Coupled with its sharply reduced computational expense, these results suggest that 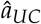 may be a good choice for many problems.

### 6.4. Unlinked data

Finally, it is instructive to evaluate uncertainties for unlinked data. In the following plots, we evaluate data generated from Model U using 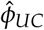. The results are shown in Figure 11 and 12, which should be compared with Figures 3 and 4, respectively. We make two observations. First, in the unlinked case, the uncertainties are significantly smaller than in the linked case, because there is no contribution from tree-based uncertainty. Second, for fixed *n* the uncertainties fall rapidly as *S* increases, in contrast to the linked case, because there is no tree-based uncertainty to provide a floor for the total uncertainty. Similar results in both cases are obtained for the EW-coalescent, as shown in Supplement C.

**Figure 11:**
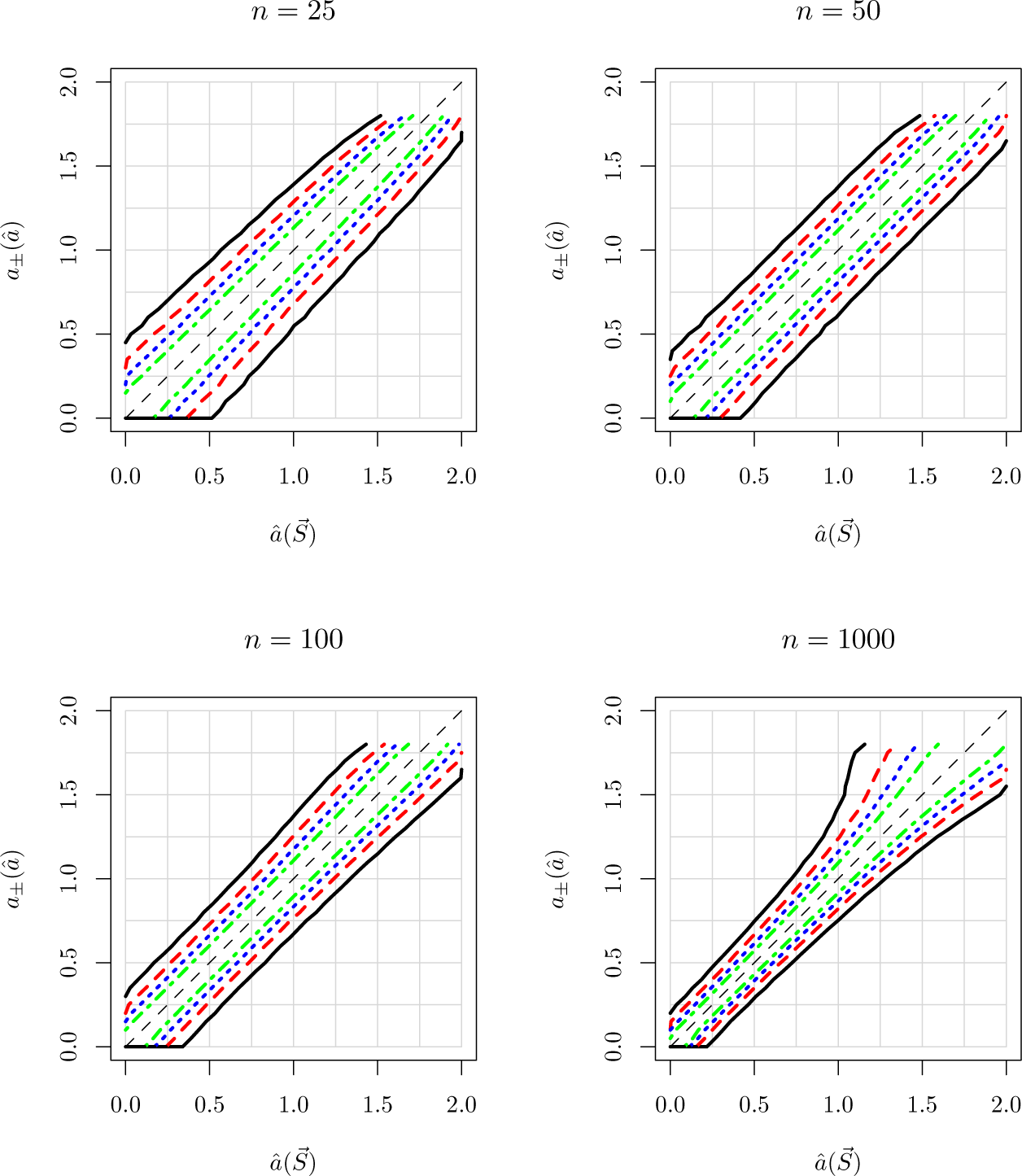
95% CI’s for beta coalescent, using 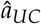 on (unlinked) data generated by Model U. Plots show CI’s as function of *S* for fixed *n*. The color/line type conventions are the same as in Figure 3.

**Figure 12:**
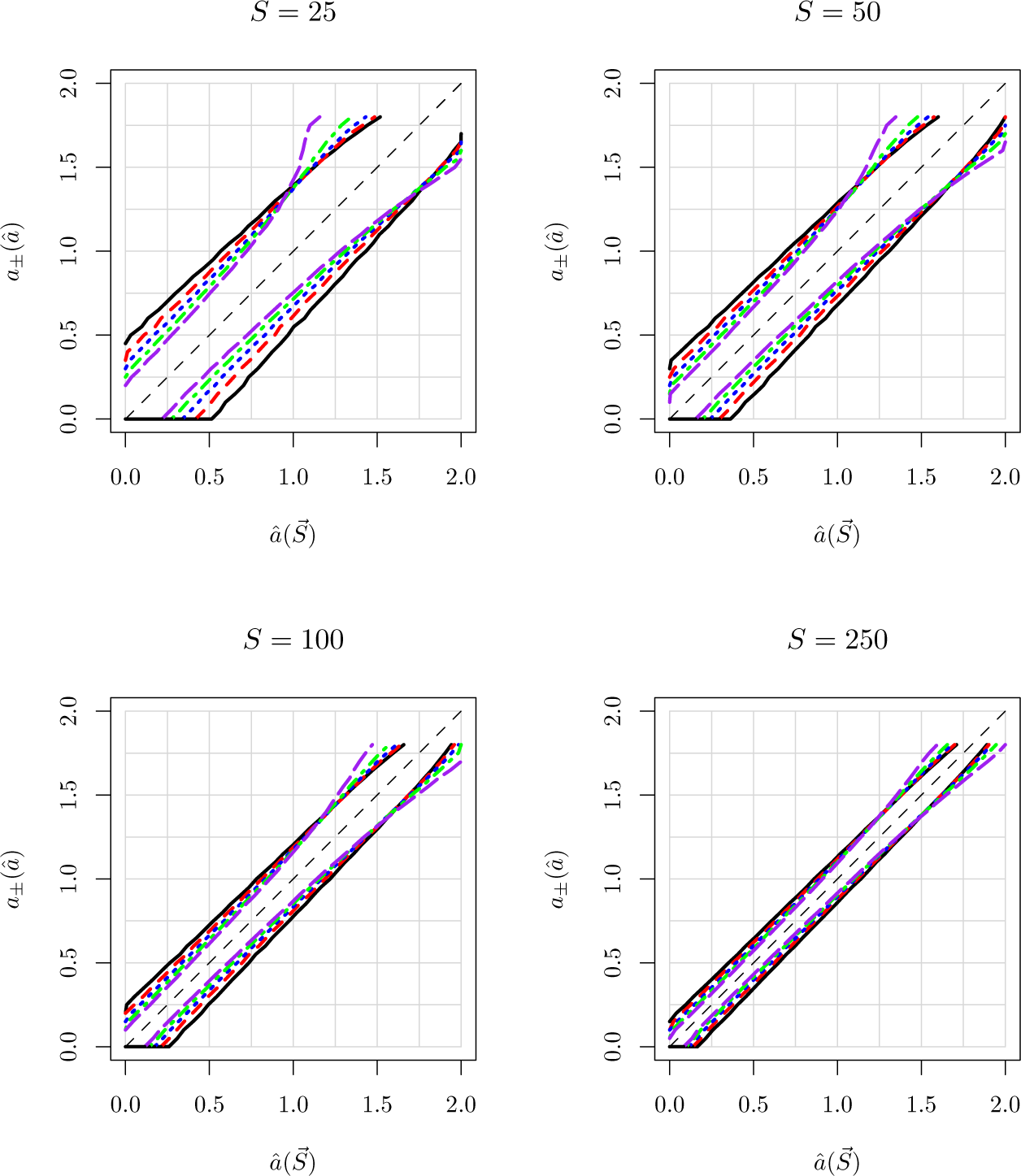
95% CI’s for beta coalescent, using 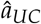 on (unlinked) data generated by Model U. Plots show CI’s as function of *S* for fixed *n*. The color/line type conventions are the same as in Figure 4.

As noted above, for linked data, the *σ_G_*(*φ*, *n*, *S*) for fixed *n* approach an asymptotic value as *S* increases, which may be identified with the tree-based uncertainty. The present analysis, in turn, isolates the mutation-based uncertainty in the absence of tree-based uncertainty. We have done preliminary analyses (data not shown) that indicate that the total uncertainty, when both types of uncertainty are included, is well-approximated by summing the tree-based and mutation-based uncertainties in quadrature.

### 6.5. How badly misspecified are our models?

As explained above, the ratio 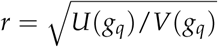 should give some indication as to the degree to which our models are misspecified. In Figure 13, we compute this ratio for *g_AC_*. There is clearly some model misspecification, although the discrepancy for *a* < 0.5 is only about 15% or less. We obtain similar results for *g_NC_*, for both the beta and EW-coalescents.

**Figure 13:**
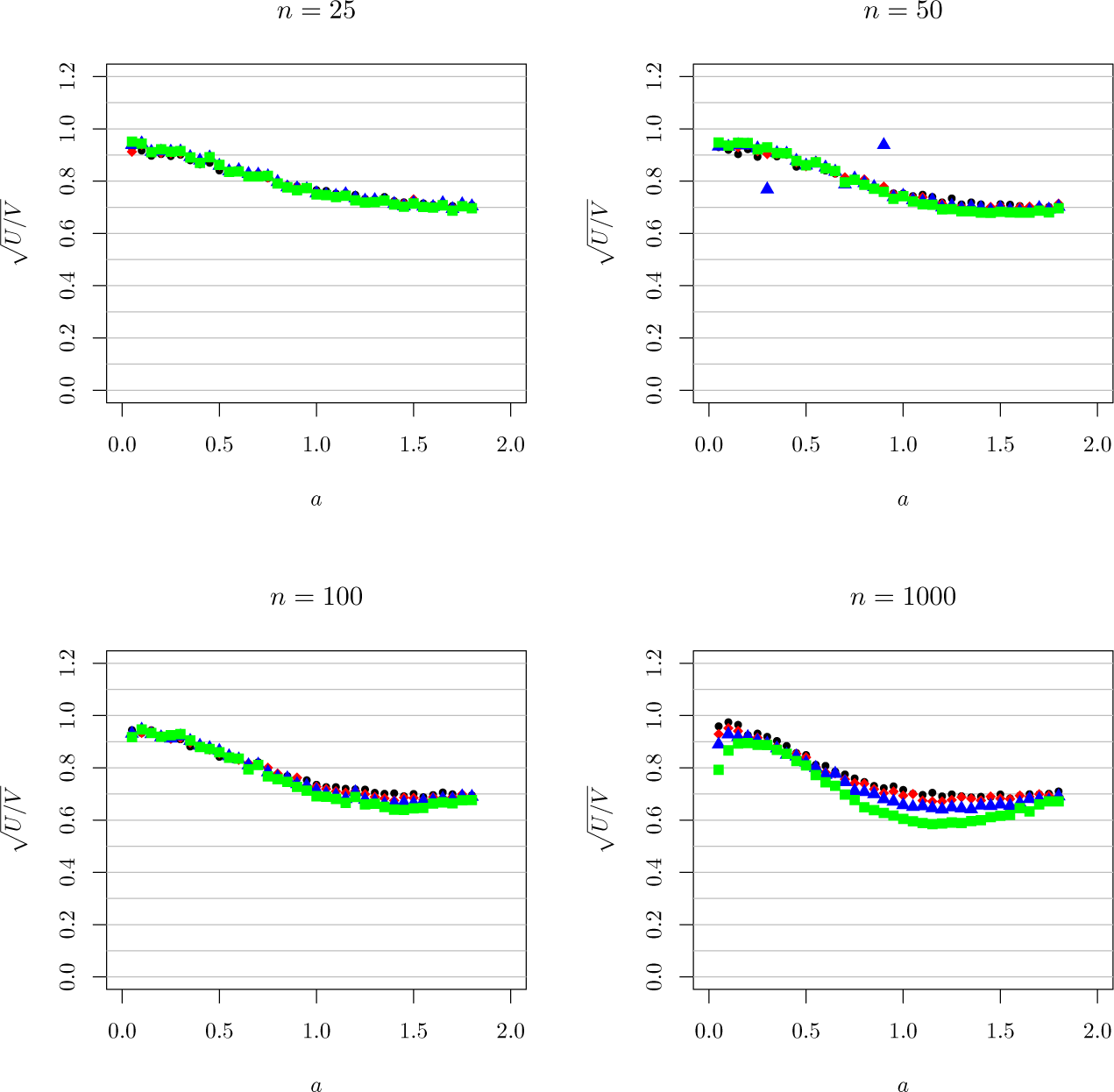
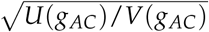 for the beta coalescent, when the true model is Model A.

In Figure 14, we plot 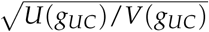 for data from Model A. As noted earlier, we expect 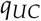 to be a dramatic misspecification of Model A, because it is not accounting for tree uncertainty. This expectation is borne out: we find 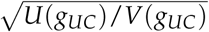 as small as 20%. Thus, even though the unlinked estimator is consistent on linked data, the naïve use of the model 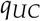 as an approximation to the correct model *p_A_* will lead to serious errors. In a Bayesian framework, for example, the naïve use of 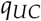 will lead to an unrealistically narrow posterior.

**Figure 14:**
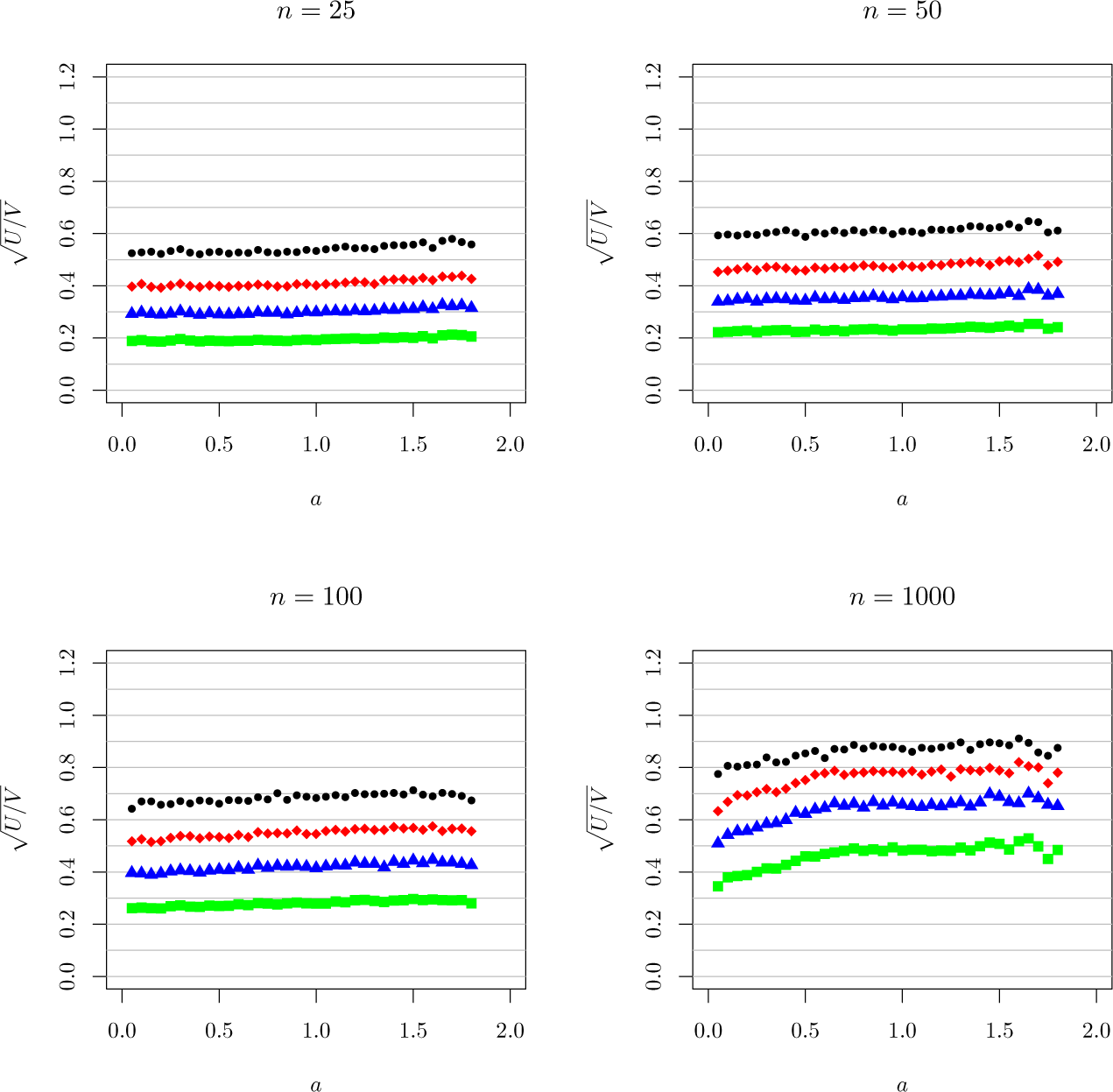
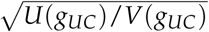 for the beta coalescent, when the true model is Model A.

### 6.6. Atlantic Cod

The Atlantic Cod *(Gadus morhua)* has been studied extensively as an organism exhibiting a broad offspring distribution, whose population dynamics might then be described by a Λ-coalescent; see Arnason et al. (2000), Arnason (2004), Arnason and Halldórsdóttir (2015), Birkner et al. (2013), Sigurgíslason and Arnason (2003), Steinrücken et al. (2013), and references therein. Focusing on three published datasets of mitochondrial sequence data, we use our methods to estimate parameters for both the beta and EW-coalescents. Since these datasets describe mitochondrial sequences, it is appropriate to assume complete linkage. These datasets all have *S* < 50, but *n* varying from 74 to 597 to 1278. Thus, they provide a good test of our finding that the uncertainty is reduced by increasing *n*. The results are given in Tables 6–8. The ℓ^2^ and pseudo-likelihood values are taken from Birkner et al. (2013); the full likelihood values are taken from Birkner et al. (2011); and the “simple” estimate is *S*_1_/*S* (Eq. 15). Values of 0.20 for the EW-coalescent reflect the fact that we did not compute *σ_G_*(*ψ*, *n*, *S*) for *ψ* > 0.20, and effectively took the parameter domain for *ψ* to be (0, 0.20).

**Table 6:** Analysis of mitochondrial sequence data from *gadus morhua*, from Sigurgíslason and Arnason (2003). 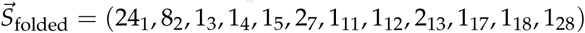, *n* = 74, *S* = 44.

| Estimator | Beta |  |  | EW |  |  |
| --- | --- | --- | --- | --- | --- | --- |
| | $\hat{a}$ | $p$-val | 95% CI | $\hat{\psi}$ | $p$-val | 95% CI |
| $\hat{\phi}_{AC}$ | 0.61 | 0.005 | (0.15, 1.09) | 0.061 | 0.004 | (0.015, 0.20) |
| $\hat{\phi}_S$ | 0.69 | 0.003 | (0.22, 1.17) | 0.086 | 0.0002 | (0.029, 0.20) |
| $\hat{\phi}_{UC}$ | 0.75 | 0.003 | (0.24, 1.25) | 0.078 | 0.001 | (0.023, 0.20) |
| $\hat{\phi}_{NC}$ | 0.44 | 0.20 | (0, 1.24) | 0 | 0.5 | (0, 0.20) |
| $\ell^2$ | 0.65 | | | 0.084 | | |
| pseudo | 0.72 |  |  | 0.06 |  |  |
| full | 0.7 |  |  |  |  |  |
| simple | 0.55 |  |  |  |  |  |

**Table 7:**
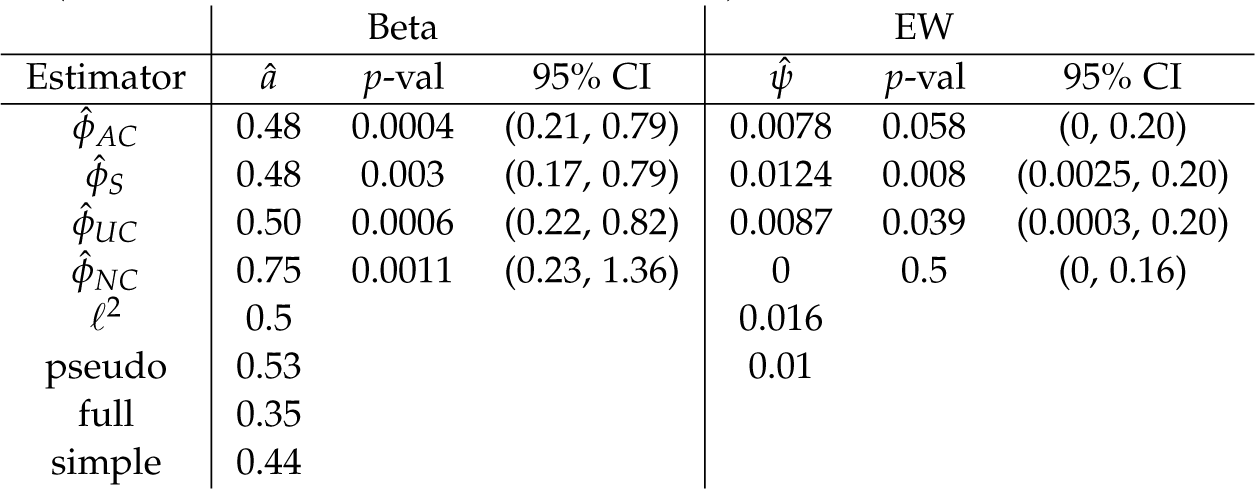
Analysis of mitochondrial sequence data from *gadus morhua*, from Arnason et al. (2000). 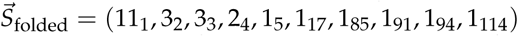, *n* = 597, *S* = 25.

**Table 8:** Analysis of mitochondrial sequence data from *gadus morhua*, from Arnason (2004). 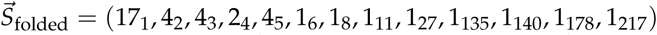, *n* = 1278, *S* = 39.

| Estimator | Beta |  |  | EW |  |  |
| --- | --- | --- | --- | --- | --- | --- |
| | $\hat{a}$ | $p\text{-val}$ | 95% CI | $\hat{\psi}$ | $p\text{-val}$ | 95% CI |
| $\hat{\phi}_{AC}$ | 0.50 | $1 \cdot 10^{-6}$ | (0.30, 0.73) | 0.0043 | 0.04 | (0.0, 0.20) |
| $\hat{\phi}_S$ | 0.47 | $1 \cdot 10^{-4}$ | (0.24, 0.70) | 0.0066 | 0.004 | (0.0009, 0.20) |
| $\hat{\phi}_{UC}$ | 0.53 | $6 \cdot 10^{-6}$ | (0.32, 0.76) | 0.0046 | 0.04 | (0.0, 0.20) |
| $\hat{\phi}_{NC}$ | 0.57 | $7 \cdot 10^{-4}$ | (0.20, 1.23) | 0.110 | $1 \cdot 10^{-17}$ | (0.05, 0.18) |
| $\ell^2$ | 0.5 | | | 0.012 | | |
| pseudo | 0.52 |  |  | 0.005 |  |  |
| simple | 0.44 |  |  |  |  |  |

We make the following observations:

- For the beta coalescent, the width of the CI’s decreases considerably with *n*, and the *p*-values also decrease consistently. The Kingman model is strongly ruled out even for *n* = 74.
- For the EW-coalescent, the width of the CI’s decreases with *n*, but 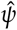 also decreases, so the confidence that *ψ* > 0 does not improve. Specifically, the *p*-values do not decrease with increasing *n*. This suggests that the model does not fit the data very well, because if a model is a good fit, then the ability to rule out an alternative model should increase with *n*. The Kingman model is sometimes ruled out by the data at the 5% level, and sometimes not.
- 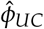 and 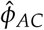 give intervals that agree closely, and comparable *p*-values as well.
- 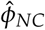 consistently gives the broadest uncertainty intervals for both models. Nevertheless, for the beta coalescent, the point estimates are consistent with those from 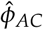 and 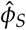. 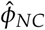 is able to exclude the Kingman model for both of the larger datasets, even though the number of non-singletons is very small.
- The behavior for the EW-coalescent under 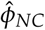 is peculiar. In the 2004 dataset, the estimate from 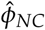 is twenty times higher than those from the other models, which suggests model misfit.

In Figures 15-17, we plot the expected SFS of the beta and EW models for the parameter values inferred from Model AC, together with the experimental data, for each of the three Atlantic cod datasets we have analyzed. Note that both of the models fit the observed singletons very well for all three datasets, but that there are frequently large discrepancies for other *k* values, although we expect large fluctuations for small *S_k_*, and it is difficult to know whether the fluctuations are reasonable without further analysis. Note also that the shapes of the EW and beta spectra are fundamentally different, with the former showing an abrupt drop between *k* = 1 and *k* = 2 and a gentler slope thereafter, and the latter showing a more linear fall-off with a steeper slope.

**Figure 15:**
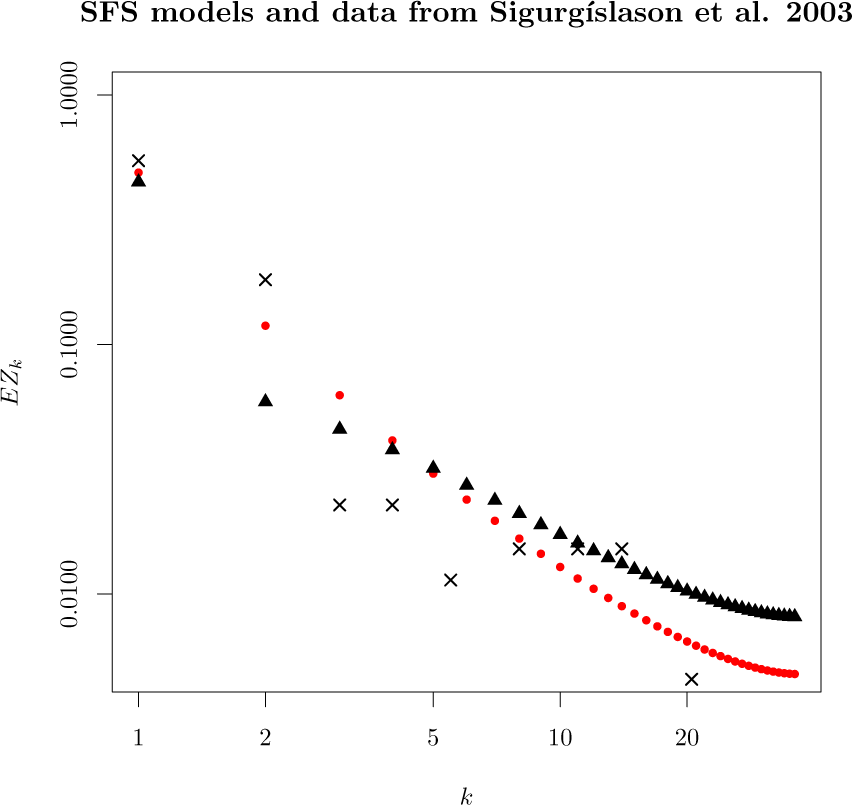
The mitochondrial sequence data from Sigurgíslason and Arnason (2003) together with the expected folded site-frequency spectrum for the data. The data are plotted in log-log coordinates. The spectrum marked with red circles is the value of 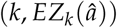, where 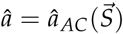. (Recall that *EZ_k_* ≡ *EL_k_*/*EL*.) The spectrum marked with black triangles is the value of 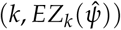, where 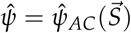. The data are the values of the (normalized) SFS, *S_k_*/*S*, which are plotted as ×’s. For higher values of *k* the data are binned, using a procedure similar to that described above, but modified to ensure that Σ*_k_*_∈_*_bin_ S_k_* > 0 for each bin. The expected SFS values are computed using the algorithm in Spence et al. (2016).

**Figure 16:**
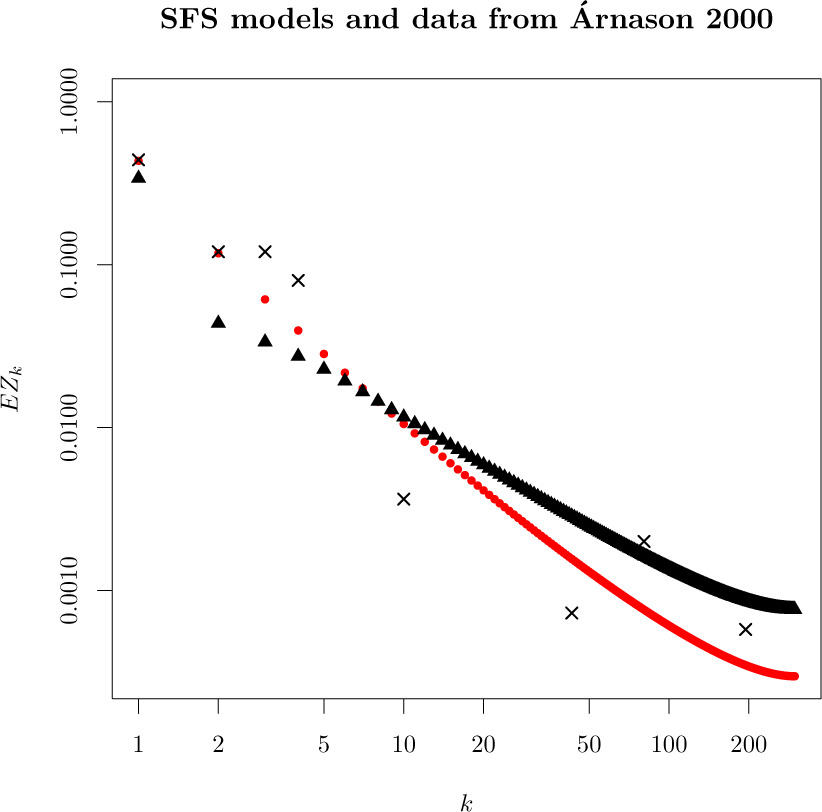
Mitochondrial sequence data from Arnason et al. (2000), plotted as in Figure 15.

**Figure 17:**
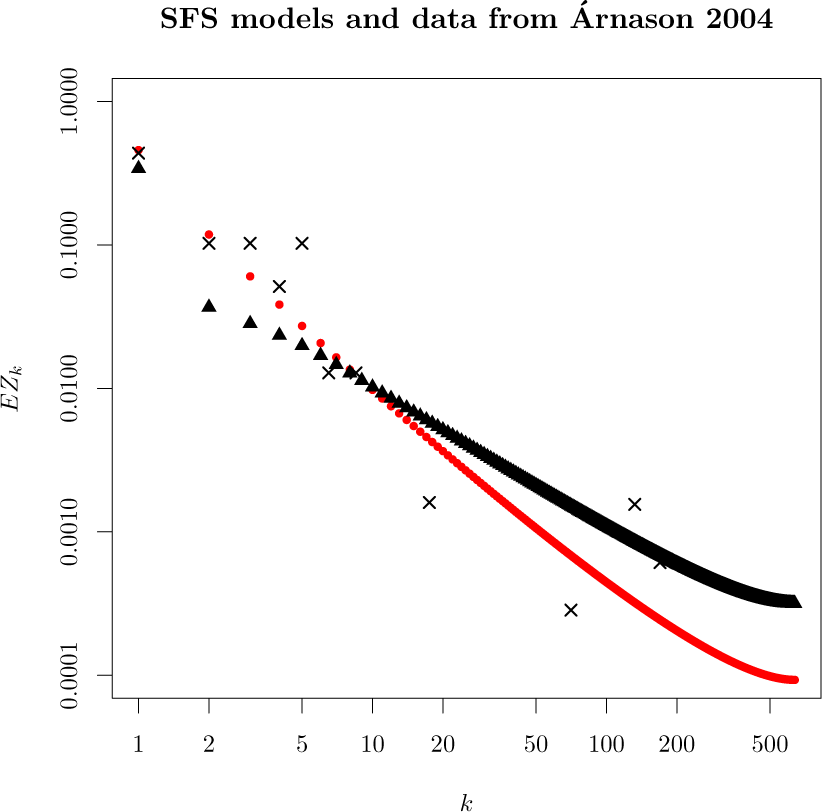
Mitochondrial sequence data from Arnason (2004), plotted as in Figure 15.

The observed singleton fraction is rather stable as *n* increases, going from 0.55 to 0.44 to 0.44. We might therefore expect that our parameter estimates would also be stable. The estimates of *a* are indeed stable as *n* increases, but the estimates of *ψ* go down by an order of magnitude. This can be understood as fitting to the singleton fraction. As noted earlier, *E_a_R*_1_ is stable as *n* increases, with a known asymptotic limit of *a*. In the EW-coalescent, by contrast, it can be shown numerically that the expected fraction of singletons increases with *n* when *n* ≳ 1/*ψ* (data not shown). To maintain the same fraction of singletons, *ψ* must be reduced to compensate, and this is why the estimates of *ψ* fall dramatically with increasing *n*. (Note that for this reason it is not possible to characterize a population with a particular value of *ψ*, since the optimal parameter depends on the sample size *n*.)

## 7. Discussion

In this paper we have used unbiased estimating functions from misspecified models to provide parameter estimates and CI’s from site frequency data, and studied the dependence of these estimates on *n* and *S*, for two different models.

We first discuss the analysis of the synthetic data. For the range of *n* and *S* values we examined, and in the parameter regimes considered, we found that in order to reduce uncertainty about the parameter of the generalized coalescent it is more useful to increase *n* than increase *S*. For example, if we have a sequencing “budget” of 6,250 nucleotides and we infer 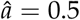, our CI is (0, 1.2) if (*n*, *S*) = (25, 250), but (0.12, 0.83) if (*n*, *S*) = (250, 25). In particular, we would be able to exclude the Kingman model in the second case, but not in the first. The same calculation for the EW-coalescent and 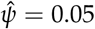 changes the CI from completely uninformative to providing a lower bound of 0.025, which again makes the difference as to whether we could rule out the Kingman model. Our guidance is specific to the range of *n*, *S*, and *a* (or *ψ*) considered, and may be different in other regimes. Nevertheless, our method provides the computational tools for assessing this tradeoff in any regime of interest.

One of our key findings is that, particularly in the beta coalescent, the use of singletons alone is often nearly as useful as the entire SFS in inferring CI’s. Although inferences have been made using only singleton data before, the validity of the approximation has not previously been studied. The near sufficiency of the singletons is encouraging, because it validates earlier results, and justifies a relatively simple approach to obtaining CI’s.

The dependence on singletons is problematic, however, if there are sequencing or data cleaning errors that might inflate or depress the number of singletons. Due to the sensitivity of the inference to the singletons, these errors might have a large effect on the resulting estimate. When this is an issue, our methods also provide an effective means of estimating parameters without using the singletons. Our results show that such inferences are possible with feasible numbers of sequences and segregating sites. For example, when *a* ~ 0.5, it is possible to rule out the Kingman coalescent with several hundred sequences, even when the number of non-singleton segregating sites is on the order of 20 or 30.

Finally, we have studied the use of an unlinked model on linked data. We have shown that when appropriately defined, 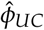 is a consistent estimator of *φ*_0_ for data generated by Model A, and that the CI’s are only mildly inflated above those of 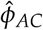 in the models we have considered. These results strongly validate the use of such an estimator.

One question we have not studied in this paper is the accuracy of our estimators for data generated by our fundamental model, Model F, in which the overall treelength varies as well as the shape. We expect that the estimators would continue to perform well, but this expectation could be tested computationally.

We now consider the issues involved in applying these results to actual genetic datasets. In calculating our parameter estimates, we have *assumed* that the coalescent model is known, i.e., that the data arises from the model for *some* parameter value. We have *not* addressed the question of how these models are chosen, or whether they fit the actual genetic data. The problem of determining appropriate models, and of testing their goodness-of-fit to the data, is essential to further progress.

In fact, the near sufficiency of the singletons is problematic for goodness-of-fit, because it suggests that the estimated parameters are not reflecting a fit of the model as a whole, but merely a fit to the observed singleton ratio. For example, our results show that within each model, there are coalescents that fit the data better than the Kingman coalescent. But this may merely reflect the fact that with the inferred parameter, the singleton ratio is closer to that of the data than predicted by the Kingman coalescent. And this is particularly worrisome if there are reasons to doubt the accuracy of the singleton count.

In this regard, it is reassuring that in the Atlantic Cod dataset, the use of singletons and non-singletons give us compatible estimates of *a* in the beta coalescent when *n* is sufficiently large. This lends some support to the hypothesis that the non-singleton data are also well-described by the beta coalescent. There are more powerful ways of assessing goodness-of-fit, which we hope to develop in subsequent work.

## Author contributions

The project was conceived jointly by TB, JFW, and TCW. The paper was written by TCW, with input from TB and JFW. The idea of using composite likelihood is due to JFW, and the generalization to unbiased estimating equations is due to TCW, as are the results in Appendix B. The modeling and theory development for generalized coalescents was carried out by TCW, with input from TB and JFW. TCW wrote the code and performed the simulations.

## Acknowledgments

We gratefully acknowledge support from the U. S. Department of Energy through the LANL Laboratory Directed Research Development Program, and from the U. S. Department of Energy under contract DE-AC52-06NA25396. Computations were performed using the Darwin Computational Cluster at Los Alamos National Laboratory.

## Appendix A: Derivation of Multinomial Model

In the main body of the paper we showed how the multinomial Model A, which involves only the ratios of branch lengths, can be derived from the Poisson Model F by assuming a modification of the experimental procedure. Although it is plausible that such a modification would not greatly affect our estimate of *φ*, it is also useful to consider how we might derive Model A directly from Model F.

Note that Model F can be written

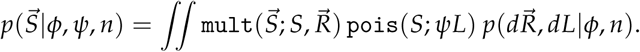

The multinomial gives one when summed over 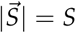, so

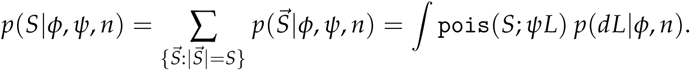

Thus,

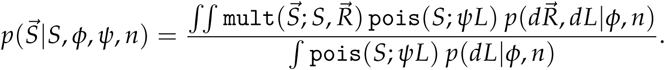

If we now assume that the shape and the size of the tree are independent,

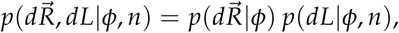

which is often probably a fairly good approximation, the integral in the numerator factors and the Poisson terms cancel, and we are left with

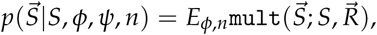

in which the *ψ*-dependence has disappeared. This is Model A.

If we use Model F in a Bayesian context and marginalize over *dψ* using the “noninformative” Jeffreys prior (Jaynes, 2003, Jeffreys, 1946), *p*(*dψ*) = *dψ*/*ψ*, then we recover the multinomial Model A without ever having to make an independence assumption. Indeed,

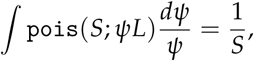

so

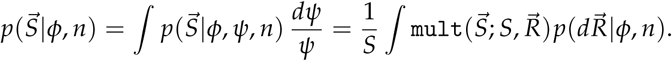

The model has a logarithmic divergence, which it inherits from the improper Jeffreys’ prior. If we (formally) condition on *S*, noting that *p*(*S*|*φ*) = 1/*S* because the multinomial sums to one, we find that

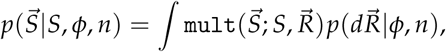

which again is Model A. Although we have adopted a frequentist approach in this paper, the last result provides the basis for an alternative Bayesian analysis.

## Appendix B: Consistency of unlinked likelihood

### Theorem 1

*Let* 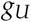 *be the score function for Model* 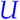. *Then* 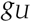 *is an unbiased estimating function for Model A.*

*Proof*. We need to show that

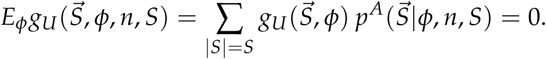

From Eq. U, 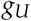 is given by

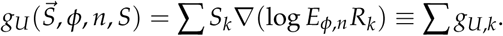

Consider the *k*th term. We have

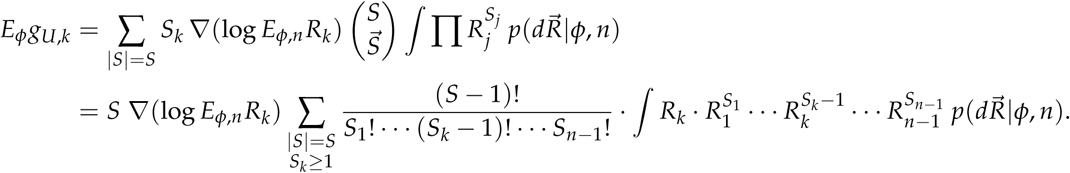

But

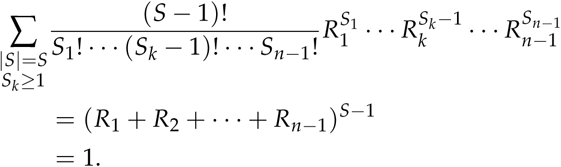

Therefore

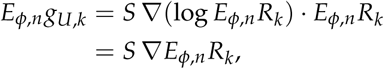

and

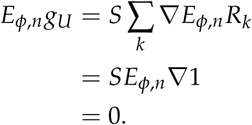

Note that we do *not* have 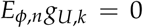 for each *k*, so that the individual terms cannot be marginal or conditional distributions, and the unlinked likelihood is not a composite likelihood for the linked model.

### Theorem 2

*Let* 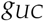 *be the score function for Model* 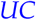. *Then* 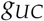 *is an unbiased estimating function for Model A.*

*Proof*. We need to show that

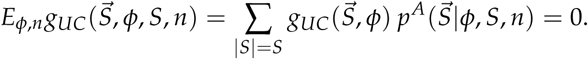

From Eq. UC, 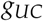 is given by

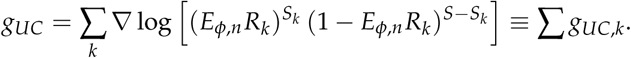

Consider the *k*th term. The structure of 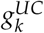 is actually the same as that of the sum in Theorem 1, so it follows immediately that 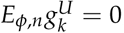, but we also provide a direct proof. We have

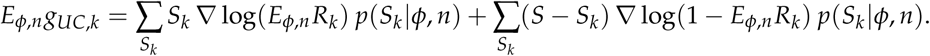

The first term is

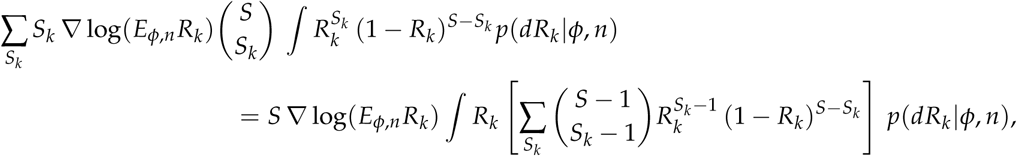

which is just *S* ∇*E_φ_*_,*n*_*R_k_*. A similar derivation gives *S* ∇(1 − *E_φ_*_,*n*_*R_k_*) for the second term. The sum is

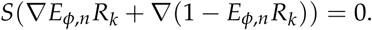

Since this is true for each *k*, the expectation of the sum is also zero.

Note that in Model UC the expectations of the individual terms are indeed zero. Nevertheless, it is not obvious how they could be interpreted in the context of composite likelihood, as the score functions of marginal or conditional distributions of Model A. Thus, it appears that we still need the broader setting of unbiased estimating functions.

## Appendix C: *p*-values and confidence intervals

We address technical issues involved in the definition of *p*-values and confidence intervals.

### Use of asymptotic distributions

In computing the distribution of 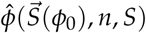, we assume that the distribution is Gaussian with mean *φ*_0_ and variance 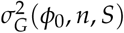. This latter quantity, however, is not the variance of the estimator for a single sample, but the asymptotic variance, which is defined by the formula

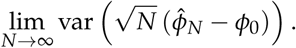

For a single sample, the distribution need not be Gaussian, particularly if *φ*_0_ is near a boundary, which will lead to a point mass on the boundary, and the variance may not be equal to *σ_G_*. The approximation will improve as the distribution becomes more concentrated, which will occur as *n* and *S* are increased.

The asymptotic distribution will be Gaussian if *φ*_0_ is in the interior of the parameter domain. If *φ*_0_ is on the boundary, however, even the asymptotic distribution will not be Gaussian. This is an important special case, because one of our objectives is to provide tests for ruling out the Kingman coalescent, and the parameter value for the Kingman coalescent lies on the boundary in both of the genealogical models we consider. The limiting forms of the sampling distribution of the estimator for a boundary parameter have been worked out for the usual ML-estimator (Moran, 1971, Self and Liang, 1987), but these results do not appear to have been extended to estimators obtained from unbiased estimating equations. We will assume that the limiting forms for the usual ML-estimators are valid with *σ_G_* replacing *σ_F_*.

For *φ*_0_ = 0, which is on the boundary of the parameter space, the asymptotic distribution of the ML-estimator is the sum of a *δ* measure of mass one-half at *φ* = 0 and a half-Gaussian for *φ* > 0; see Self and Liang (1987), Theorem 2, or Moran (1971) for more details and extensions to higher dimensions. Substituting *σ_G_* for *σ_F_*, and we get the formula

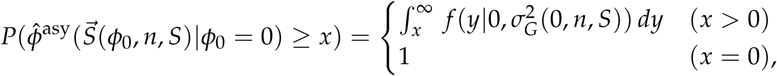

where *f*(·|*μ*, *σ*^2^) is the density of *N*(*μ*, *σ*^2^), the normal distribution with mean *μ* and variance *σ*^2^. A *p*-value of *γ* is attained when

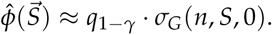

(Here, *q_α_* is the quantile function for the standard normal, *N*(0,1). I.e., if *X* ~ *N*(0,1), then *P*(*X* < *q_α_*) = *α*. For *γ* = 0.05, for example, we use *q*_0.95_ = 1.64.)

### Confidence Intervals

We need to define 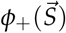 and 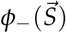 so that

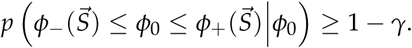

To satisfy this condition, it is sufficient that, for all *φ*_0_,

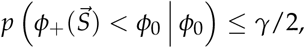

and

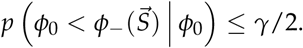

These two conditions correspond to the cases that the CI is entirely to the left, or right, of *φ*_0_.

To define 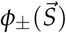, let

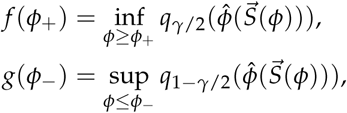

where 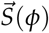 is a random sample from 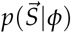, and *q_α_*(*X*) = *F*^−1^(*α*), where *F* is the distribution function of *X*. (When the argument *X* is omitted, the standard normal is assumed.) By construction, the right hand sides are monotonically increasing functions of 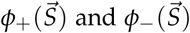: if 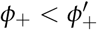, then 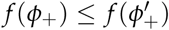, and similarly for *g*. Let

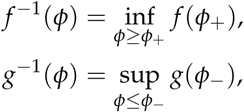

and define

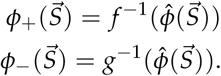

With these definitions, it follows that

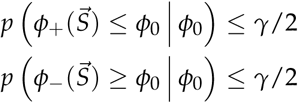

for all *φ*_0_. For example, if 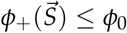, then 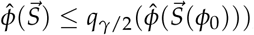, which is an event of probability at most *γ*/2. An analogous argument holds in case 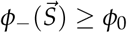. Note that with these definitions, *φ*_+_ and *φ*_−_ depend on 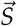 only through 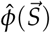.

In this paper, we approximate the estimators with their asymptotic distributions:

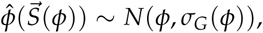

truncated at the domain boundaries. Then

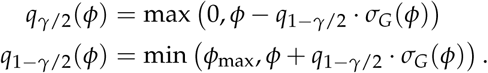

In computing *φ*_+_ and *φ*_−_, we were not able to take the supremum over all values of *φ*, because we were not able to evaluate *σ_G_* near the star endpoints. In computing the CI’s, therefore, we limited ourselves to values in (0, 1.80) for the symmetric beta coalescent and (0, 0.20) for the E-W coalescent. In effect, we assume that the parameter space is restricted to this smaller interval.

## Supplement A: Random trees, beta coalescent

**Figure 18:**
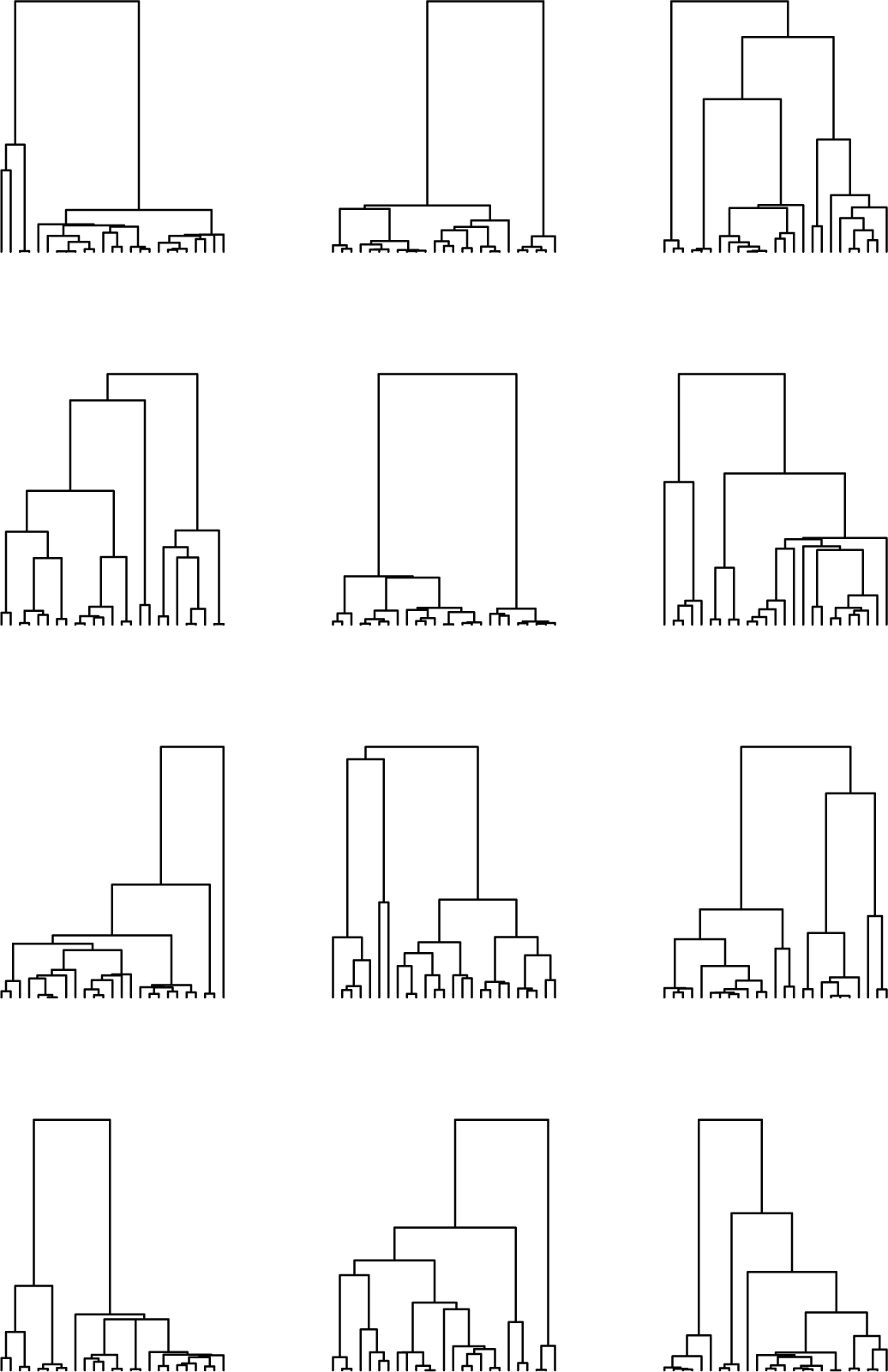
Twelve random trees with *n* = 25 for *a* = 0 (Kingman model). The height of the trees varies considerably; we have normalized them to constant height for ease of presentation.

**Figure 19:**
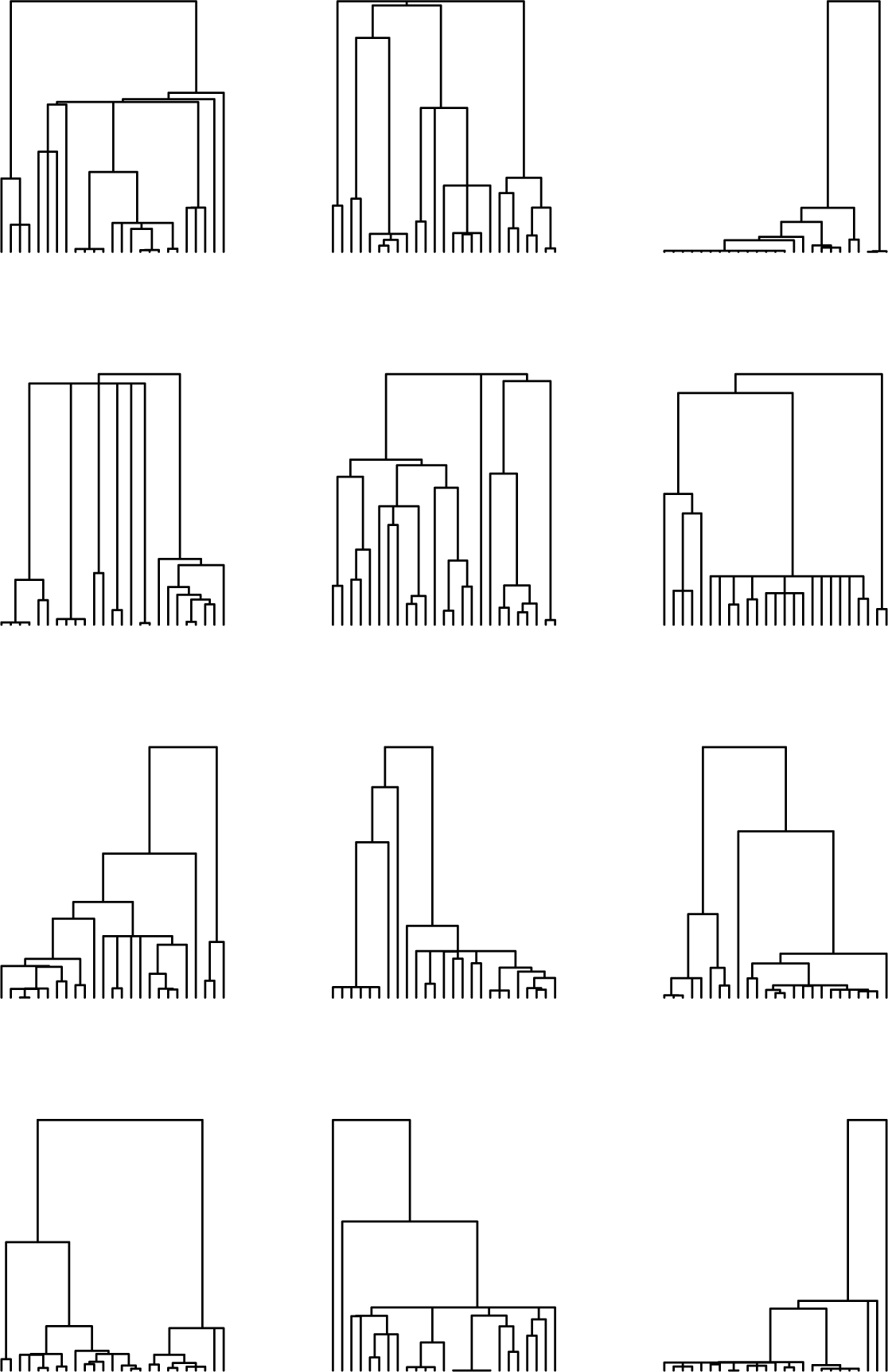
Twelve random trees with *n* = 25 for *a* = 0.5, normalized to constant height.

**Figure 20:**
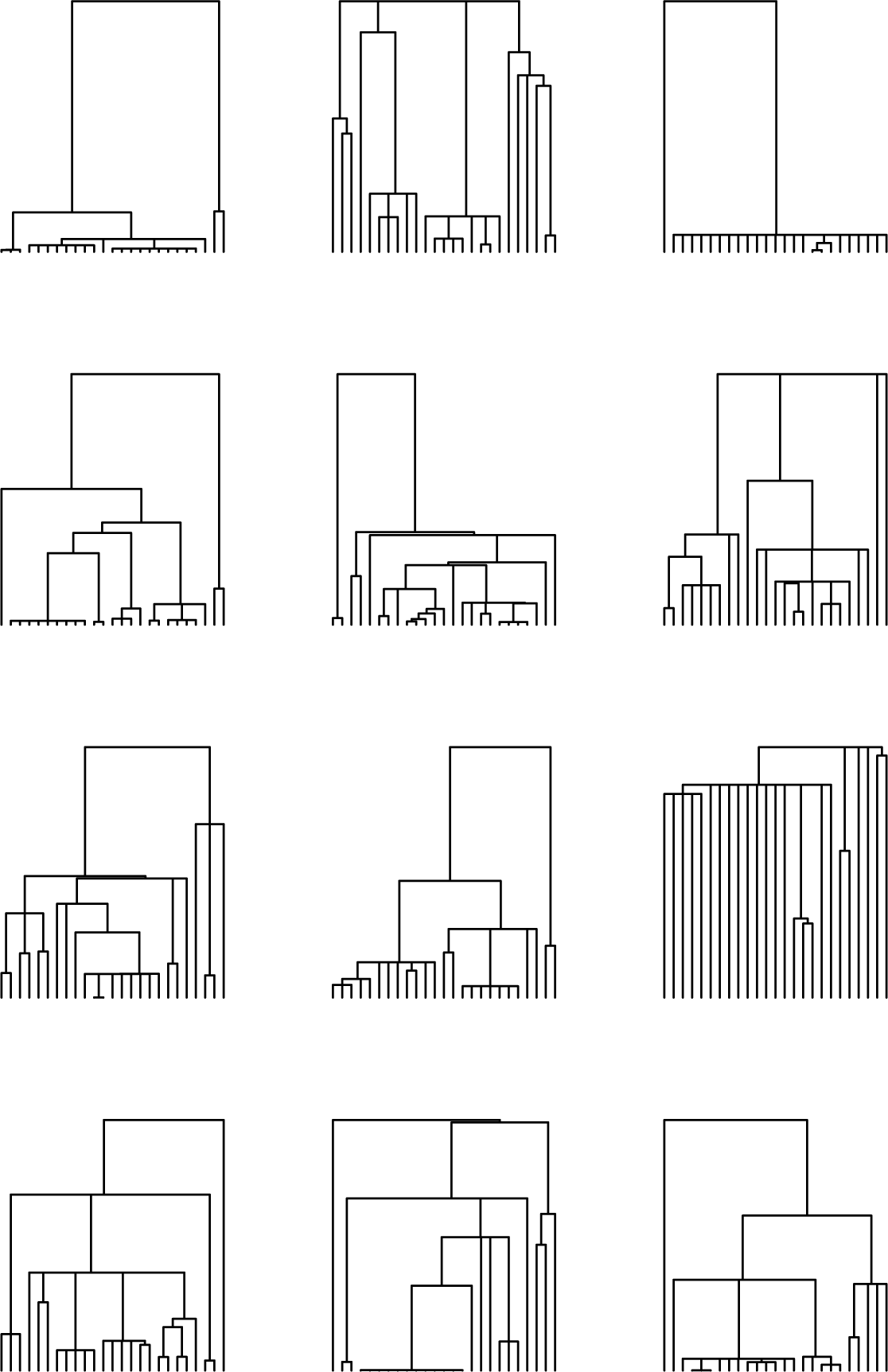
Twelve random trees with *n* = 25 for *a* = 1.0, normalized to constant height.

**Figure 21:**
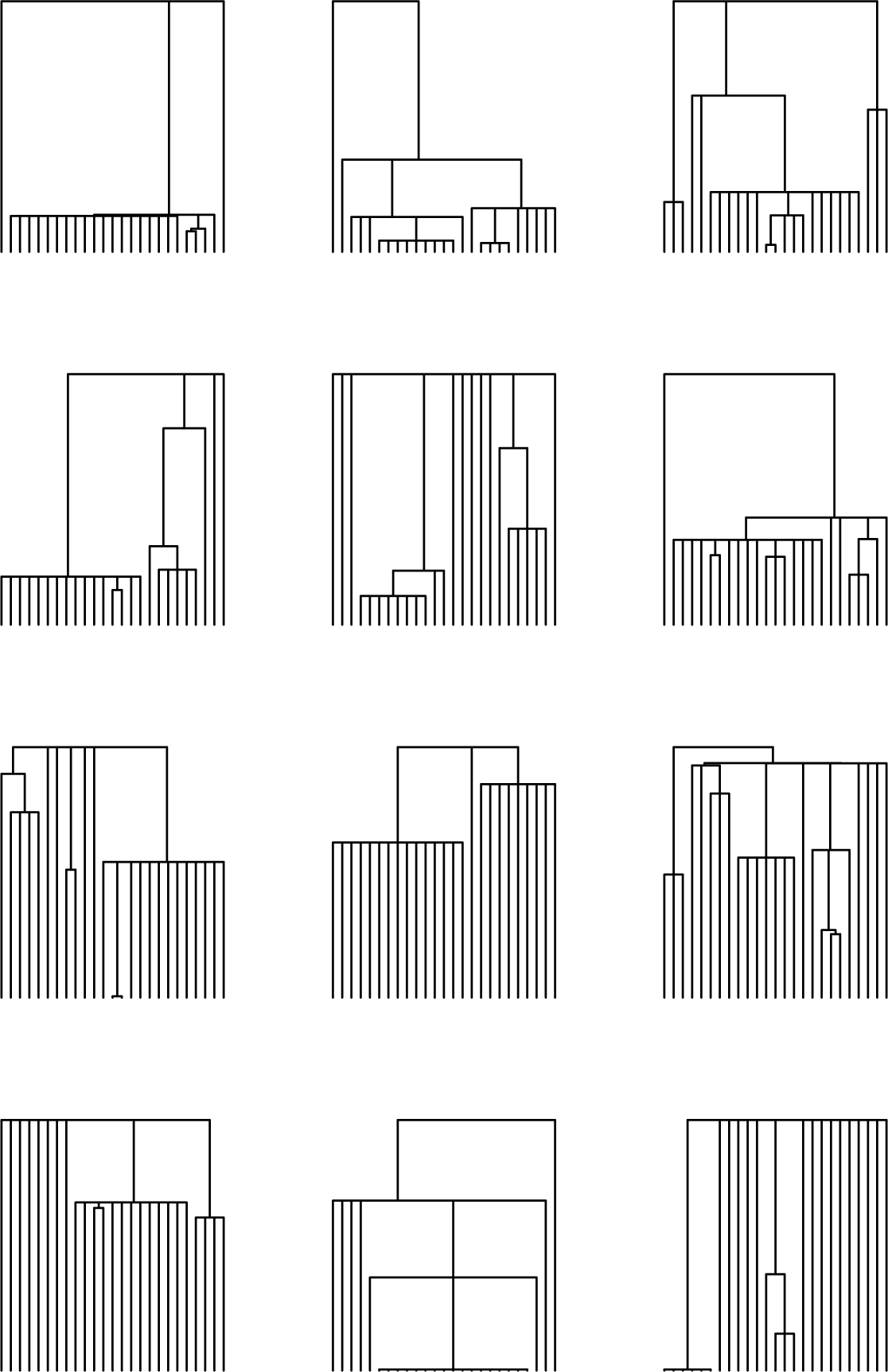
Twelve random trees with *n* = 25 for *a* = 1.5, normalized to constant height.

## Supplement B: Uncertainty plots

### Beta model

**Figure 22:**
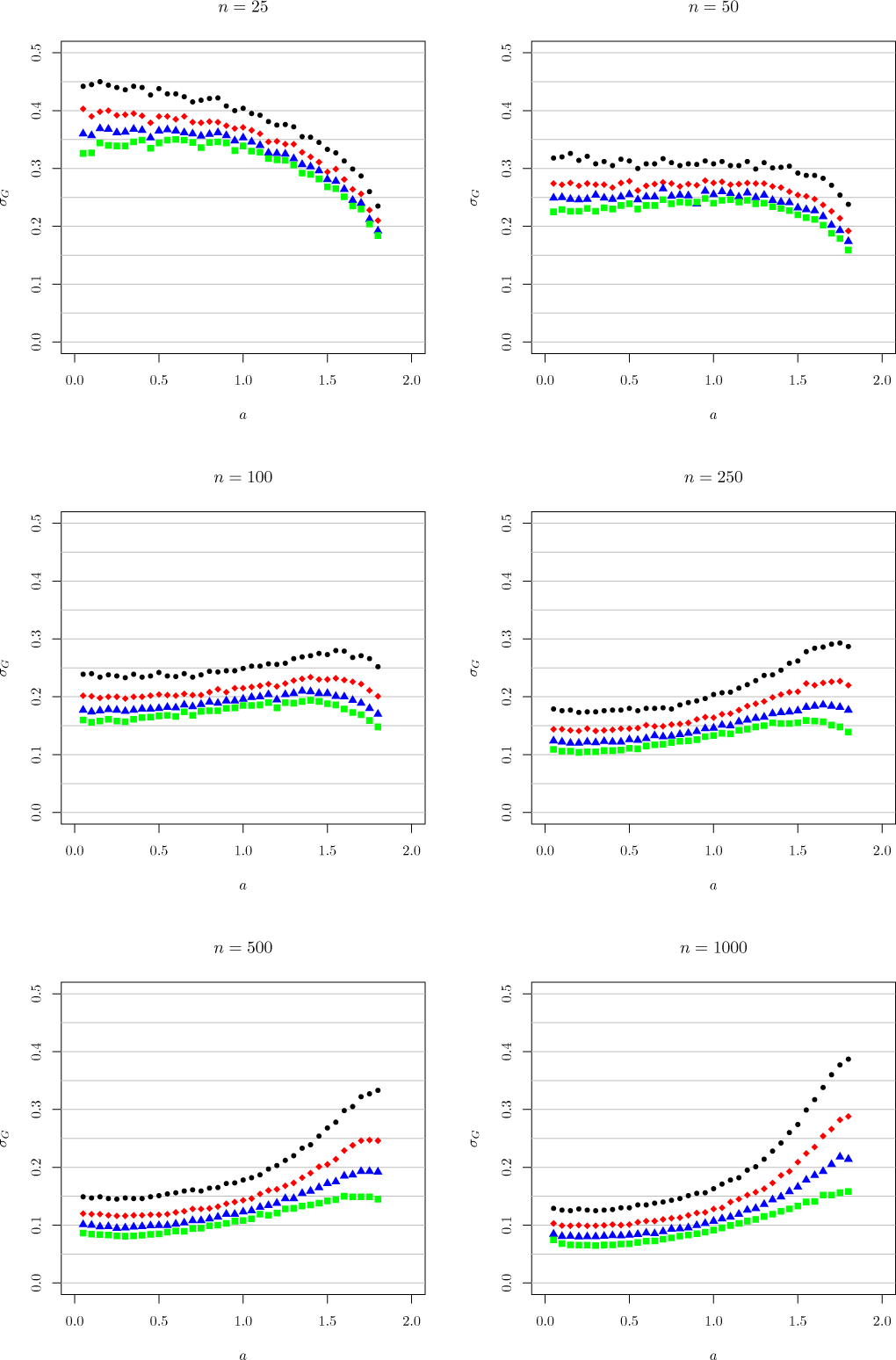
The asymptotic sampling standard deviation, 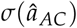, for data from 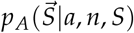 (Model A). Each plot is for fixed *n*, and shows how *σ* varies with *a* for several values of *S*. The data is given as black dots for *S* = 25, red diamonds for *S* = 50, blue triangles for *S* = 100, and green squares for *S* = 250. *σ* is inferred by numerical estimates of the inverse Godambe information, based on simulations using *m_R_* = 100, 000 and *N* = 5, 000.

**Figure 23:**
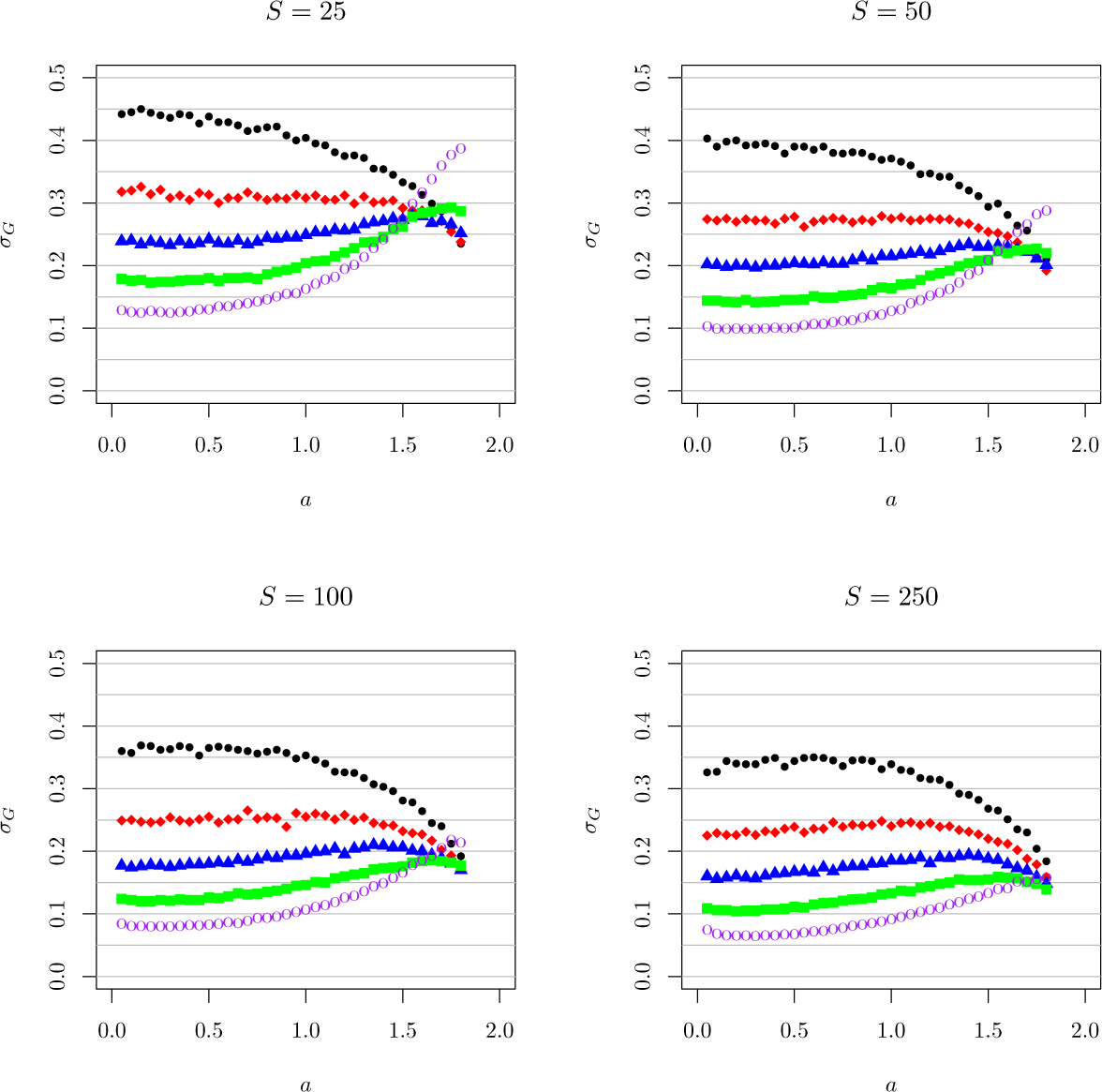
The same data as in Figure 22, but for fixed *S* and varying *n*. The colors/line shapes refer to the same numbers as in the previous figures, although they correspond to the value of *n* instead of *S*. *n* = 1000 is represented by purple open circles. Note that for small *S*, *σ* actually increases with increasing *n* for fixed *S*, when *a* ≳ 1.5, for the reasons noted in the main text.

**Figure 24:**
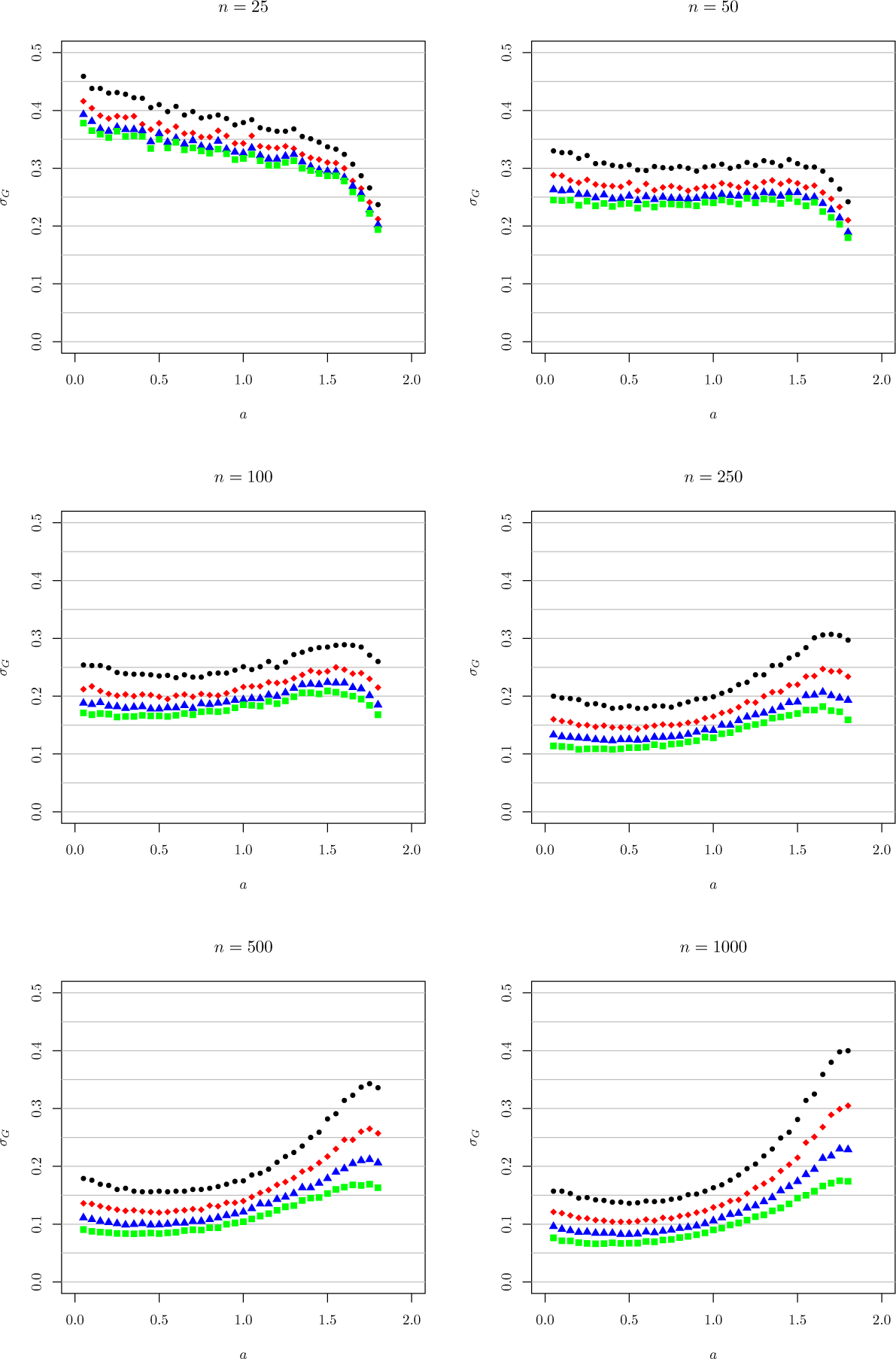
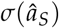, for data from 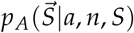 (Model A). Otherwise as in Figure 22.

**Figure 25:**
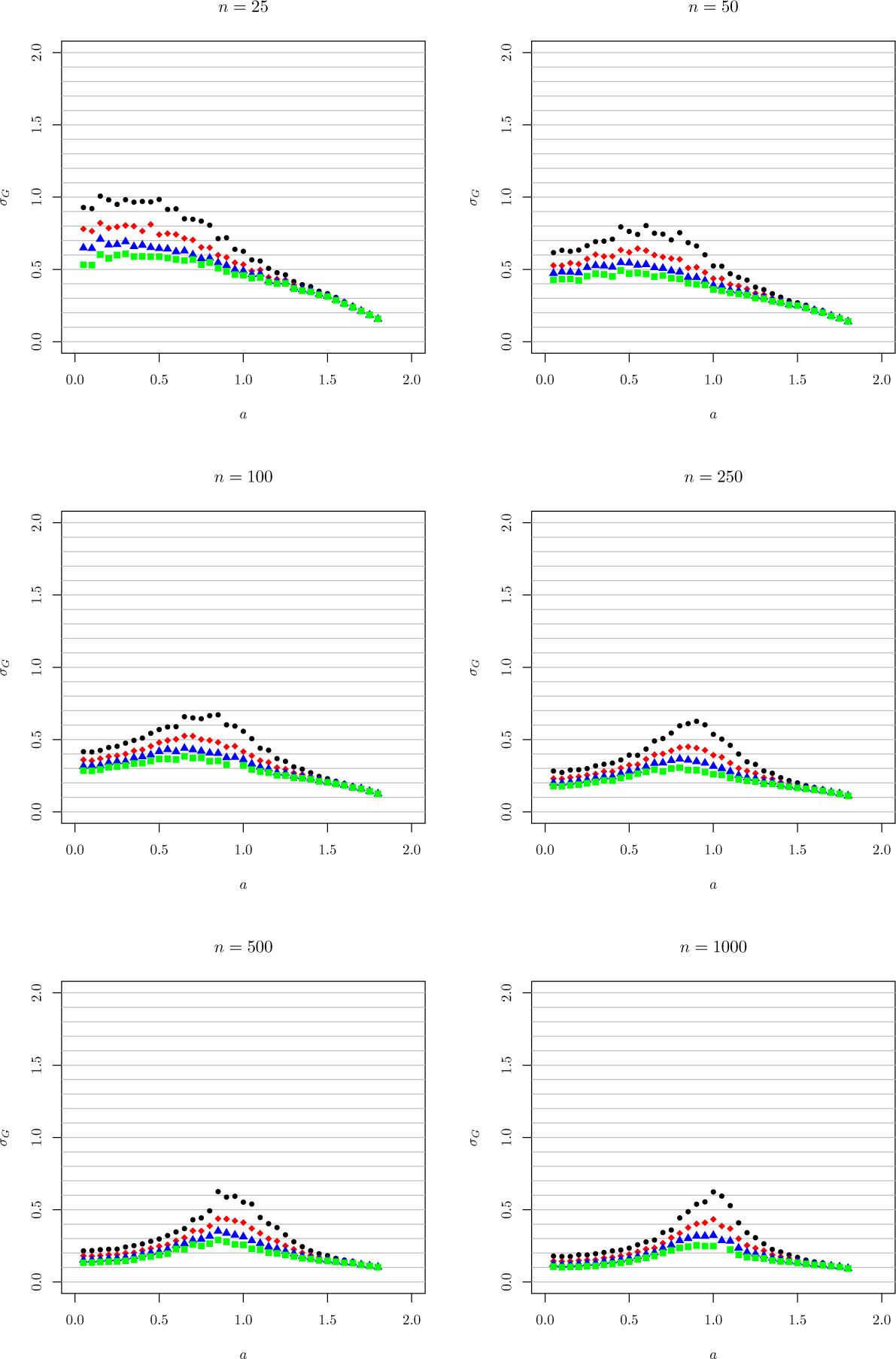
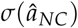, for data from 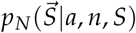 (Model N). Otherwise as in Figure 22.

The hump around *a* = 1 in Figure 25, for the non-singletons, is readily understood from inspection of the expected site-frequency spectra. When the singletons are removed, inference depends mainly on the slope of the SFS. In Figure 26 we show the slope of the expected and folded SFS. We see that as *a* increases, the slope falls as *a* approaches one, and then increases again as *a* approaches two. In the region where the slope turns around, it changes very slowly with *a*. As a result, the nonsingleton slopes are difficult to distinguish for a wide interval of values near *a* = 1.

The log-likelihood for Model NC, 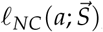, is generally bi-modal, with more or less symmetric peaks on either side of *a* = 1. The uncertainties plotted here do not account for that bimodality, because they were calculated only in a small neighborhood of the true value of *a*. In general, 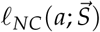 is unable to distinguish between the two peaks, although in many biological systems, it may be argued that only the peak with 0 < *a* < 1 is biologically relevant (Arnason and Halldórsdóttir, 2015, Eldon and Wakeley, 2006).

**Figure 26:**
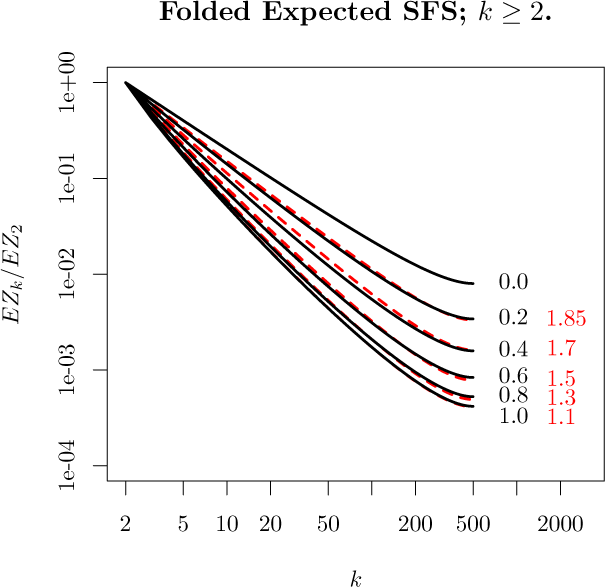
Expected folded site frequency spectrum for beta coalescent with *n* = 1000 and *a* ranging from zero to 1.85. The curves for *a* ≤ 1 are shown as solid black lines; the remaining curves as dashed red lines. The curves are normalized by *EZ*_2_. For computational reasons, *EZ_k_* ≡ *EL_k_*/*EL*_2_ is used as an approximation for *ER_k_*, as discussed above. The curve for *a* = 1.1 is mostly obscured by that for *a* = 1.

**Figure 27:**
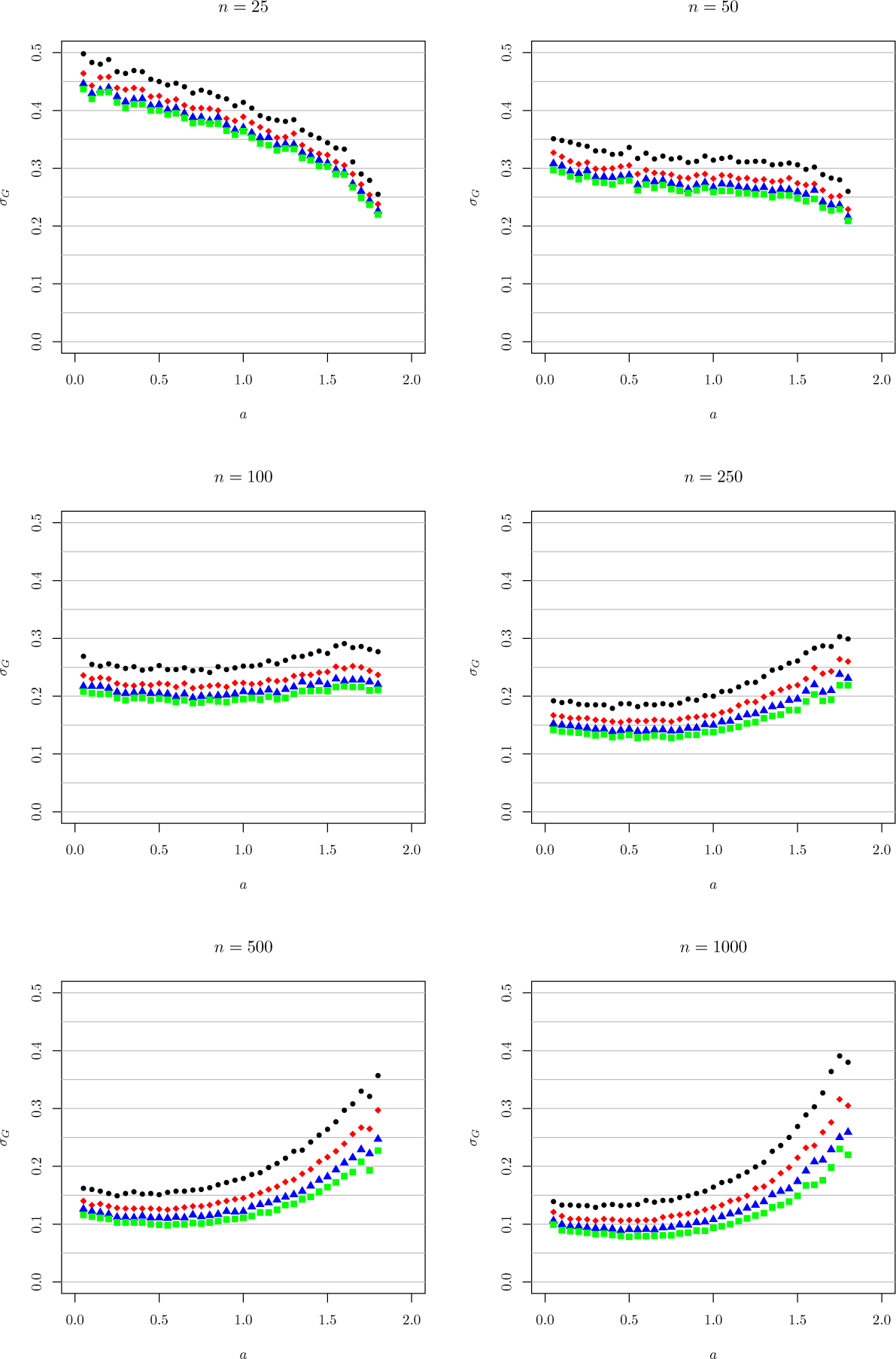
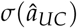, for data from 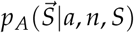 (Model A). Otherwise as in Figure 22.

**Figure 28:**
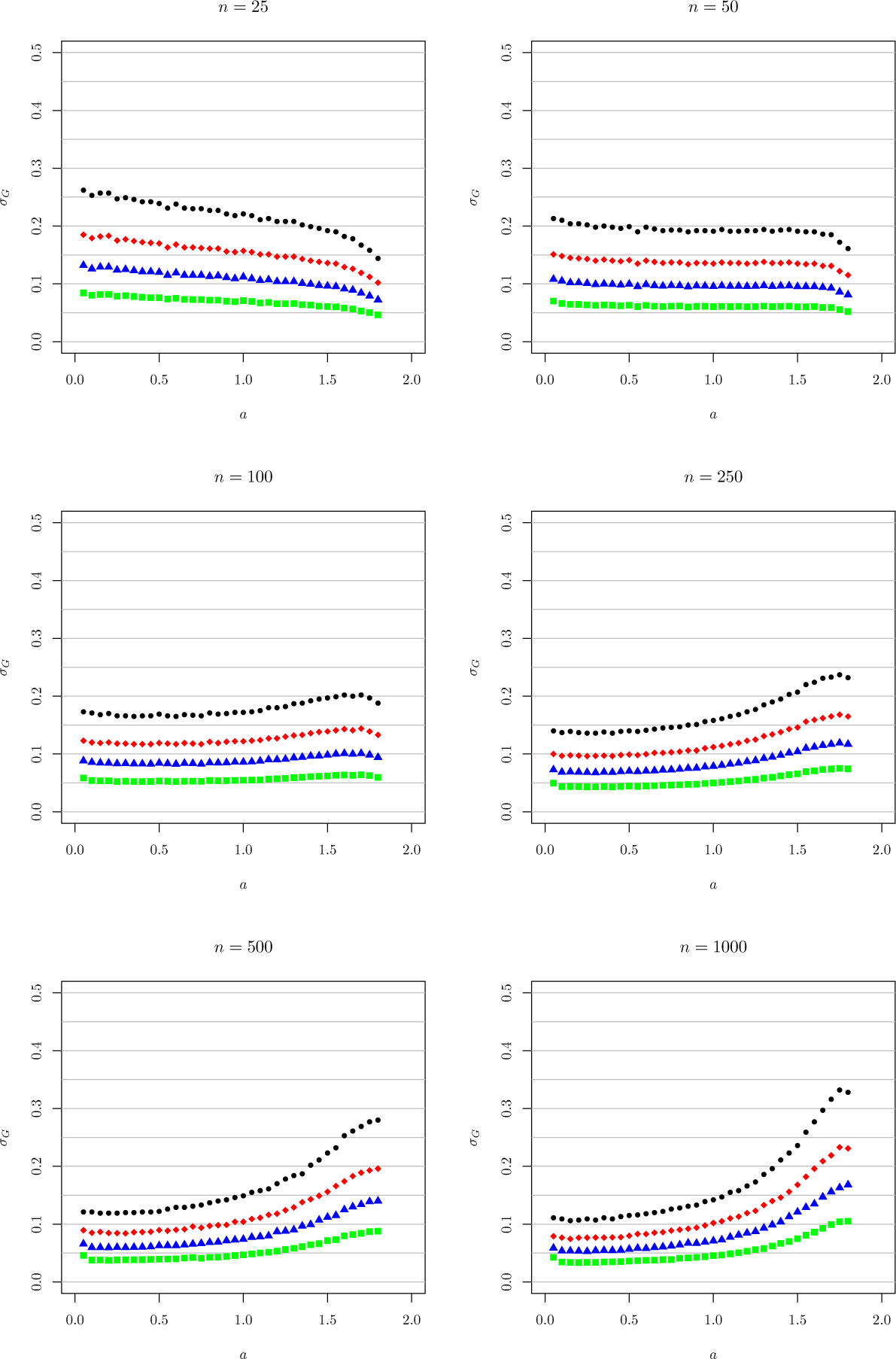
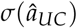, for data from 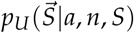 (Model U). Otherwise as in Figure 22.

### Eldon-Wakeley model

**Figure 29:**
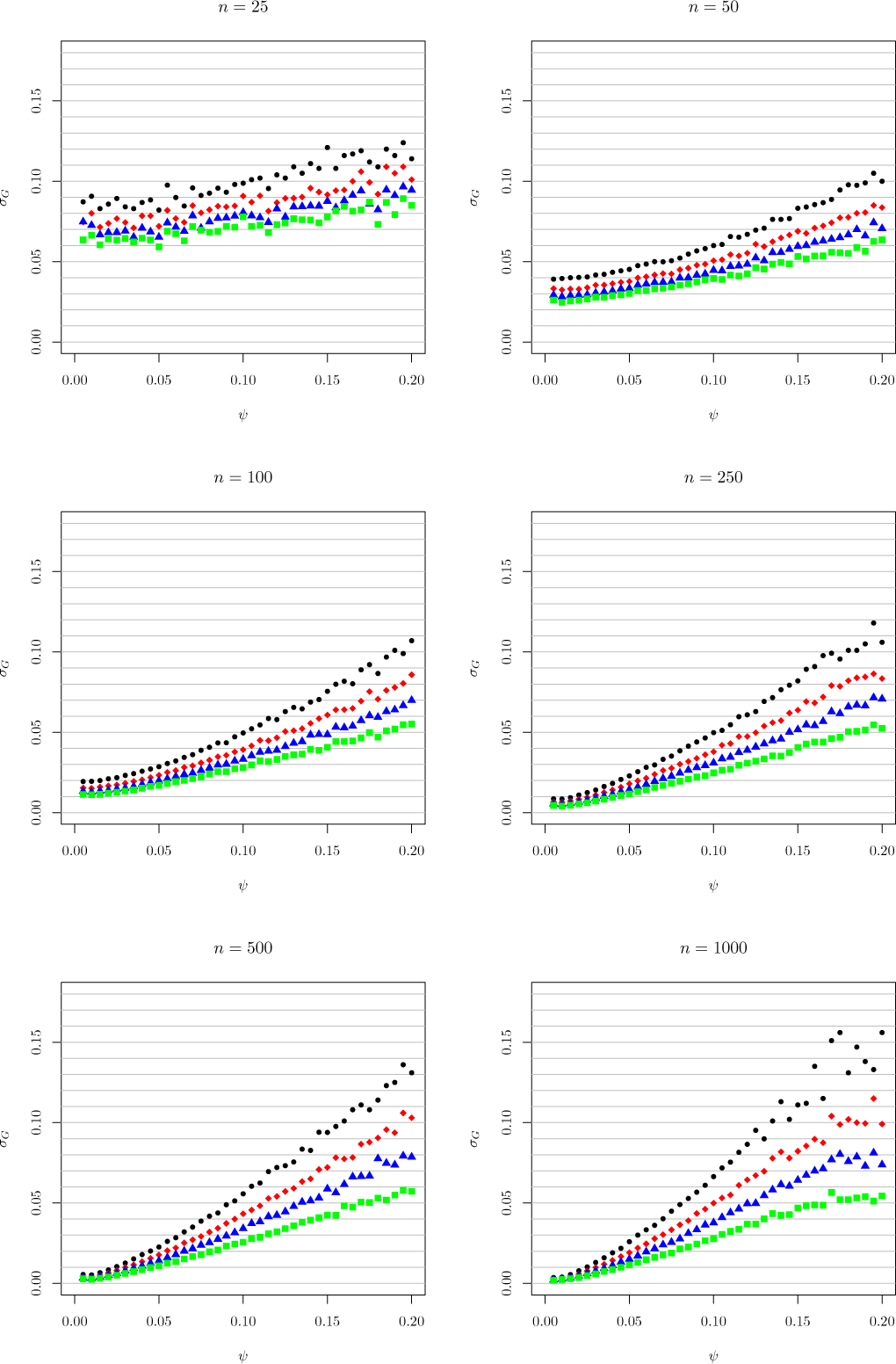
The asymptotic sampling standard deviation, 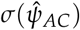, for data from 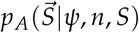 for the EW-coalescent. Otherwise as in Figure 22.

**Figure 30:**
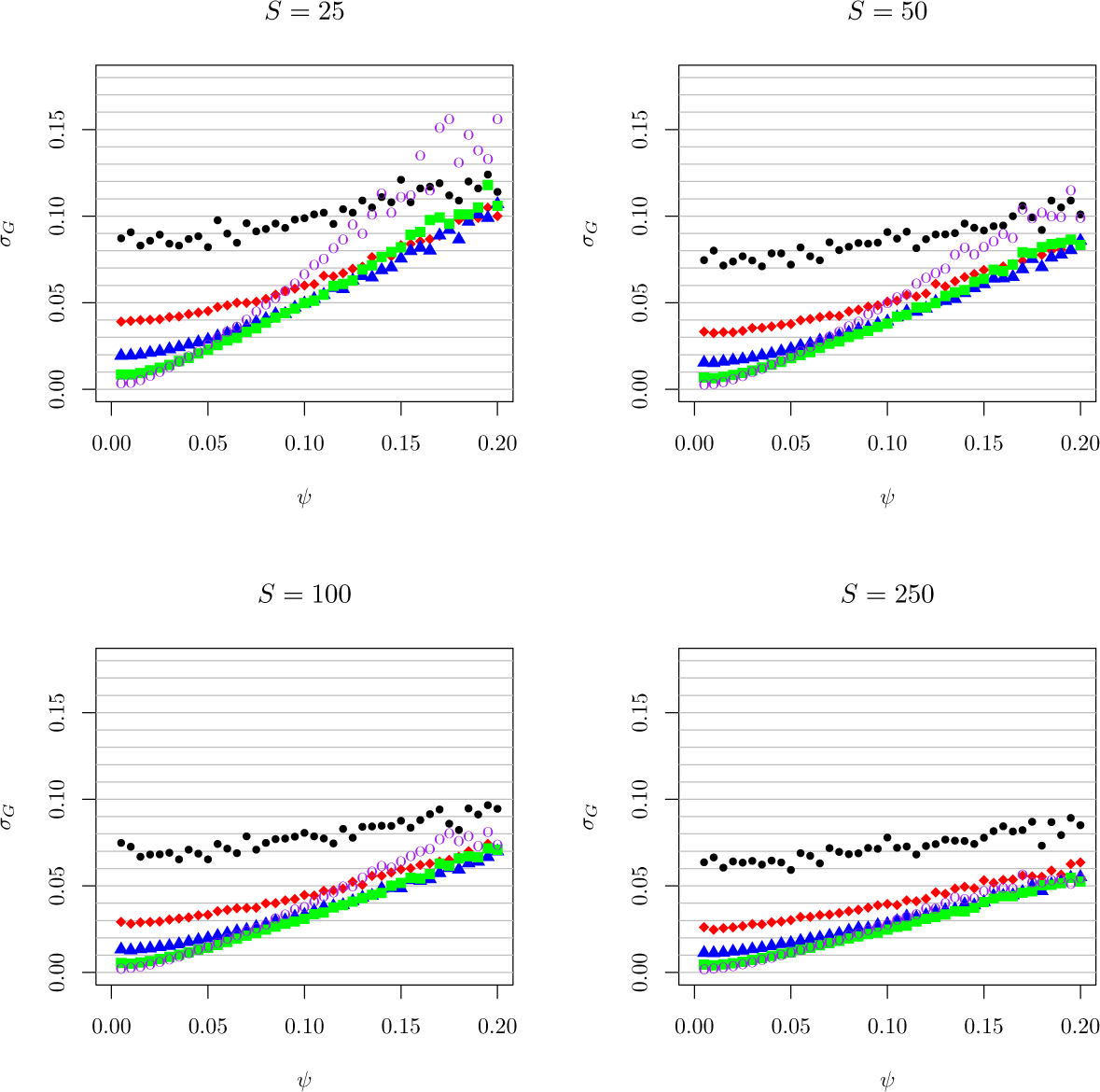
The same data as in Figure 29, plotted for fixed *S* with varying *n*. Otherwise as in Figure 22. Note that the uncertainties increase with *n* for values of *ψ* ≳ 0.05; a similar result was seen in Figure 23 for *β* ≳ 1.5, and was discussed in the Results section in the text.

**Figure 31:**
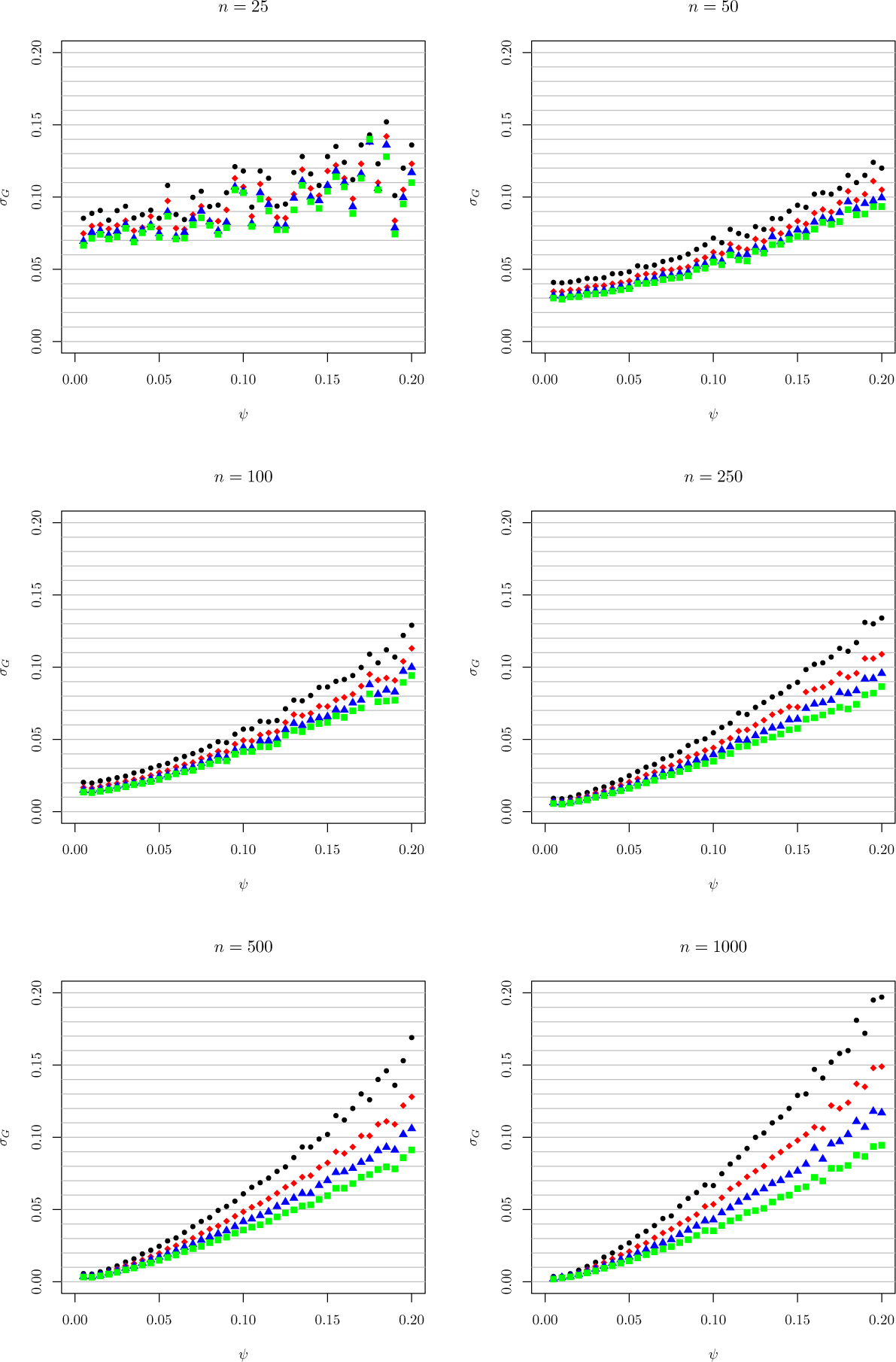
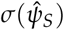, for data from for data from 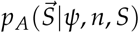. Otherwise as in Figure 29. Note that the data are very noisy for *n* = 25, an effect that is also seen in the plot of 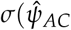.

**Figure 32:**
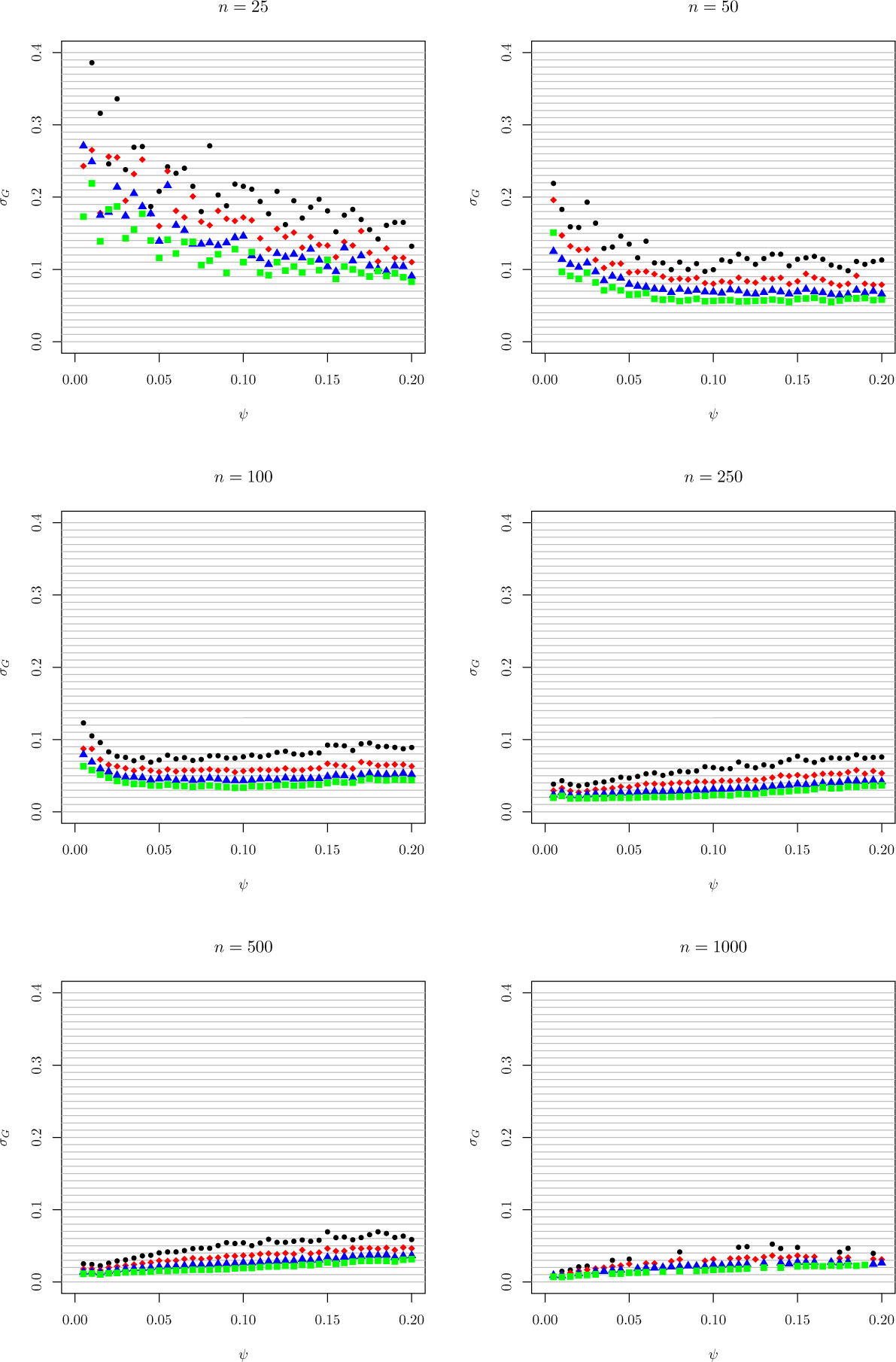
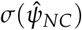, for data from 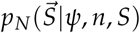. Otherwise as in Figure 29.

**Figure 33:**
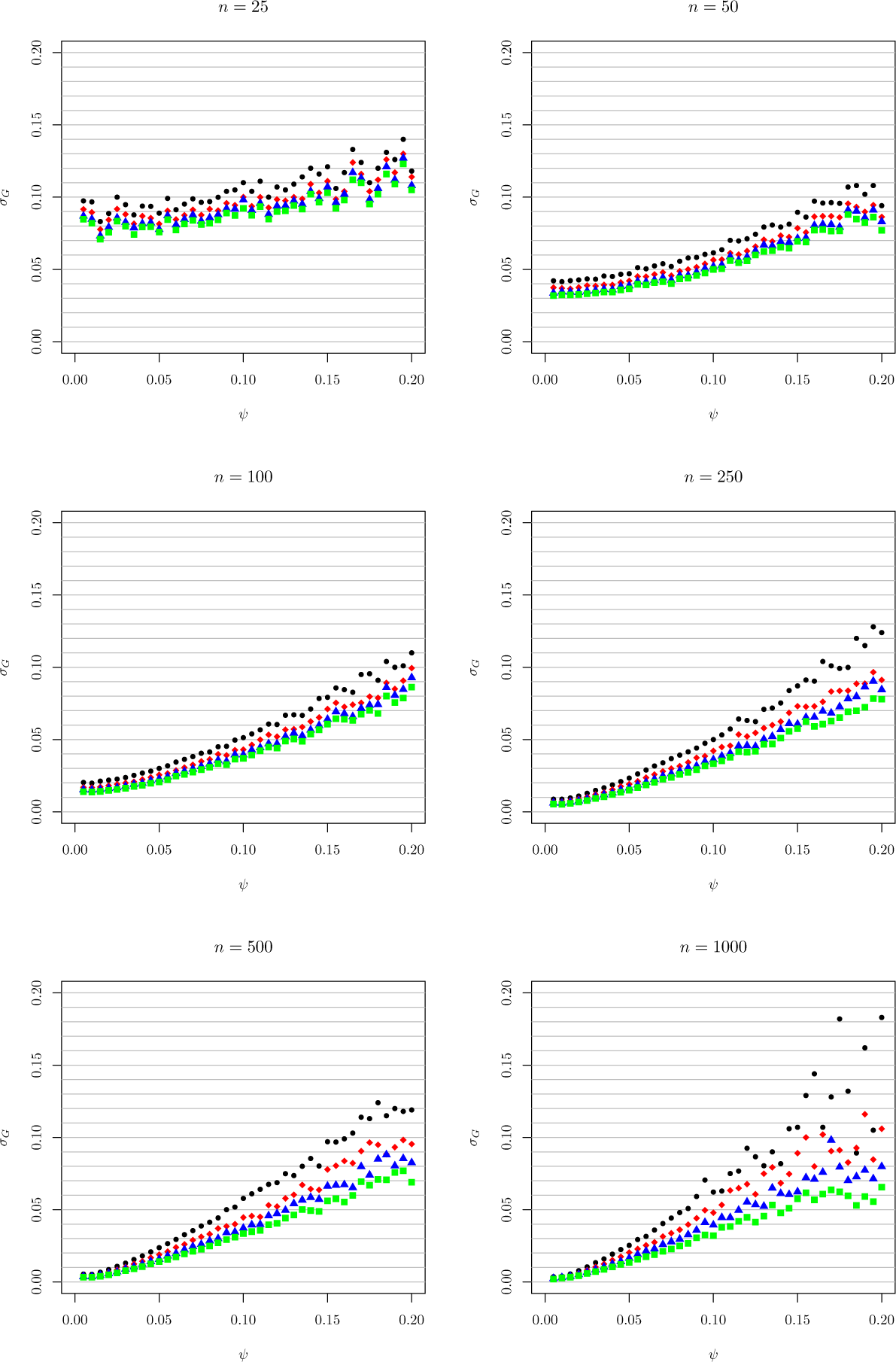
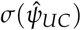, for data from 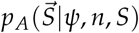. Otherwise as in Figure 29.

## Supplement C: Confidence intervals

### Beta model

**Figure 34:**
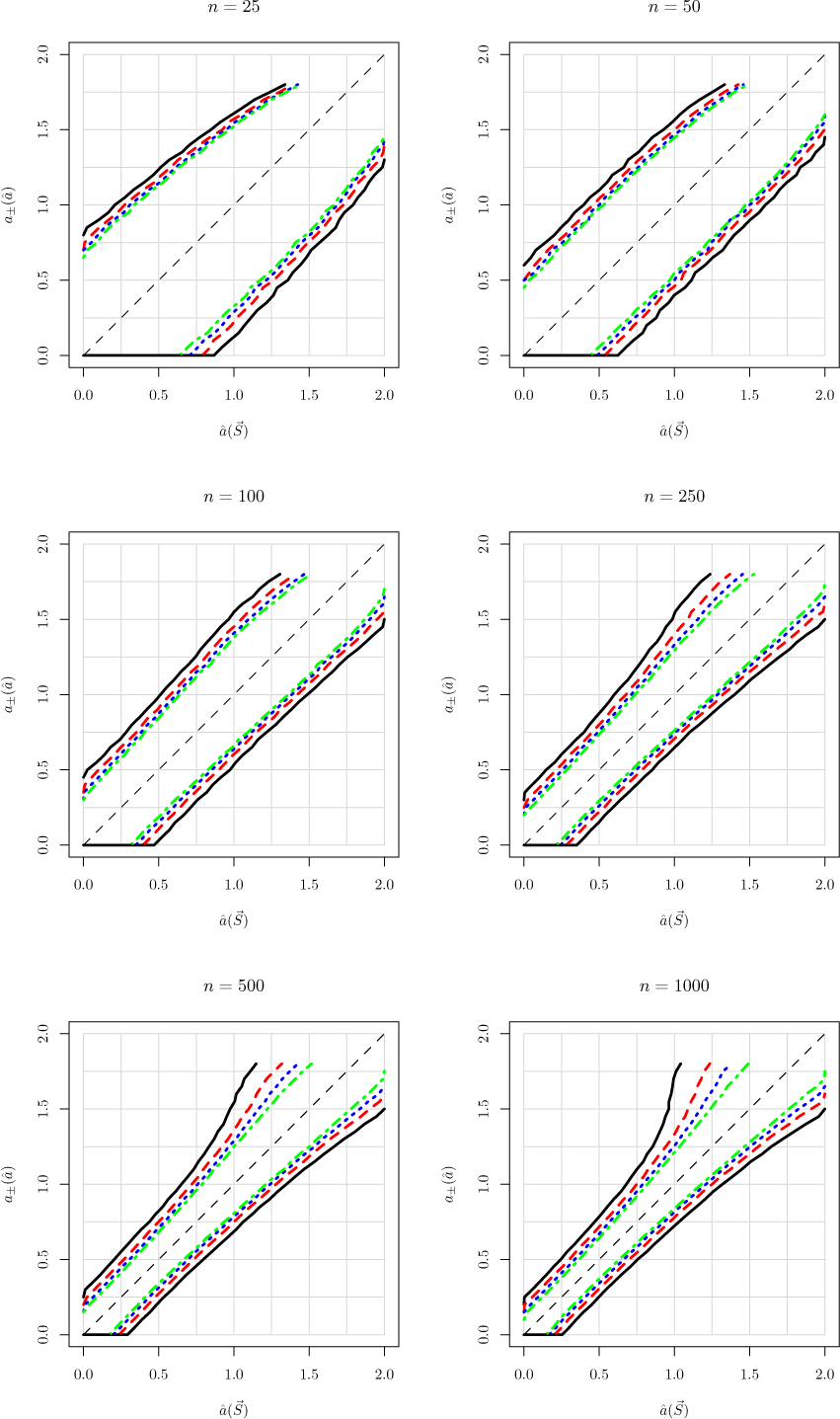
95% CI’s from 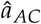, for data generated by Model A for the Beta coalescent, as a function of *n* and *S*. As in the text, the colors and line types are black/solid for *S* = 25, red/dashes for *S* = 50, blue/dots for *S* = 100, and green/dash-dots for *S* = 250. The CI for a sample 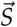, given *n* and *S*, is obtained by computing 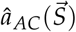, picking the graph corresponding to the value *n*, and the lines corresponding to *S*. The left and right endpoints of the CI are given by the intersection of the horizontal line at 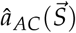 with the CI lines.

**Figure 35:**
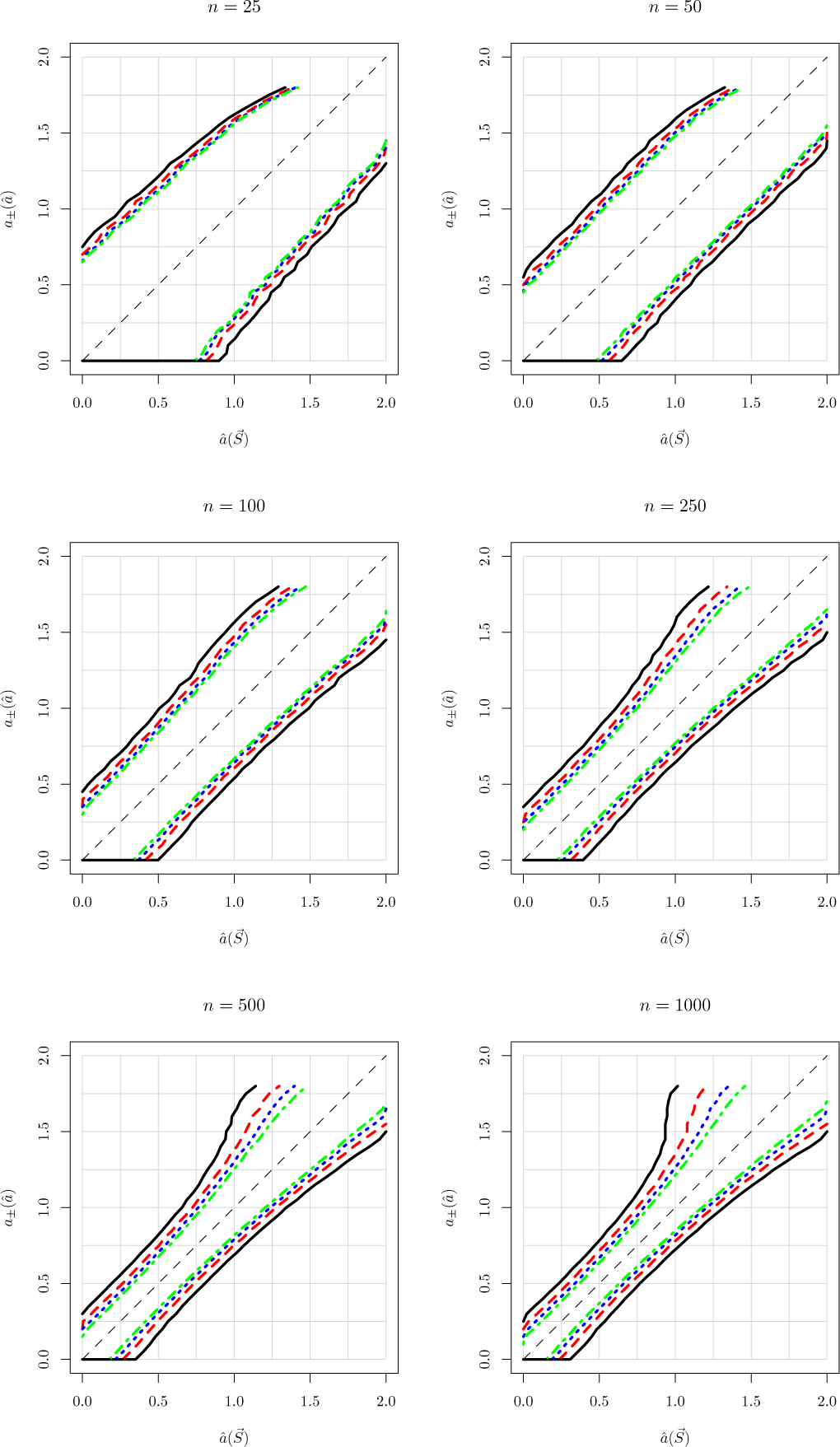
95% CI’s from 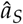, for data generated by Model A for the Beta coalescent, as a function of *n* and *S*. Otherwise as in Figure 34.

**Figure 36:**
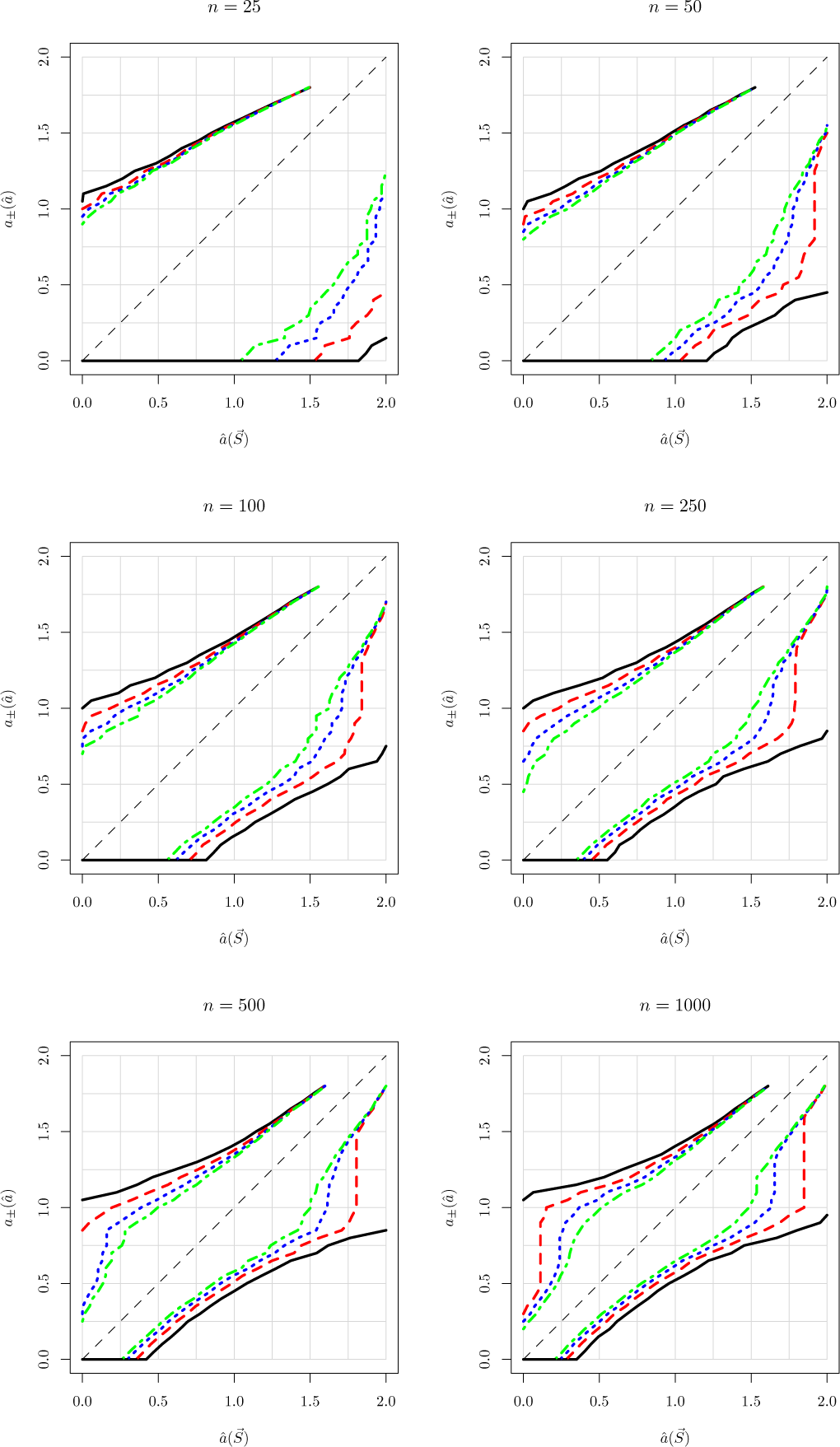
95% CI’s from 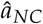, for data generated by Model N for the Beta coalescent, as a function of *n* and *S*. Otherwise as in Figure 34.

**Figure 37:**
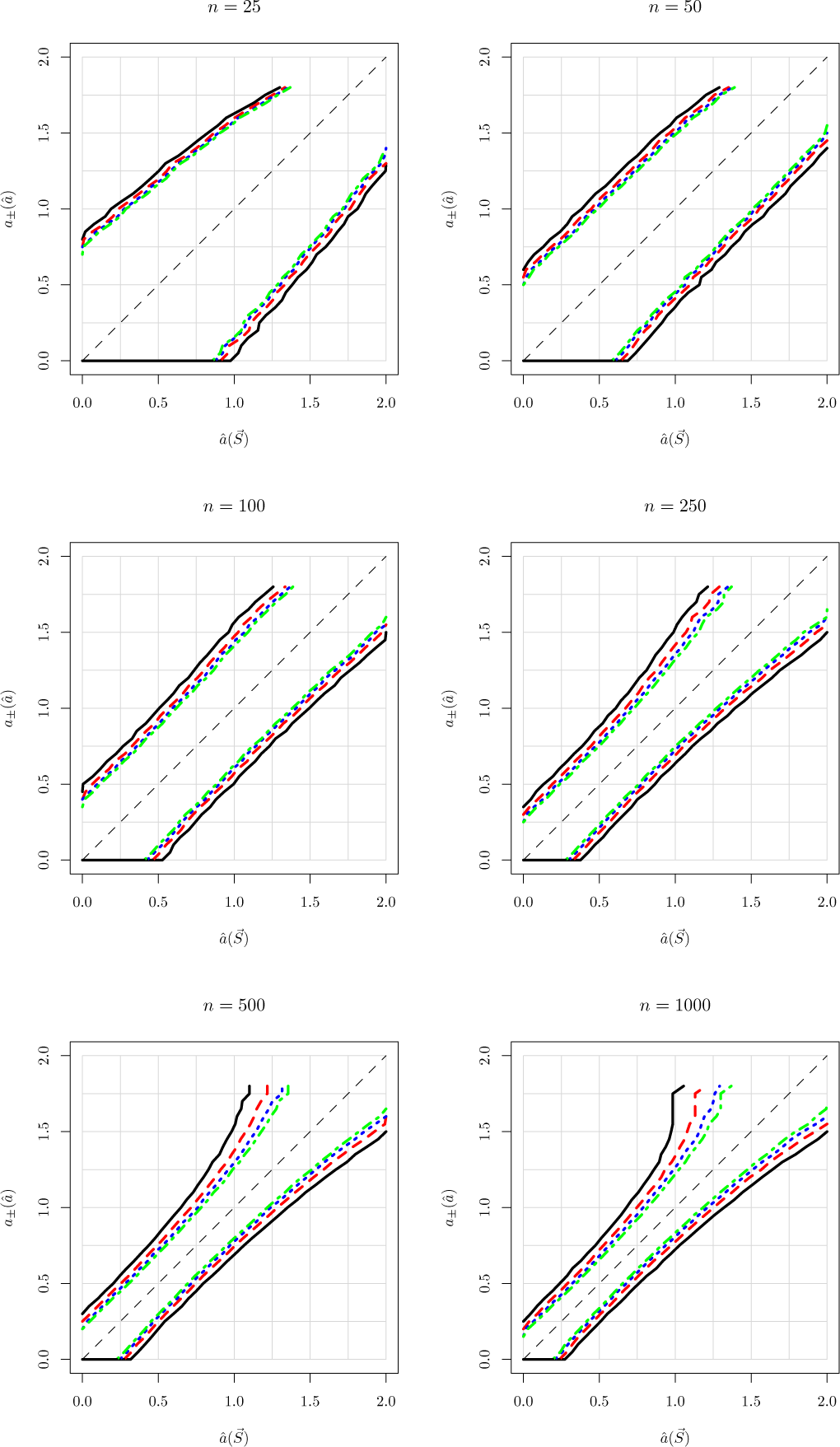
95% CI’s from 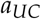, for data generated by Model U for the Beta coalescent, as a function of *n*, *S*, and *a*. Otherwise as in Figure 34.

### Eldon-Wakeley model

**Figure 38:**
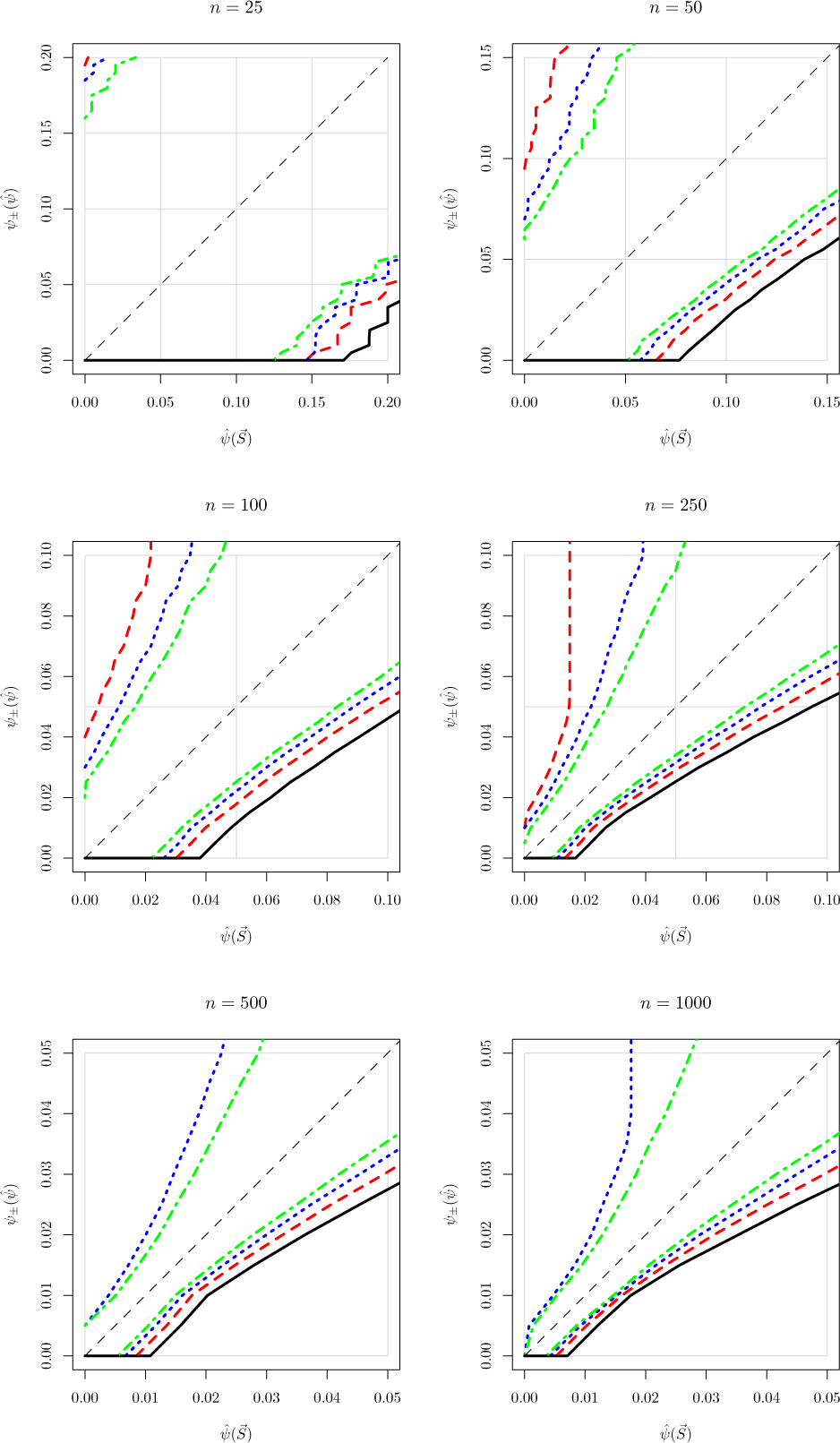
95% CI’s from 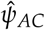 for data generated by Model A for the E-W coalescent, as a function of *n* and *S*. Otherwise as in Figure 34.

**Figure 39:**
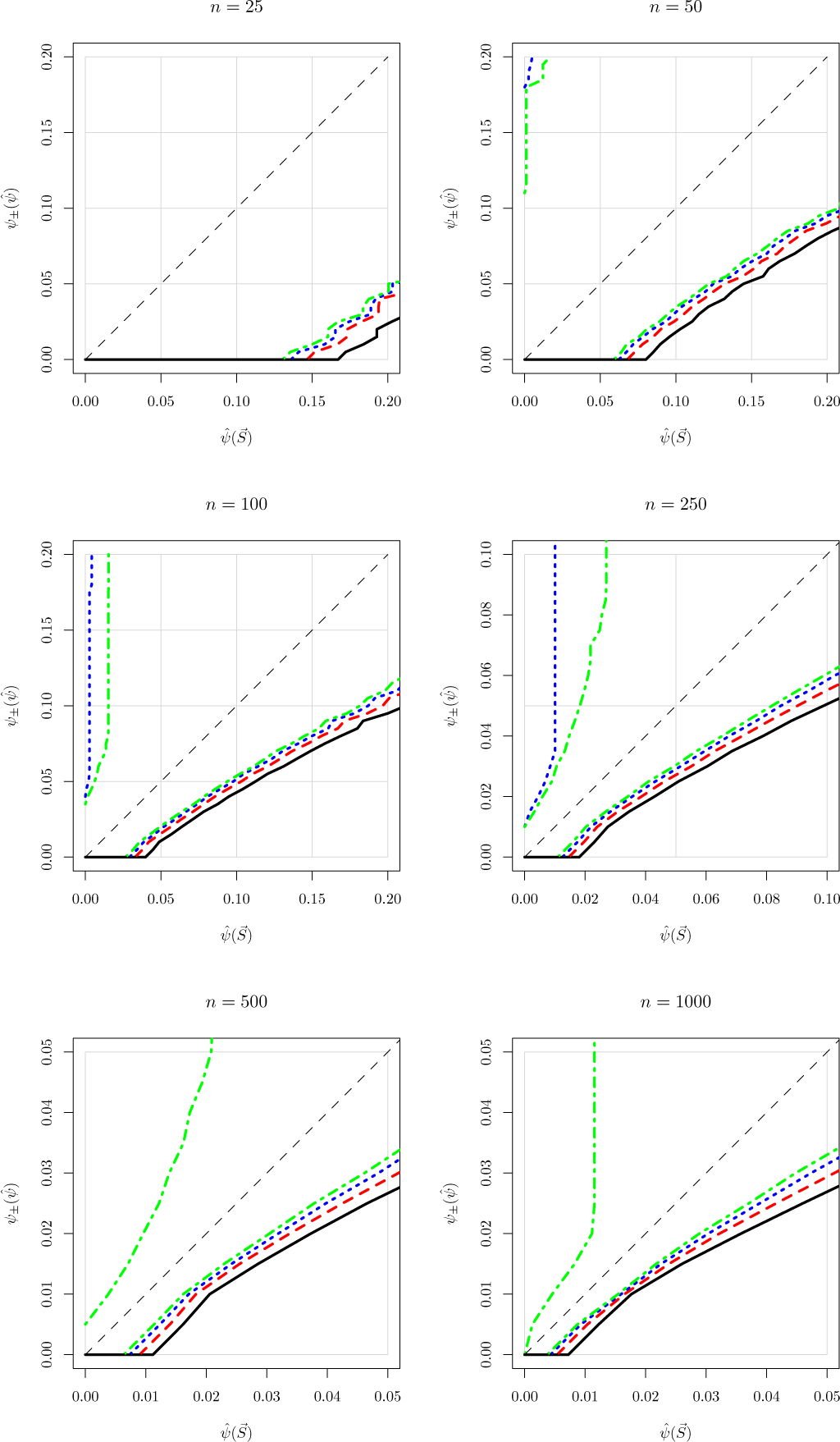
95% CI’s from 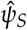 for data generated by Model A for the E-W coalescent, as a function of *n* and *S*. Otherwise as in Figure 38.

**Figure 40:**
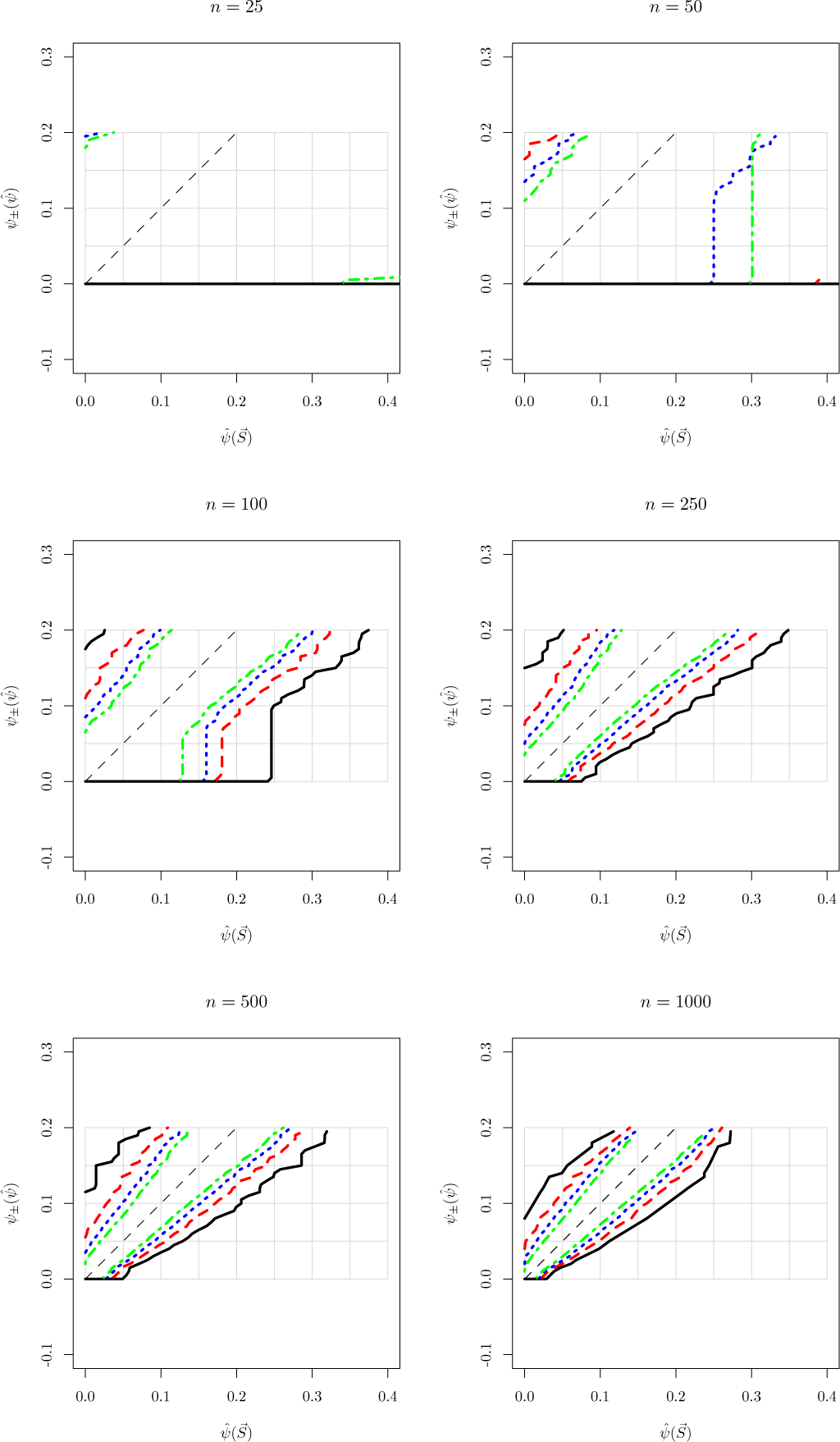
95% CI’s from 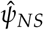 for data generated by Model N for the E-W coalescent, as a function of *n* and *S*. Otherwise as in Figure 38.

**Figure 41:**
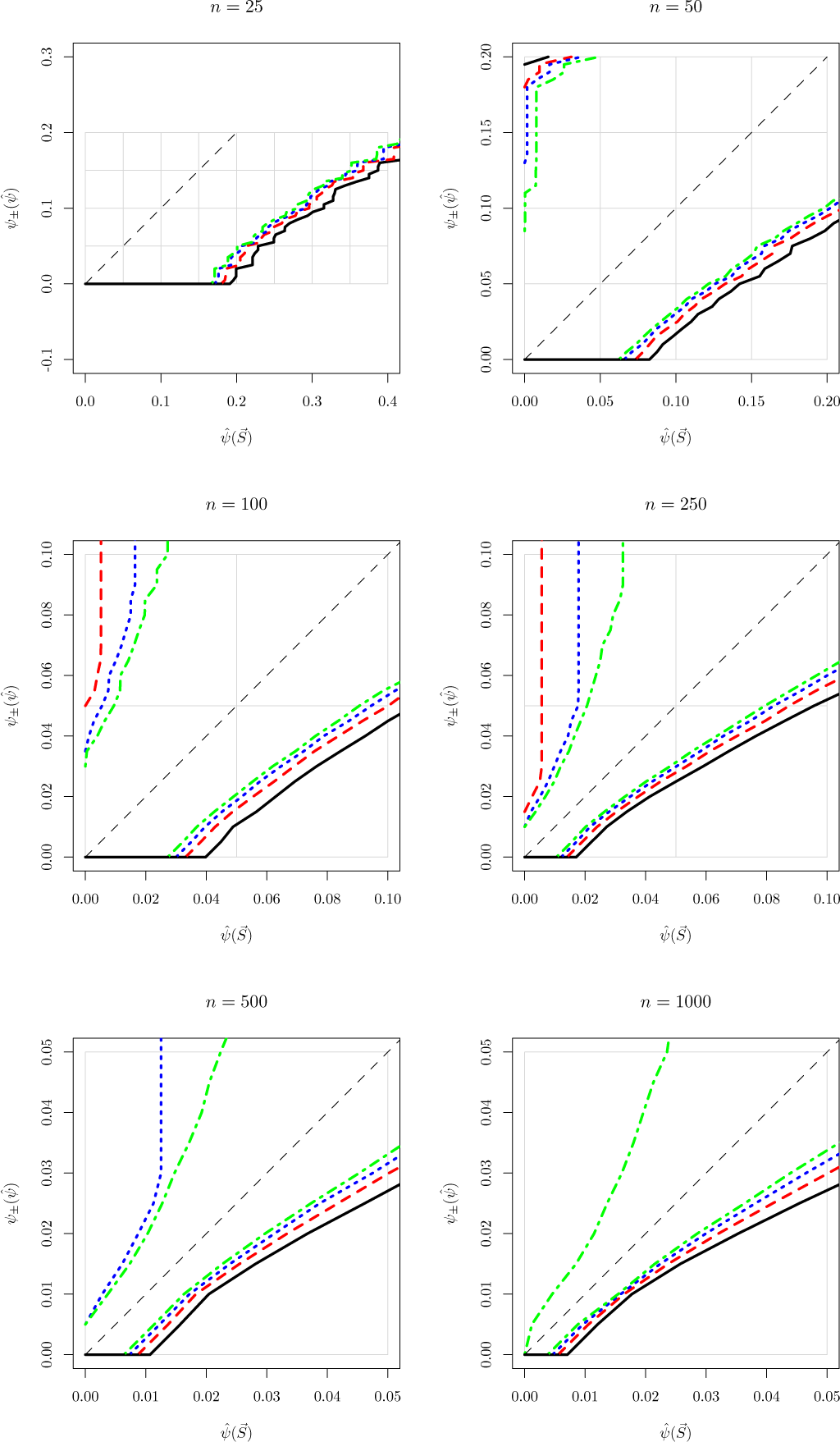
95% CI’s for 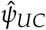 for data generated by Model A for the E-W coalescent, as a function of *n* and *S*. Otherwise as in Figure 38.

## Footnotes

1 In some papers, λ_*m*,*k*_ is used for the rate for a specific subset of *k* lineages. Our λ*_m_*_,*k*_ is 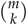 times the λ*_m_*_,*k*_ in these papers.

2 The interpretation is that a coalescent event involving a fraction of the population in the interval (*x*, *x* + *dx*) occurs at the rate *x*^−2^Λ(*dx*). The probability that *k* of *m* lineages will coalesce when the fraction is *x* is binom(*k*; *m*, *x*) (Eq. 7), and the total rate is given by the integral over the *x*-dependent rate.

3 Here, *θ* is a generic statistical parameter, which should not be confused with the scaled mutation rate, which does not appear in this paper.

4 The usual definition of efficiency applies only to unbiased estimators (Lehmann and Casella, 2006), and 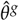 need not be unbiased, even if *g* is an unbiased estimating equation. Godambe (1960) generalized the Cramér-Rao theorem to the class of unbiased estimating functions, and in so doing, he also generalized the notion of efficiency to such functions, which includes the estimators in this paper.

5 In fact, it is slightly more efficient to not subtract the estimated mean, because the estimator then preserves all n degrees of freedom, instead of using up one degree of freedom on the estimation of the mean; see Zhang (1996).

